# A high-content imaging workflow to screen for molecules that reduce cellular uptake of α-synuclein preformed fibrils

**DOI:** 10.64898/2025.12.22.695981

**Authors:** Wolfgang E Reintsch, Andrea I Krahn, Chanshuai Han, Emmanuelle Nguyen-Renou, Carol X-Q Chen, Wen Luo, Irina Shlaifer, Tom A Pfeifer, Edward A Fon, Thomas M Durcan

## Abstract

A classical pathological hallmark of many neurodegenerative diseases is the formation of protein-rich aggregates and inclusions. In Parkinson’s disease (PD), α-synuclein (α-syn) constitutes a major protein component of pathological inclusions, termed Lewy bodies. These α-syn aggregates are hypothesized to spread throughout the nervous system by cell-to-cell transmission acting as templates to amplify aggregate formation. In vitro generated α-syn aggregates, commonly called preformed fibrils (PFFs), have been used to investigate a number of aspects related to α-syn mediated pathology across different model systems. Here we describe a semi-automated assay to screen for small molecules that interfere with the cellular uptake and accumulation of PFFs. The assay uses dopaminergic progenitor cells (DPCs), derived from human induced pluripotent stem cells (hiPSCs). In an initial screen, we tested 1520 small molecules and identified several molecules that strongly reduce intracellular PFF load in DPCs. From these hits, candidate compounds were validated in dopaminergic neurons (DNs) to demonstrate the utility of the assay. This assay provides a robust, scalable and adaptable tool to screen for molecules that affect PFF uptake in hiPSC-derived cell models. Within the scope of this screen, it led to the identification of a set of compounds with diverse annotated targets that effectively reduce the uptake of synuclein aggregates in DPCs and DNs.

## 1. Introduction

One of the classical pathological hallmarks of Parkinson’s disease (PD) is the accumulation of intraneural inclusions, termed Lewy bodies (LBs) [1, 2]. Misfolded and fibrillar α-synuclein (α-syn) protein is a major protein component of LBs and it is thought to be implicated in their formation. LBs have also been detected in other synucleinopathies including multiple systems atrophy (MSA) and Lewy body dementia (LBD) [3–5]. Recent studies have shown that LBs are structurally complex and contain a range of other proteins as well as various membranous structures, including vesicles and dysmorphic organelles. [6, 7].

α-syn is an abundant neuronal protein and in its soluble, native form is implicated in vesicular dynamics and maintenance of pre-synaptic terminals [8–10]. Evidence that α-syn accumulation is causative in PD and other synucleinopathies came from observations that mutations that lead to increased expression or misfolding of α-syn lead to familial, early onset forms of PD [11–16]. Several studies confirmed that high levels of α-syn and the presence of α-syn aggregates contribute to cellular stress and toxicity across a variety of experimental systems [17–19].

Autopsy studies have led to a model whereby LBs progressively spread through the peripheral and central nervous system [20, 21]. Aggregated α-syn is thought to mediate this progression in a prion-like fashion [22, 23]. In this model, misfolded α-syn forms oligomers and higher order aggregates by recruiting monomeric, native α-syn. While this contributes to the formation of LBs, some aggregates are transferred to neighboring cells, acting as seeds to induce further aggregation [24]. Based on this model, preventing α-syn aggregate transmission and seeding represents a potential therapeutic target in PD.

The use of *in vitro* generated α-syn aggregates has been instrumental for studying the cellular mechanisms of α-syn aggregate uptake, intracellular deposition, degradation, cell to cell transfer and seeding [25–27]. Model systems to investigate these processes range from immortalized cell lines [28, 29], primary cultures [30] and human induced pluripotent stem cell (hiPSC)-derived cell types [29, 31] to mouse [32] and marmoset animal models [33]. Standardized protocols have been developed to generate and validate these aggregates, commonly referred to as preformed fibrils (PFFs) [28, 34].

In recent years, hiPSC-derived cell lineages have become increasingly important as *in vitro* models. hiPSCs can be generated from blood, skin or even urine of any individual, amplified and banked in theoretically unlimited numbers. Potentially, they can be directed into any cell type of interest found within the human brain and over the last decade different protocols have been developed to differentiate hiPSCs into a variety of neuronal cell types. However, the procedures to achieve fully differentiated neurons and glia cells are often complex and time consuming and it can be challenging to produce sufficient cell numbers for high throughput applications [35, 36]. Here we describe a simplified approach, amenable to high content screening, using hiPSC-derived dopaminergic precursor cells (DPCs). Combined with fluorescently labeled PFFs, this approach enabled us to create a comparatively fast, robust and scalable screening strategy to test a large number of small molecules for their ability to reduce cellular PFF uptake. As a proof of concept, we screened a library of 1520 FDA and/or EMA approved small molecules and identified several compounds that reduce the accumulation of intracellular PFFs by more than 50% without affecting cell viability. We confirmed the inhibitory effect of the active compounds in differentiated dopaminergic neurons (DNs), showing that these compounds are equally effective in more mature neuronal cells. Active compounds were found to belong to different classes, including for example lysosomotropic molecules, cardiac glycosides and tyrosine kinase inhibitors. From our findings, this automated screen provides a workflow towards assessing small molecules and biologics for an effect on uptake of α-syn aggregates in neurons, and potentially other cell types.

## 2. Materials and Methods

### 2.1. Cell culture

hiPSCs from the healthy control cell line AIW002-02 [37] were used to generate the neuronal cells in this research. The cell line was provided by the Clinical Biospecimen Imaging and Genetic Repository ([C-BIG](https://www.mcgill.ca/neuro/research/open-biobank)) at the Montreal Neurological Institute (C-BIG ID: [IPSC0063](https://hpscreg.eu/cell-line/CBIGi001-A)). Their use was approved by the McGill Research Ethics Board (IRB Study Number A03-M19-22A / eRAP 22-03-027). Cells were maintained in a 37 °C incubator with 5% CO2.

### 2.2 Dopaminergic progenitor cell induction and dopaminergic neuron differentiation

Induction of ventral midbrain DPCs was based on a protocol previously described [38, 39], using a combination of small molecules over three weeks. hiPSCs were cultured on Matrigel (Corning Millipore, 354277)-coated plates and daily mTeSR1 medium (STEMCELL Technologies, 85850) changes until cells reached ∼80% confluence (usually 5-7 days after plating). Next, cells were dissociated with Gentle Cell Dissociation Reagent (GCDR) (STEMCELL Technologies, 07174) and cultured on Matrigel-coated flasks for one week in neural induction medium (DMEM/F12 (ThermoFisher Scientific, 10565042) supplemented with N-2 (ThermoFisher Scientific, 17502048), B-27 (ThermoFisher Scientific, 17504044), MEM non-essential amino acids (NEAA) (ThermoFisher Scientific, 11140050), 1 mg/mL bovine serum albumin (BSA) (ThermoFisher Scientific, 15260037), 200 ng/mL Noggin (PeproTech, 120-10C), 200 ng/mL SHH (GenScript, Z03067), 3 µM CHIR-99021 (Selleckchem, S7146), 10 µM SB431542 (Selleckchem, S1067), and 100 ng/mL FGF-8 (PeproTech, 100-25) with daily medium changes. Cells were dissociated with GCDR and passaged on Matrigel-coated flasks on day 7 and 14 and cryopreserved on day 21 in fetal bovine serum (ThermoFisher Scientific, 12484028) containing 10% DMSO (Fisher, BP231-1). For experimentation, frozen DPC aliquots were defrosted and cultured for one week in flasks that were coated with 1 μg/mL poly-L-ornithine (PLO) (Sigma, P3655) and 5 μg/mL laminin (Sigma, L2020), and maintained in STEMdiff Neural Progenitor Basal Medium (STEMCELL Technologies, 05834) supplemented with STEMdiff Neural Progenitor Supplement A (STEMCELL Technologies, 05836), STEMdiff Neural Progenitor Supplement B (STEMCELL Technologies, 05837) and 2 μM Purmorphamine (Sigma, SML-0868) with medium changes every two days. Following this one-week expansion, cells were dissociated with StemPro Accutase Cell Dissociation Reagent (ThermoFisher Scientific, A1110501) into a single-cell suspension and seeded in PLO/laminin-coated black 96 well microplates (Corning, 353219) at a density of 15,000 cells per well in dopaminergic neural differentiation medium (Brainphys Neuronal medium (STEMCELL Technologies, 05790) supplemented with N2A Supplement A (STEMCELL Technologies, 07152), NeuroCult SM1 Neuronal Supplement (STEMCELL Technologies, 05711), 20 ng/mL BDNF (PeproTech, 450-02), 20 ng/mL GDNF (PeproTech, 450-10), 0.1 mM compound E (STEMCELL Technologies, 73954), 0.5 mM db-cAMP (Biosynth, ND07996), 0.2 mM ascorbic acid (Sigma Aldrich, A5960) and 1 mg/mL laminin).

### 2.3 Immunofluorescence

DPCs were fixed for 15 min with 4% formaldehyde (ThermoFisher Scientific, 28908) in PBS, permeabilized for 5 min with 0.2% Triton X-100 (Sigma-Aldrich, X-100) in PBS, then blocked for one hour in 2% BSA and 0.02% Triton X-100 in PBS. Primary antibodies were diluted in blocking buffer and added for overnight incubation at 4°C. The cells were then washed three times with 0.2% Triton X-100 in PBS and secondary antibodies diluted in blocking buffer were added for one hour in the dark. Nuclei were stained with 2.5 μg/mL Hoechst33342 (ThermoFisher Scientific, H3570) in PBS for 10 min, then cells were washed three times with 0.2% Triton X-100 in PBS before imaging on a CellInsight CX5 HCS Platform (ThermoFisher Scientific Scientific) and analysis with the HCS Studio 5.0 Cell Analysis Software (ThermoFisher Scientific).

### 2.4 Quantification of mRNA expression markers by quantitative real-time PCR analysis

RNA was isolated using a NucleoSpin RNA kit (Takara, 740955) according to the manufacturer’s instructions. cDNA was synthesized using the iScript Reverse Transcription Supermix (BioRad, 1708840). qPCR was performed on the QuantStudio 5 Real-Time PCR System (Applied Biosystems) using the primers listed below.

### 2.5 Production of labelled α-synuclein preformed fibrils

Production and labelling of recombinant PFFs followed a previously described protocol [40]. Briefly, glutathione S-transferase (GST)–tagged α-syn was expressed and isolated from E. *coli*. After removal of the GST-tag and purification, monomeric α-syn was concentrated to 5 mg/mL and shaken for 5 days to induce fibril formation. The formed α-syn fibrils were then sonicated to produce final PFFs with an average length of <100 nm. The length and shape of PFFs was assessed by electron microscopy and dynamic light scattering [41]. For the screen, the same batch of PFF was used throughout. The PFFs were fluorescently labelled with Alexa Fluor 488 (Molecular Probes, A20000) or Alexa Fluor 633 (Molecular Probes, A20005) using a Slide-A-Lyzer MINI dialysis device (ThermoFisher Scientific, 88400). Labeled PFFs were stored in 50 ul aliquots at −80°C.

### 2.6 Uptake of α-synuclein preformed fibrils in dopaminergic progenitor cells

DPCs were seeded in PLO/laminin-coated black 384 well microplates at a density of 4,000 cells per well in DMEM/F12 (ThermoFisher Scientific, 10565042) supplemented with antibiotics-antimycotic (ThermoFisher Scientific, 15240-062), N-2, B-27, and NEAA, and allowed to attach for 24 hours. Varying concentrations of fluorophore-labeled PFF (PFF-Alexa488 or PFF-Alexa633) was added to the wells at different time points. Cells were washed twice with PBS and fixed for 10 min with 2% formaldehyde in PBS and then counterstained with 2.5 μg/mL Hoechst33342 in PBS for 10 min. To validate the PBS wash efficiency, live cells were washed with PBS or treated with Trypan Blue (ThermoFisher Scientific, 15250061), counterstained with Hoechst and imaged immediately. Images were acquired on a high content imaging system, using a X10 objective (NA 0.45) (CellInsight CX7, ThermoFisher Scientific or ImageXpress Micro Confocal, Molecular Devices) and analysed for cytoplasmic PFF content [42], using the software included in either system. Nuclei were identified in the first channel, and the obtained mask was expanded. Using a threshold value in the PFF channel slightly above background, the total intensity of PFFs within the expanded mask was obtained. The average intensity of the nuclear Hoechst33342 stain and the area of the nucleus were also determined. Per cell values were averaged per well and normalized against the mean of the combined vehicle controls in each plate separately.

For PFF uptake inhibition experiments, Heparin (Sigma, H3149), Rottlerin (Sigma, 557370) and Tilorone (Sigma, 220957) were manually added to wells at different concentrations 1 hour before PFF-Alexa488 addition. After 24 hours, microplates were processed, imaged and analysed as described above. Statistical significance was tested using one-way ANOVA and Dunnett’s corrected multiple comparisons test in GraphPad Prism (GraphPad Software, LLC).

### 2.7 Prestwick Chemical Library Screen

The Prestwick Chemical Library® (Prestwick Chemical) was transferred from 96 well plates into five 384 well plates with 304 molecules per plate as well as Rottlerin and Tilorone at three different concentrations as positive controls and DMSO-only as the vehicle (negative) control. The screen was automated using an EL406 Washer Dispenser (BioTek) for plate coating, cell seeding, PFF addition, fixing, and staining as previously described in detail [43]. After DPC seeding and incubation for 24 hours, compounds were added at a final concentration of 8 µM using a 384 well pinning tool (VP Scientific Inc.) installed on a plate handler (Matrix PlateMate Plus, Thermo Fisher Scientific) and incubated for one hour before the addition of 80 nM PFF-Alexa488. After 24 hours, cells were fixed, stained, imaged, and analysed as described in **section 2.6**. Normalization against the DMSO-only controls was performed separately for each plate first and then the data was combined into a full set.

To confirm active compounds, dilution series of selected compound were prepared in a 384 well plate and transferred to test plates with the pinning tool (12, 4, 1.3 and 0.4 μM final concentration).

Data normalization and Z’ values calculation was performed in Excel (Microsoft). Z-prime values were calculated according to [44].

### 2.8 Active compound confirmation in dopaminergic neurons

DPCs were cultured and differentiated to DNs in 96 well plates as described in **section 2.2**. The cells were seeded at a lower concentration (10,000 cells/well) to reduce cell aggregation and allow for clear identification of single cell bodies. After 2 weeks of differentiation, candidate compounds and control compounds were added at 4 different concentrations, using a non-contact liquid dispenser (I-DOT, Dispendix). After 1 hour incubation, PFF treatment and plate processing were performed as described in **section 2.6**. The plates were then imaged using an Opera Phenix+ High Content System (Revvity) using a X20 water immersion objective (NA 1.1), acquiring 25 fields per well. Image analysis was performed with the Opera software (Harmony, Revvity). The principal analysis steps were similar to the steps described for PFF uptake analysis in DPCs with the following adaptations to allow proper detection of PFF in DNs: To exclude dead cells and other, non-neuronal cells from the analysis, the cell population was filtered first by nuclear size and Hoechst staining intensity. The exclusion parameters were determined by comparing nuclear sizes and intensities in neuronal and dead/ non-neuronal cells. Dead cells were identified as having smaller, brighter nuclei, non-neuronal cells as having bigger, less bright nuclei. Additionally acquired bright field images were used to visually confirm the relationship between nuclear parameters and cell type. In addition, since we noted that non-neuronal cells, compared to DNs, incorporated very high amounts of PFF, DN cell bodies that overlapped with these cells and therefore showed falsely attributed very high PFF intensity values, were also excluded. The per cell total PFF intensity values of the pure DN population were then aggregated to calculate the mean per well and normalized to the mean PFF intensity of the DMSO-only controls in the same plate.

## 3. Results

### 3.1. Characterization of dopaminergic neuronal precursor cells

As part of the proposed workflow, access to large numbers of DPCs is essential. For the cells used in these studies, an extensive quality control process was followed to ensure their midbrain, dopaminergic precursor lineage. As a first step in this process, bright field images of DPCs were obtained to monitor for growth and proliferation in culture flasks (**Fig. 1A**). Next, they were seeded at low density in microplates and allowed to attach and recover for a minimum of 24 hours (**Fig. 1B**). To verify their neuronal precursor state, immunofluorescence staining (IF) for the progenitor markers Nestin, Sox1, LMX1A and TUBβ3 was performed to ensure that >80% of the cells are positive for each of these markers. In contrast, staining for the neuroectoderm marker PAX6 was negative, indicating that the cells are in an advanced neuronal progenitor stage (**Fig. 1C, D**). This observation was also supported by the presence of long, neurite-like Nestin- and TUBβ3-positive cell extensions (**Fig. 1D**). To confirm the potential of DPCs to form DNs, the medium was changed to differentiation medium, culture was continued for another 2 weeks and then tested for markers of neural differentiation (**Supplementary Figure S1A-C**). IF for the DN markers NURR1, FOXA2, and TH and the pan-neuronal marker TUBβ3 confirmed that between 60% (TH) and 80% (NURR1, FOXA2, TUBβ3) expressed these markers and displayed a neuronal morphology with condensed nuclei and cell bodies as well as an extensive neurite network. BRN2, a marker for cortical neurons [45] was not detected. To confirm the results obtained by immunofluorescence, mRNA expression profiles of hiPSCs, DPCs and DNs were compared by qPCR (**Supplementary Figure S1D**). The stem cell markers nanog and Oct3/4 were only expressed in hiPSCs, while progenitor and neuronal markers were either absent or only weakly expressed at this stage. The proliferation marker Ki67 was strongly expressed in hiPSCs and gradually downregulated during the differentiation process. Neuronal precursor markers SOX1, FOXA2 and Nestin were expressed at their highest levels in DPCs, with a gradual decline during neural differentiation. Pan neuronal markers like TUBβ3, MAP2 and NCAM1 were already expressed in DPCs and continued to be expressed later during differentiation. Markers of dopaminergic lineage were either continuously (e.g. DBH, DDC, LMX1B) or increasingly (TH) expressed from the DPC stage to 4 weeks of differentiation. Taken together, the data confirms that the hiPSC induction protocol provides a cell population that consists almost exclusively of DPCs, to be applied towards assessing the effect of small molecules on the uptake of α-syn aggregates.

**Figure 1:**
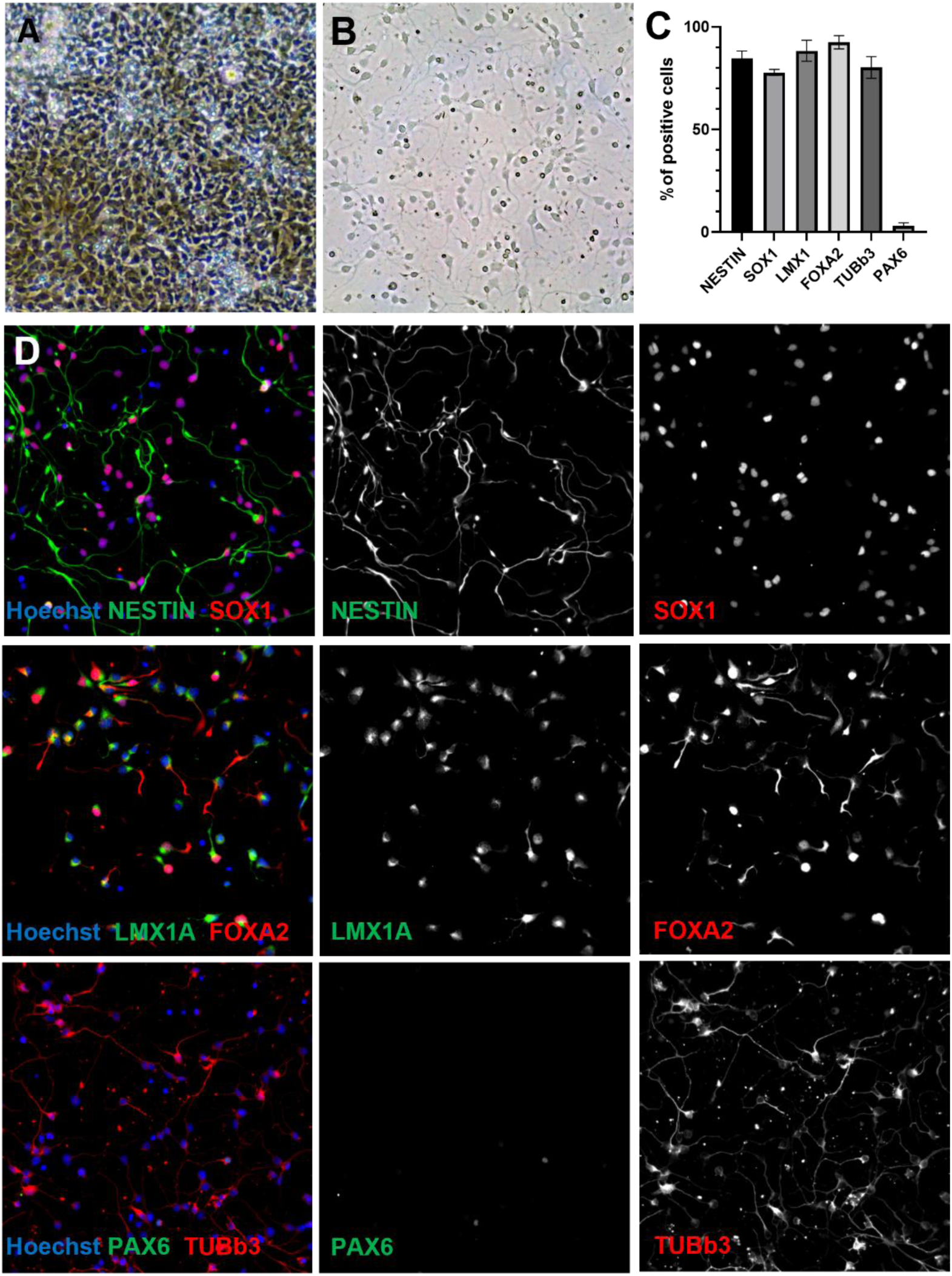
Characterization of DPCs by developmental marker expression. A) Initial expansion culture in T25 flasks at high density. B) DPCs seeded in 96 well microplates (15,000 cells/well. Bar size: 100 µm. C) Panel of immunofluorescence (IF) staining of DPCs with neural precursor markers. Bar size: 100 µm. D) Quantification of IF, percentage of marker–positive cells, mean of at least 5 independent experiments. Error bars: Standard deviation.

### 3.2. Assay optimization and analysis of preformed fibrils uptake in dopaminergic progenitor cells

During the development of the assay, we optimized several assay parameters, with a number of the most important ones described in this section. In general, DPCs were seeded in 384 well microplates and allowed to attach and recover for 24 hours before fluorophore-labeled PFF (for batch specific PFF quality control data, see **Supplementary Fig. S2**) was added to the culture media of each well. PFFs were readily taken up by DPCs and typically accumulated in the perinuclear region. They were often highly enriched on one side of the nucleus, a pattern typical for misfolded protein aggregates (**Fig. 2A-C**) [46]. To automatically quantify cellular PFF content, we optimized an image analysis protocol included in the high content microscopes software (Studio, ThermoFisher Scientific). Briefly, Hoechst-counterstained nuclei were identified as separate objects in the first channel (**Fig. 2A’**, blue border), the obtained mask was expanded to encompass the perinuclear space (**Fig. 2B’**, orange border) and the total intensity of PFF around each nucleus was measured (**Fig. 2C’**, green border). In addition, the total nuclear count, nuclear size and the average nuclear intensity of the Hoechst stain were measured as indicators of cell health.

**Figure 2:**
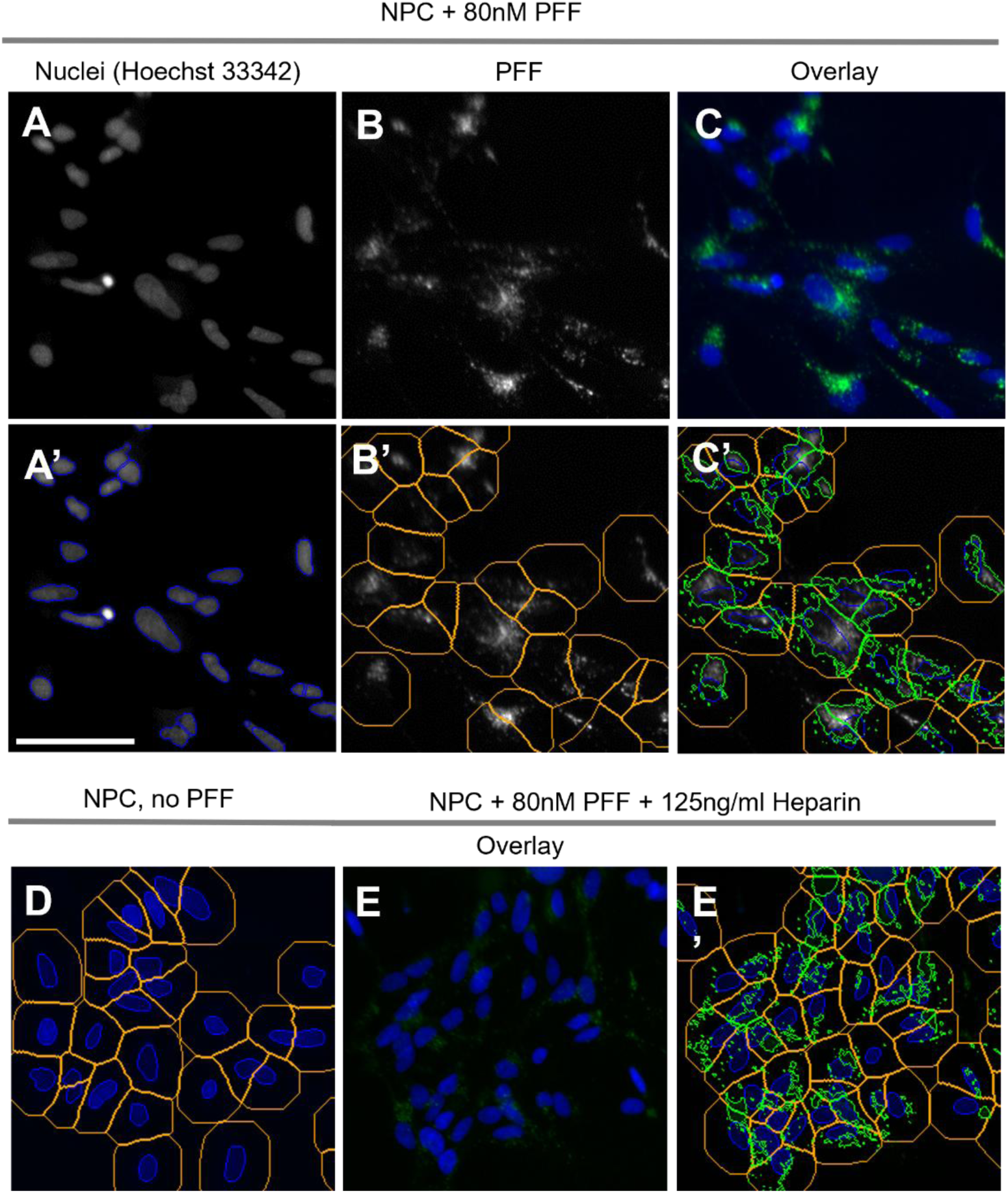
A high content image analysis sequence allows to quantify cellular PFF uptake in DPCs. A-C) DPCs + PFF. A) Channel 1, nuclei. A’) Channel 1, nuclei with object mask (blue outline). B) Channel 2, PFF. B’) Channel 2, PFF + expanded object mask (orange boundaries). C) Overlay nuclei and PFF. C’) Channel 2, PFF with nuclear mask (blue), expanded nuclear mask (orange) and identified signal within expanded mask (green boundaries). D) DPCs without PFF. Overlay of channel 1 and 2 and all masks. No green fluorescence signal in the expanded mask detected. E-E’) DPCs + PFF + Heparin. E) Channel 1+2. PFF signal is reduced. E’) Channel 1+2 + all masks. Identification of remaining PFF signal (green boundaries). Size bar: 50 µm.

Prior to fixation and counterstaining, cells were washed with PBS to remove remaining extracellular PFF. To confirm that the PBS washes were sufficient to eliminate fluorescence stemming from extracellular PFF, live cells were in parallel either washed with PBS or treated with Trypan Blue, a fluorescence quencher. Trypan Blue is frequently used in these types of experiments, since it is non-cell permeable and therefore quenches only extracellular fluorescence [47]. The measured perinuclear PFF signal intensity was at the same level after either treatment, showing that the PFF washes were effective at reducing extracellular PFF signal (**Supplementary Fig. S3**). Thus, the Trypan Blue treatment was omitted in the final PFF uptake workflow.

To determine the optimal concentration of PFFs for uptake and quantification of signal, we tested both Alexa488- and Alexa647-fluorophore-labeled PFF at a range from 20 nM to 320 nM. Both PFF versions accumulated in a similar perinuclear pattern and the signal showed a continuous concentration-dependent increase (**Fig. 3A, B** and **Supplementary Fig. S4A, B**). There was no indication of adverse effects on cell health at any of the concentrations tested, as cell counts as well as nuclear parameters were similar to non-treated controls (**Fig. 3C**, **Supplementary Fig. S4C**). Since the choice of fluorophore did not seem to affect PFF uptake, Alexa488-labeled PFF was used from this point onwards (referred to as PFF). The PFF concentration for the final protocol was set to 80 nM since, between experiments, we noted an increased variability in the intensity values at higher concentrations.

**Figure 3:**
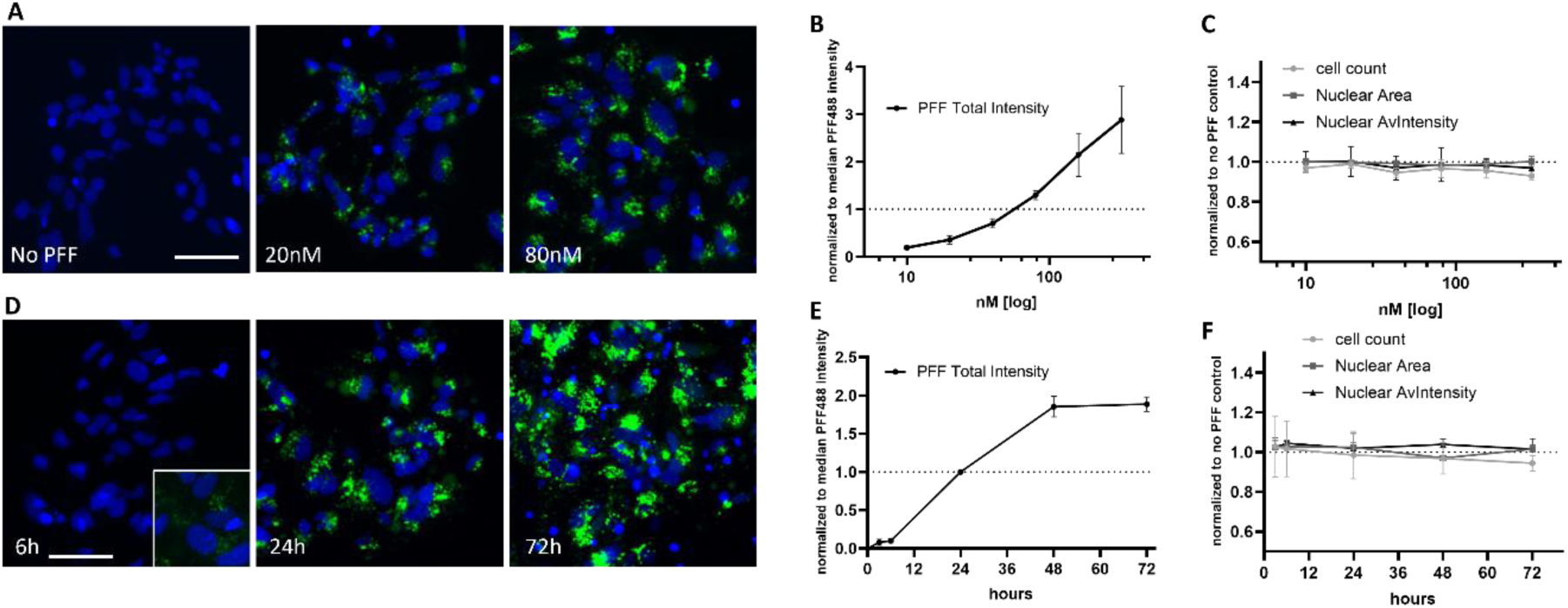
Dose and time dependence of Alexa488-PFF uptake in DPCs. A) PFF uptake at different concentration, overlay of Hoechst (blue) and PFF (green) channels. Size Bar: 50 µm. B) Quantification of cellular PFF, normalized to median total intensity of PFF. F) Quantification of cell count, nuclear size and Hoechst intensity as indicators of cell health. Graphs represent 4 independent experiments. Error bars: standard deviation. D) PFF uptake at different time points, overlay of Hoechst (blue) and PFF (green) channels. Size Bar: 50 µm. 6 hour timepoint insert: close up of nuclei with 3-fold enhanced PFF contrast. E) Quantification of cellular PFF load, normalized to median total intensity of PFF. F) Quantification of cell count, nuclear size and Hoechst intensity as indicators of cell health. Graphs represent 3 independent experiments. Error bars: standard deviation.

To determine the time dependence of uptake, cells were incubated with 80 nM PFF across a time range of 3 to 72 hours (**Fig. 3D, E**). While PFF uptake was detectable by 3-6 hours, its signal increased strongly at longer time points up to 48 hours. Longer incubation for 72 hours did not lead to a further increase, indicating that the uptake process was saturated after 48 hours. Nuclear parameters did not indicate any signs of effects on cell health, even after 72 hours (**Fig. 3F**). In all further experiments, a 24-hour incubation was used to avoid saturation while obtaining robust PFF uptake with a high signal to noise ratio.

### 3.3. Characterization of tool compounds

To date, several compounds have been published to reduce PFF accumulation in other cell models. We selected Heparin, Rottlerin [48] and Tilorone [49] to validate their inhibitory activity in our assay. These three compounds are structurally and functionally different, Heparin being a glycosaminoglycan that acts as anticoagulant, Rottlerin being a polyphenol natural product that inhibits a range of kinases and Tilorone being a synthetic small molecule with antiviral activity. When added 1h prior to PFFs, all three compounds reduced intracellular PFF load in a dose-dependent manner, at higher doses attaining at least 80% reduction compared to untreated controls (**Fig. 4**, see also **Fig. 2E**). In addition, the measurements of nuclear parameters allowed us to add more information regarding changes in cell state due to the compounds. Most notably, Rottlerin treatment, at 4 µM and 8 µM, led to a reduction in the average nuclear size, corresponding to an increase in the average nuclear staining intensity and a small, but consistent drop in cell number (**Fig. 4B**). This likely indicates an adverse effect on cell health at these high doses, potentially through apoptosis. Interestingly, Heparin treatment gradually reduced the cell count over a broad concentration range, leading to a ∼20% lower cell count at the highest dose, without any effect on nuclear parameters (**Fig. 4A**). This effect is potentially caused by Heparin reducing cell adhesion, leading to an increased detachment of cells and reduced numbers by the time of assay readout [50]. Tilorone had no significant effect on nuclear parameters up to the highest dose tested (**Fig. 4C**). Taken together, these results confirm that the PFF uptake assay could consistently assess small molecule-induced changes in the cellular PFF load, accompanied by measurements for any potential effects on cell health and viability.

**Figure 4:**
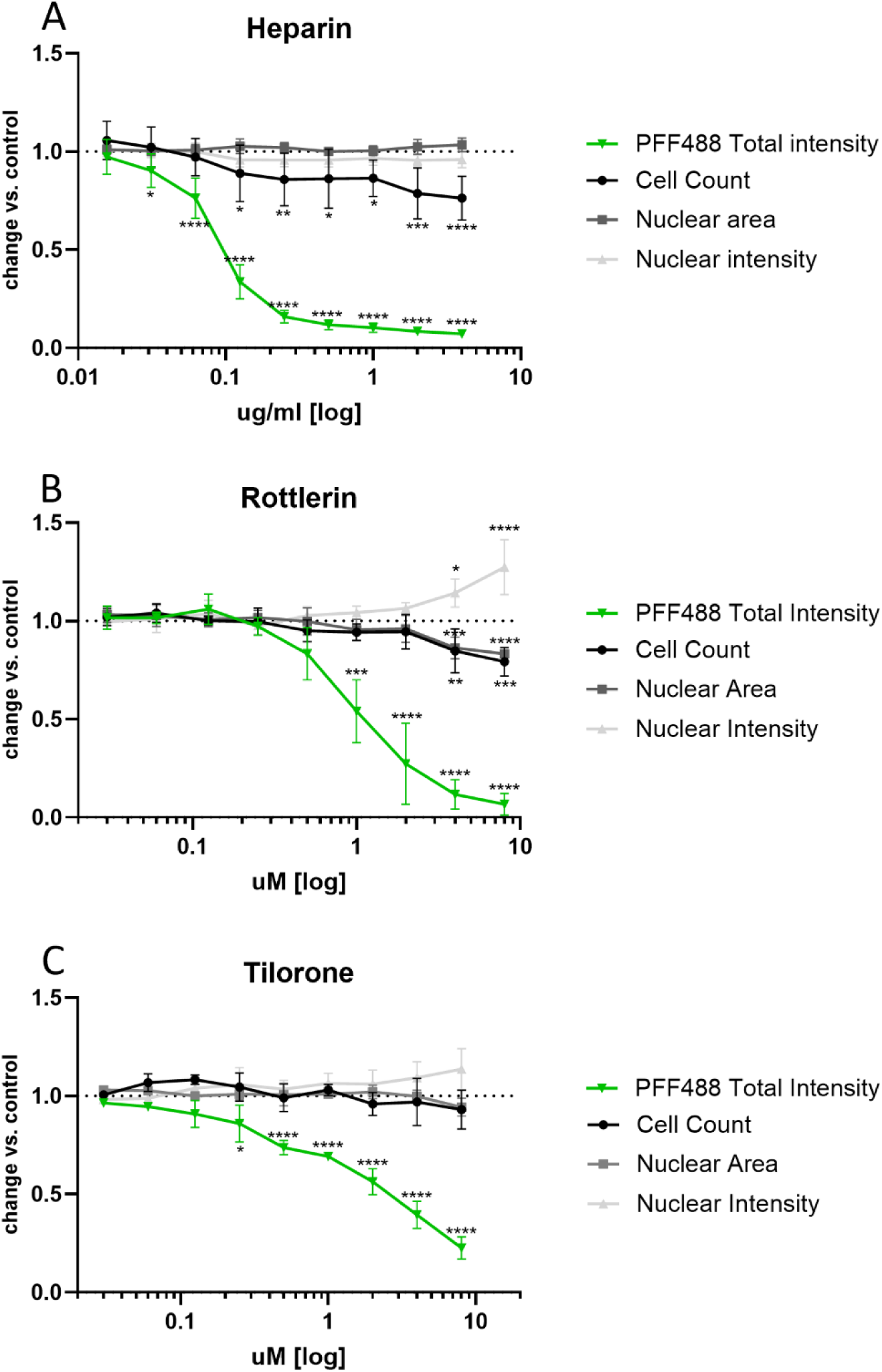
Tool compounds reduce intracellular PFF accumulation. PFF total intensity, nuclear area and mean nuclear intensity per cell are averaged per well. Cell count mean per well. Values are normalized to vehicle-only (DMSO) control, (control=1). A) Heparin (µg/ml). B) Rottlerin (µM). C) Tilorone (µM). Results present the mean of at least 4 independent experiments. Bars represent standard deviation. P-values: **** <0.0001 to *<0.1, one-way ANOVA and Dunnett’s corrected multiple comparisons test.

### 3.4. Screening a library of known pharmacologically active compounds

With the PFF assay and tool compounds in place, we next explored whether our workflow could allow us to identify other small molecules that reduce intracellular load of PFF. To this end we screened the Prestwick chemical library® of 1520 FDA and/or EMA approved compounds (**Fig. 5**). The compounds were distributed into five 384 well plates. The exact plate layout is illustrated in the schematic in Figure 5. In each plate, Rottlerin and Tilorone were added as positive controls and DMSO-only in several columns as the negative control. PFF addition was omitted in the last DMSO-only column. This column served as an additional control to confirm the accuracy of the threshold for PFF detection during the analysis and to detect any green fluorescence artefacts in the culture. After cell seeding and incubation for 24 hours, the source plates were pinned to their respective screening plates at a final concentration of 8 µM. After 1 hour of incubation, PFFs were added to the screening plates and after an incubation period of 24 hours the plates were processed, imaged and analysed. Normalization of the compound effects against the DMSO-only controls was performed separately for each plate first and then the data was combined into the full data set.

**Figure 5:**
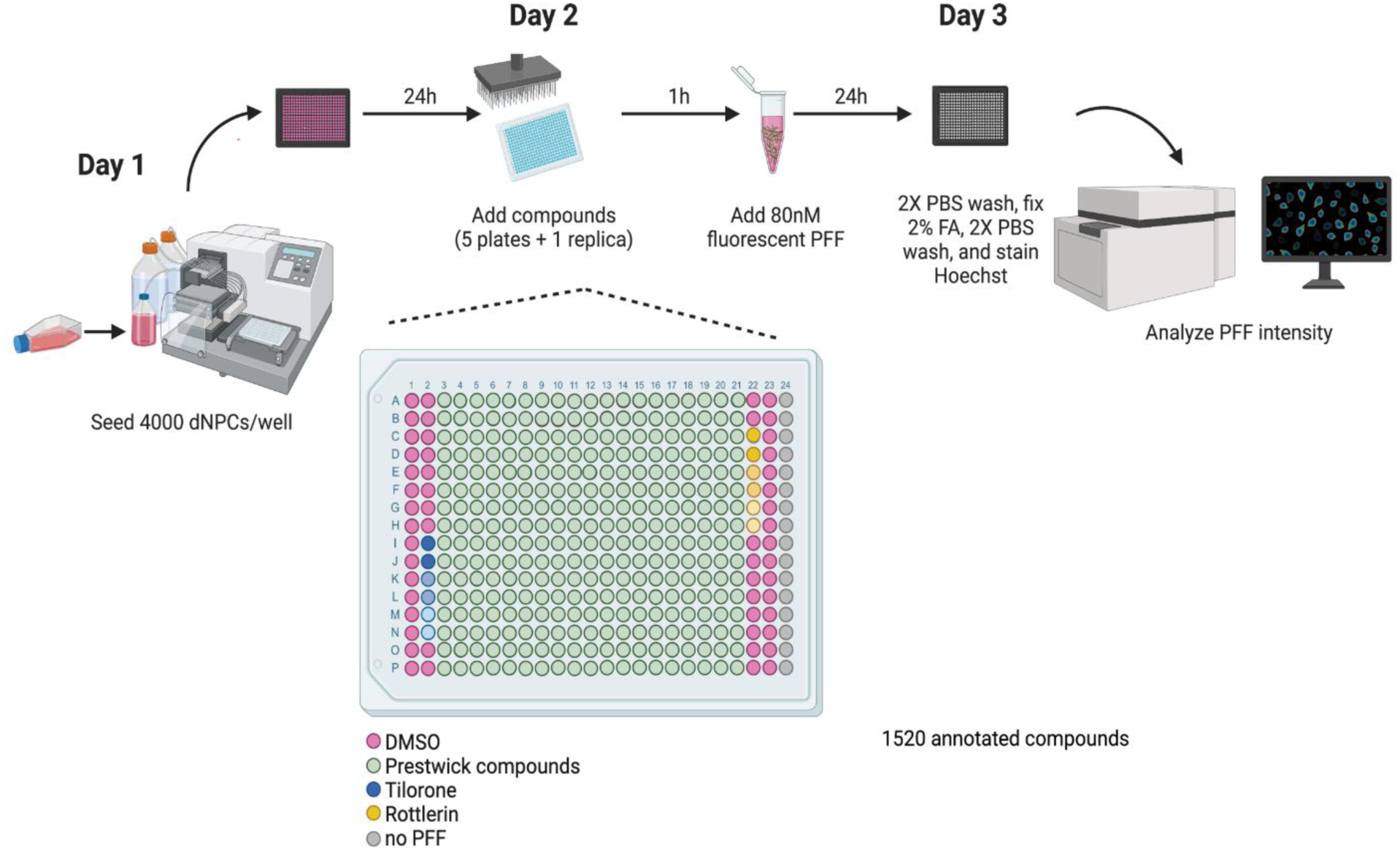
Schematic of screening workflow. DPCs were seeded in 384 well microplates at 4000 cells/well. After 24 hours recovery, library compounds were pinned into the test plates. Per plate 320 compounds at a final concentration of 8 µM were added. 5 plates were processed to transfer the 1520 compounds of the Prestwick library. In addition, one plate was replicated to verify the quality of the set. One hour later PFF is added to the wells (80 nM final concentration). 24 hours later test plates were washed with PBS, formaldehyde fixed and counterstained. Images were acquired at x10 magnification. Platemap: Each plate contained 320 test compounds, positive control compounds Tilorone and Rottlerin, vehicle only treated wells and one column without PFF.

Plotting the complete data set (**Fig. 6A**) shows that the PFF intensity values of the DMSO-only controls distribute in a range from −20% to +20% around the mean (standard deviation: 7.7%). Thus, this range of signal fluctuation is intrinsic to the assay (**Fig. 6A**, grey spots). In contrast, a substantial number of library compounds decreased the signal further, indicating compound specific effects. In addition, a few compounds increased the PFF signal above +20% (**Fig. 6A**, dark blue spots). In total, 108 compounds decreased PFF levels by at least 50% (7.1% of all compounds, **Fig. 6B**). However, a considerable number of these compounds also strongly reduced the cell count. While cell counts in DMSO-only control wells fluctuated between −20% and +20% (standard deviation: 7.5%), these compounds reduced cell count further. There is a clear correlation between reduced cell count and reduced PFF load, indicating that cell stress and toxicity is a confounding factor in the assay. Adding cell count to the selection criteria reduced the number of active compounds to 60 and 34 at a −30% cell count threshold and a −20% cell count threshold, respectively (**Fig. 6B**). The complete data set, including all normalized PFF and cell parameter measurements is provided in **Supplementary Table S1**.

**Figure 6:**
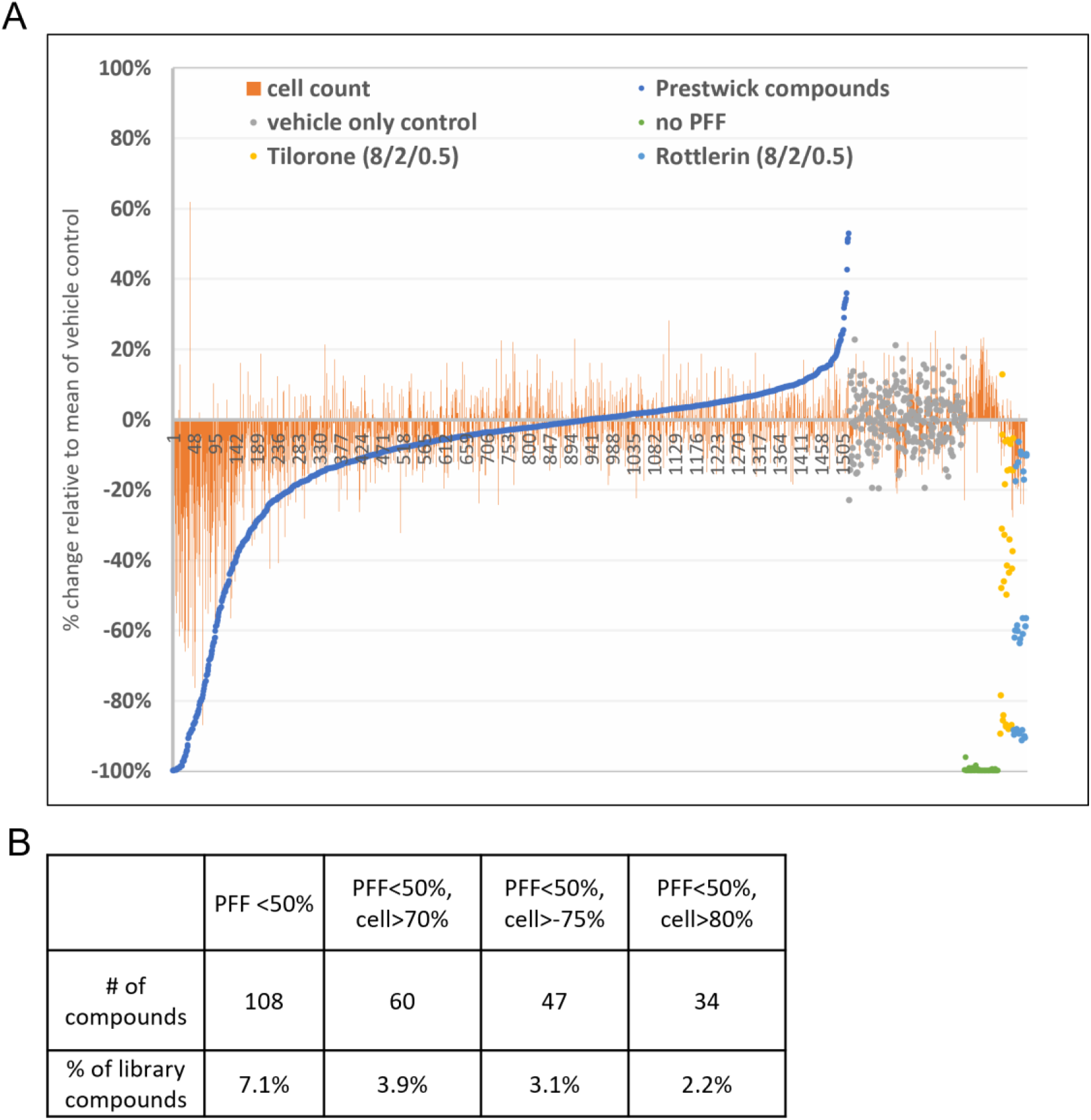
Summary of screening results. A) Waterfall plot of PFF uptake inhibition screen results (change in PFF intensity) with Prestwick library compounds. Data is normalized (per plate) against mean of vehicle (DMSO) control. Blue: Compound treated wells. Grey: Vehicle-only. Blue: Rottlerin treated wells (8, 2, 0.5 µM). Yellow; Tilorone treated wells (8, 2, 0.5 µM). Green: Control wells without PFF. Orange bars: Relative cell count in the well corresponding to data point. B) Number of compounds that reduce PFF load by at least 50% compared to vehicle only control with/ without additional filtering by cell number.

To ensure the accuracy and robustness of the assay during screening, we verified the following controls: the positive controls Rottlerin and Tilorone reduced PFF intensity in a dose-dependent manner (**Fig. 6A**, yellow and light blue spots, also **Supplementary Fig. S5A**). The effective concentration range was similar to the range determined during assay optimization (**Fig. 4**), also confirming the accuracy of the compound transfer. Furthermore, Pearson correlation coefficient values that were calculated between a screening plate and an additionally included replica plate ranged from 0.5 for cell count to 0.8 for PFF intensity and nuclear size, confirming the reproducibility between plates (**Supplementary Fig. S5B**). The relatively high number of controls over all plates allowed us also to determine Z-prime values as an indicator of dynamic range of PFF signal in the screen [51]. Values were 0.75 between PFF only and no PFF controls, 0.69 between PFF only and 8 µM Rottlerin and 0.60 between PFF only and 8 µM Tilorone. These values are well above the Z-prime value of 0.5 that is widely considered the threshold for an excellent range between positive and negative controls [52]. Taken together, the control data confirmed the quality of the assay and allowed us to set an appropriate threshold to select potentially active compounds.

### 3.5. Confirmation of active compounds in dopaminergic progenitor cells

To confirm active compounds, we selected 52 compounds that decreased PFF levels by a minimum of 50% and reduced cell counts by no more than 30%. Retesting was done in a concentration range from 12 to 0.4 µM (**Fig. 7**). At 12 µM (therefore at a 4 µM higher concentration then in the original screen), for 50 out of 52 compounds the reduction of PFF intensity was confirmed, 11 of these compounds showed no adverse effects on cell count and nuclear size (at thresholds). On the other hand, 30 compounds showed effects on cell health, reducing cell number by more than 30% and shrinking nuclei by more than 20%. The remaining 9 compounds reduced cell count but did not affect nuclear size. At 4 µM the number of compounds that reduced PFF intensity without affecting cell health peaked at 21 compounds, at 1.2 µM and 0.4 µM the number fell to 8 and 6 compound, respectively. Overall, 17 out of 52 compounds showed a dose response, reducing PFF intensity at least at 2 concentrations and had no adverse effects on cell health, using cell count and nuclear area as threshold parameters (**Supplementary Figure S6**). Sorting the confirmed compounds by target/ activity annotations showed that several compounds are functionally and/ or structurally related. For example, 3 compounds belong to the quinoline-related group of plasmodicial drugs and 4 compounds to the cardiac glycoside group of sodium potassium adenosine triphosphatase inhibitors (**Table 3**, see also discussion).

**Figure 7:**
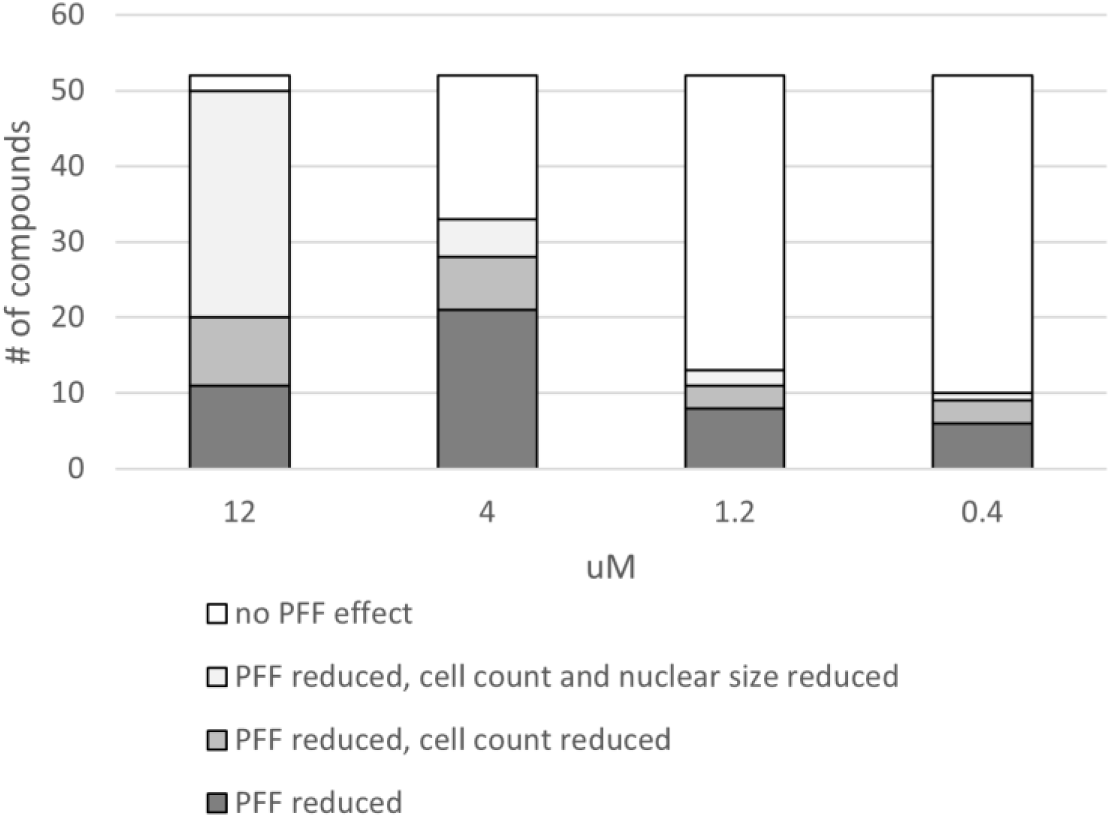
Active confirmation of 52 compounds at 4 doses. Summary of results. At each concentration, compounds are grouped in either PFF reduced and little effect on cell parameters (PFF < 70%, cell count > 70% and nuclear size> 80%), PFF reduced and cell count reduced (PFF < 70%, cell count <70%, nuclear size > 80%), all parameters reduced (PFF < 70%, cell count <70%, nuclear size < 80%) or no effect (PFF > 70%). All measurements are normalized against the mean of PFF + vehicle only control (=100%).

### 3.6. Compound validation in dopaminergic neurons

While the use of DPCs allows to screen a large number of compounds efficiently, ultimately the activity of isolated candidate molecules needs to be retained in differentiated neurons to be of value. Potentially, these molecules could be sensitive to the changes in expression profiles and physiology that occur during differentiation, reducing or eliminating their activity. To test the inhibitory effect of compounds in differentiated DNs, DPCs were seeded in 96 well plates in differentiation medium and cultured for 2 weeks before performing the PFF uptake assay. At this point the cells have adopted a clear neural morphology (**Fig. 8A and B**) and robustly express markers of dopaminergic neurons (**Supplementary Fig. S1**). We retested 11 out of the 17 active molecules across different concentrations, with the remaining conditions of the assay being kept as before. Similar to our observations in DPCs, PFF accumulates in DNs in a unipolar, perinuclear localisation in the cell body (**Fig. 8A**). We noted that, compared to DPCs, the PFF intensity in DNs was markedly lower, indicating that PFF uptake in DNs is less efficient than DPCs. However, all 11 compounds, as well as the Heparin control, showed a clear dose dependent reduction of intracellular PFF accumulation with a maximum reduction ranging from 60% to 80% compared to PFF-only controls (**Fig. 8C**). Using cell count as an indicator for cell toxicity, only 3 compounds (Quinacrine, Bosutinib, Cycloheximide) showed, at the highest doses, a significant reduction in cell number. These compounds still reduced PFF intensity at doses 9 to 27-fold lower than the dose that reduced cell number, indicating that, at the low doses, the inhibition of PFF accumulation is not a direct effect of toxicity. Taken together, the observation that the compounds are effective in blocking PFF accumulation both in DPCs and DNs confirms our reasoning that screening in DPCs, allowing to test thousands of molecules efficiently, yields active compounds that translate well to differentiated neuronal cultures. This also indicates that neuronal cells at both developmental stages employ similar mechanisms to take up and/ or transport PFFs. Therefore, this strategy provides an adaptable and scalable tool to screen for therapeutics that modulate the uptake of α-synuclein aggregates in DNs.

**Figure 8:**
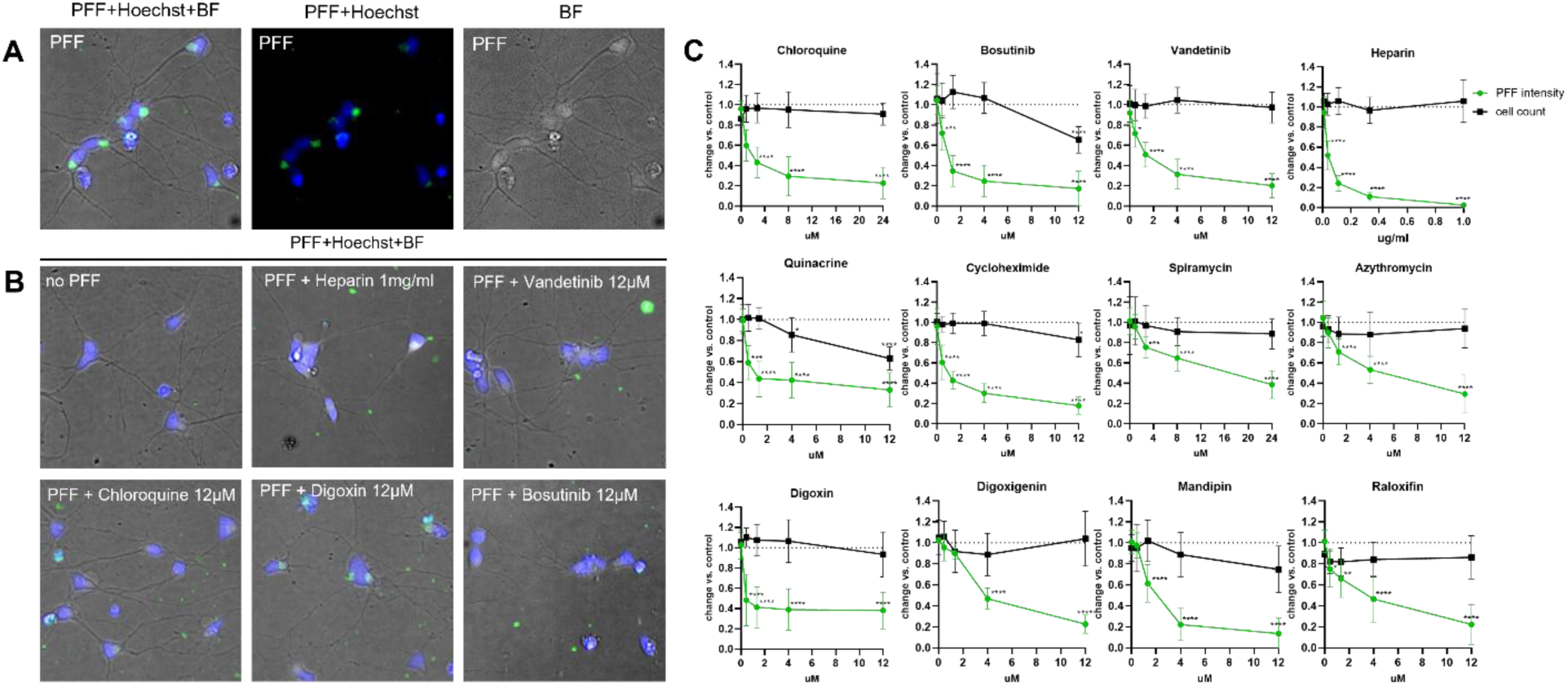
Screening actives reduce PFF content in DNs. A) PFF uptake in 2 week differentiated DN. PFF is concentrated in a perinuclear, polarized fashion. Channel overlays from left to right: PFF (green)/Hoechst (nucleus, blue)/ BF (brightfield), PFF/Hoechst, BF only. B) Treatment examples and no-PFF negative control. Channel overlay PFF /Hoechst/ BF. Bar: 15 µm. C) Dose dependent reduction in intracellular PFF and cell count. Values are normalized to vehicle-only (DMSO) control, (control=1). Results present the mean of 3 independent experiments. Error bars represent standard deviation. P-values: **** <0.0001 to *<0.1, one-way ANOVA and Dunnett’s corrected multiple comparisons test.

## 4. Discussion

In this study, we present a semi-automated workflow to quantify changes in cellular uptake of PFF. We optimised the steps of the workflow and used high content imaging and analysis tools to build a simple and effective process pipeline, while still being able to provide cell level data including indicators of cell health. Screening a small molecule library of annotated pharmacologically active compounds confirmed the robustness of the workflow and led to the identification of several compounds that strongly reduce PFF accumulation in DPCs. Importantly, we were able to reproduce the inhibitory effects of these compounds in differentiated DNs. A subset of the compounds is well characterized and known to have different cellular targets, indicating that they affect different aspects of PFF uptake.

Using DPCs as a cellular model, our global aim was to optimize the different components of the workflow to render it scalable for screening a large number of small molecules. Starting with DPCs laid the foundation for generating a high number of frozen cell stocks that can be readily used and handled in many ways similarly to immortalized cell lines. After seeding in 384-well plates, the workflow takes only 48 hours before cells are ready for imaging and the steps to prepare the plates for imaging (washes, fixation, nuclear labeling) or optimized for minimal plate manipulation and reagent use. This makes the workflow cost efficient and, most importantly, reduces potential sources of variability. To confirm the accuracy of the workflow, we measured several parameters of reproducibility during the library screen and achieved excellent values in all of them, including Z-prime, positive controls and replica correlation. Despite its relative simplicity, the workflow, in addition to the quantification of PFF load, also provides the means to collect measurements of nuclear parameters on the cellular level, thus providing important information about the cell state under each treatment condition. These parameters, combined with cell counts per well, provide the basis for identifying conditions that elicit cellular toxicity. This step is particularly important since cell death leads to the loss of cytoplasmic material, including intracellular PFF, which would have led to a high number of false positive compounds. Identifying these compounds during screening considerably reduced the number of compounds that would have been unnecessarily moved forward into further testing. We also noted, as exemplified with Heparin treatment, that conditions of reduced cell count, but control-like nuclear parameters can allow the identification of compounds that can potentially affect cell adhesion rather than cell health directly. Taken together, the workflow provides a robust tool to measure intracellular PFF levels and the effect of small molecules on PFF levels and cell state.

While differentiated DNs in culture would potentially better resemble DNs of the brain, deploying them in an automated, high throughput environment that is required in mid to large scale screens remains challenging [35, 36]. For instance, the long timeframe of differentiation in microplates can potentially amplify differences in well populations, plate position effects (e.g. edge effects) and increase the risk of contamination. In addition, the extensive neuronal networks that form during differentiation render the culture more delicate. In our experience, as differentiation progresses, it substantially increases the risk of partial or complete detachment during automated pipetting steps, especially on the small well surfaces of 384 well plates. DPCs, in contrast, provide a comparatively uniform and robust model, while providing a genetically and physiologically background that is more relevant than that of immortalized cell lines. Using IF and qPCR, we showed that the DPCs used in the assay already express a set of dopaminergic markers, that the early NPC marker PAX6 is suppressed and that the cells display neurite-like extensions after 1 day of plating. We also found that their potential to proliferate is already relatively low at that point, with Ki67 expression reduced and cell density remaining stable over the time frame of the assay (not shown). Lastly, we tested the activity of candidate molecules on cultures of differentiated DNs and confirmed their ability to reduce PFF accumulation. Overall, our assay provides an ideal platform for assessing large numbers of small molecules in neuronal cells by providing a balance between physiological relevance and throughput.

To isolate novel compounds that reduce intracellular accumulation of PFF, we screened 1520 FDA and EMA approved compounds. We found 17 compounds that strongly reduced PFF accumulation, at a minimum of 2 out of 4 tested concentrations while passing cell health criteria. Three of these compounds (Chloroquine, Amodiaquine and Quinacrine) belong to the same group of structurally related quinoline anti-malaria drugs, known to act as lysosomotropic agents [53]. Their partial hydrophobicity allows them to pass membranes, however as weak bases they become protonated in an acidic environment, as in lysosomes and late endosomes, which adds a positive charge and traps them in the cellular compartment. The accumulation of such molecules interferes with lysosomal function and induces secondary effects, which can include exocytosis and increased synthesis of lysosomal and autophagic proteins [54]. It has recently been demonstrated that PFF uptake in DNs is mediated by a novel form of macropinocytosis and that PFFs appears in acidic vesicles within minutes after addition to the medium [29]. This finding suggests that lysosomotropic compounds can directly interfere with cellular import in these cells. However, the assay does not distinguish between a direct inhibition of uptake at the plasma membrane and potential later intracellular events that could reduce PFF accumulation. It has been shown, for example, that interference with lysosomal function leads to increased lysosomal exocytosis and an increase in extracellular α-syn in cell models overexpressing α-syn [55, 56]. In this context it is worth noting that many PD risk genes have been found to be related to lysosomal function and dynamics. Deciphering their involvement in PD progression and their potential as therapeutic targets has become a focal point of PD research [57].

We also found 4 active compounds belong to the group of cardiac glycosides: Digoxigenin, Digoxin, Digitoxigenin and Lanatoside C. These compounds inhibit sodium potassium adenosine triphosphatase (Na+/K+ ATPase) at the plasma membrane, which leads directly to an increase in intracellular sodium and indirectly to an increase in intracellular calcium levels [58]. Interestingly, α3β1 Na+/K+ ATPase is one of several transmembrane proteins at the cell surface that have been shown to interact with α-syn aggregates [59, 60]. Binding of α-syn aggregates leads to clustering of the protein complex in neurons and appears to weaken its Na+-pumping activity [59]. To our knowledge, there is no evidence published that this binding and clustering can lead to cell entry of PFFs. Thus, direct interference of cardiac glycosides with endocytosis of PFF through the inhibition of Na+/K+ ATPase complexes seems unlikely. Alternatively, since cardiac glycosides interfere with ion homeostasis, this might indirectly affect vesicle dynamics and reduce PFF accumulation by interfering with cellular trafficking [61].

We also identified two tyrosine kinase inhibitors: Bosutinib and Vandetanib. These molecules are structurally closely related, but differ in their main targets, which are src/Abl and VEGFR/EGFR, respectively [62, 63]. Both targets have been implicated in PFF uptake previously. It has been reported that inhibition of c-src with Sarcatinib, another src/Abl kinase inhibitor reduces PFF uptake in rat primary cortical neurons [64] and EGFR inhibition with the highly selective EGFR inhibitor AZD3759 reduces PFF uptake in mouse cortical neurons and attenuates PFF induced pathology in a preclinical mouse model [65].

Cycloheximide, an inhibitor of eukaryotic translational elongation, was also among the active molecules, indicating that protein synthesis is required for intracellular PFF accumulation. With Cepharantine, we isolated an antiviral compound, in addition to the positive control Tilorone. While structurally different, these 2 compounds potentially share activity features, since they have been found to be active in a number of different viral entry screens [66, 67]. The remaining 6 active molecules cover a wide spectrum of annotations (Table 1). It should be noted that we identified all cardiac glycosides and all anti-malaria quinolones contained in the compound library, again highlighting the efficiency and accuracy of the assay. Whether these drugs would have any therapeutic benefit in PD, remains to be established, but clearly they can elicit an inhibitory effect on uptake and accumulation of α-syn aggregates in midbrain neurons.

**Table 1.** List of primary and secondary antibodies.

| Antibody | Source | Cat# | RRID | dilution |
| --- | --- | --- | --- | --- |
| Nestin | Abcam | AB92391 | AB_10561437 | 1/150 |
| SOX1 | R&D system | AF3369 | AB_2239879 | 1/100 |
| LMX1A | MilliporeSigma | AB10533 | AB_10805970 | 1/500 |
| FOXA2 | R&D system | AF2400 | AB_2294104 | 1/200 |
| TUBb3 | MilliporeSigma | AB9354 | AB_570918 | 1/3000 |
| PAX6 | BioLegend | 901301 | AB_2749901 | 1/200 |
| Nurr1 | Santa Cruz<br>Biotechnology | SC-990 | AB_2630633 | 1/100 |
| MAP2 | Encor Biotech | CPCA-MAP2 | AB_2138173 | 1/500 |
| TH | Peel -Freez | P40101-150 | AB_2617184 | 1/400 |
| BRN2 | Cell Signaling<br>Technology | 12137 | AB_2797827 | 1/500 |
| Dylight 488 donkey anti-rabbit IgG | Abcam | ab96891 | AB_10679664 | 1/500 |
| AlexaFluor647 donkey anti-chicken IgY | Jackson Immuno<br>Research | 703-605-155 | AB_2340379 | 1/500 |
| AlexaFluor647 donkey anti-goat | Invitrogen | A21447 | AB_141844 | 1/500 |
| Dylight 550 donkey anti-mouse | Abcam | ab96876 | AB_10679663 | 1/500 |

**Table 2.** List of primers.

| Gene ID | Gene Name | Accession number | Assay ID |
| --- | --- | --- | --- |
| ACTB | Actin, beta | NM_001101.3 | Hs01060665_g1 |
| MAP2 | Microtubule associated protein 2 | NM_001039538.1 | Hs00258900_m1 |
| NANOG | NANOG homeobox | NM_001297698.1 | Hs02387400_g1 |
| NCAM1 | Neural cell adhesion molecule 1 | NM_000615.6 | Hs00941830_m1 |
| NES | Nestin | NM_006617.1 | Hs04187831_g1 |
| POU5F1 (OCT3/4) | POU class 5 homeobox 1 | NM_001173531.2 | Hs04260367_gH |
| TUBB3 | Tubulin beta 3 class III | NM_001197181.1 | Hs00801390_s1 |
| mKi67 | Marker Of Proliferation Ki-67 | NM_020346.2 | Hs01032443_m1 |
| DBH | Dopamine Beta-Hydroxylase | NM_000787.3 | Hs01089840_m1 |
| DDC | Dopa Decarboxylase | NM_001082971.2 | HS01105048-m1 |
| EN1 | Engrailed Homeobox 1 | NM_001426.3 | Hs00154977_m1 |
| FOXA2 | Forkhead Box A2 | NM_021784.4 | Hs00232764_m1 |
| HOXA2 | Homeobox A2 | NM_006735.3 | Hs00534579_m1 |
| LMX1B | LIM Homeobox Transcription Factor 1 Beta | NM_001174146.1 | Hs00158750_m1 |
| MAOB | Monoamine Oxidase B | NM_000898.4 | Hs01106246_m1 |
| NURR1 | Nuclear Receptor Subfamily 4 Group A Member 2 | NM_006186.3 | Hs01117527_g1 |
| TH | Tyrosine Hydroxylase | NM_000360.3 | Hs00165941_m1 |
| GAPDH | Glyceraldehyde-3-Phosphate Dehydrogenase | NM_001256799.2 | Hs02786624_g1 |
| SOX1 | SRY-box 1 | NM_005986.2 | Hs01057642_s1 |

**Table 3.**
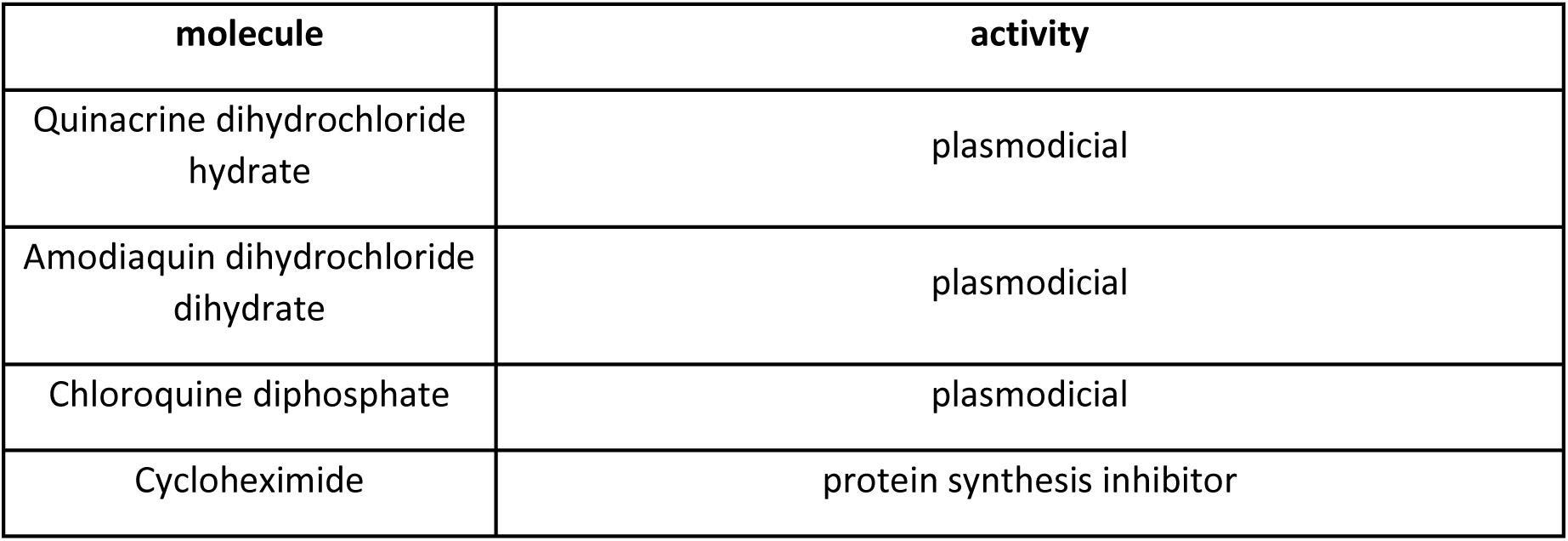
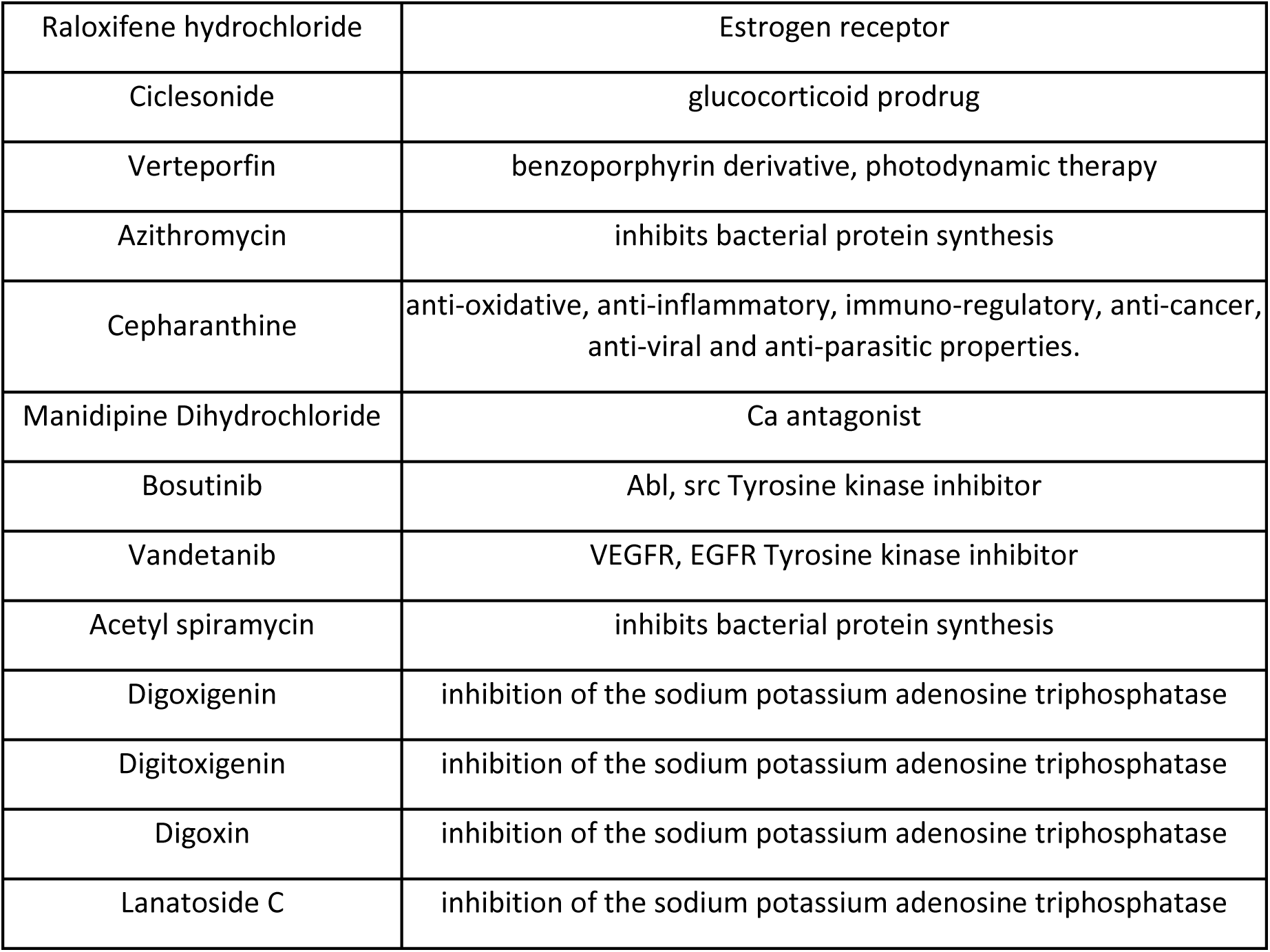
List of compounds that reduced PFF content by more than 30% at least at 2 concentrations (with low/ no toxicity, (cell count >70%, Nuclear Area >80%)).

| molecule | activity |
| --- | --- |
| Quinacrine dihydrochloride hydrate | plasmodicidal |
| Amodiaquin dihydrochloride dihydrate | plasmodicidal |
| Chloroquine diphosphate | plasmodicidal |
| Cycloheximide | protein synthesis inhibitor |
| Raloxifene hydrochloride | Estrogen receptor |
| Ciclesonide | glucocorticoid prodrug |
| Verteporfin | benzoporphyrin derivative, photodynamic therapy |
| Azithromycin | inhibits bacterial protein synthesis |
| Cepharanthine | anti-oxidative, anti-inflammatory, immuno-regulatory, anti-cancer, anti-viral and anti-parasitic properties. |
| Manidipine Dihydrochloride | Ca antagonist |
| Bosutinib | Abl, src Tyrosine kinase inhibitor |
| Vandetanib | VEGFR, EGFR Tyrosine kinase inhibitor |
| Acetyl spiramycin | inhibits bacterial protein synthesis |
| Digoxigenin | inhibition of the sodium potassium adenosine triphosphatase |
| Digitoxigenin | inhibition of the sodium potassium adenosine triphosphatase |
| Digoxin | inhibition of the sodium potassium adenosine triphosphatase |
| Lanatoside C | inhibition of the sodium potassium adenosine triphosphatase |

In recent years, collections of patient-derived hiPSCs have been established and continue to grow across different sources. In addition to screening for novel inhibitors of PFF uptake in one hiPSC line with a mechanistic or therapeutic aim, this opens the possibility to expand the PFF uptake assay towards comparing uptake across multiple PD patient cell lines. It would for example be interesting to investigate if certain PD patient lines with known PD mutations or even sporadic PD patient lines with no identified PD-related genetic defect can be differentiated by their ability to take up PFFs. The approach we report here provides a simple workflow and a selection of tool compounds that can be adapted to pursue research questions like these.

## Acknowledgements

This work was supported by the Canadian consortium on neurodegeneration in aging (CCNA), the Canada First Research Excellence Fund, awarded through the Healthy Brains, Healthy Lives initiative (HBHL) at McGill University, the New Frontiers in Research Fund (NFRF) funded TRIDENT initiative, the Canadian institutes of health research (CIHR), the Michael J. Fox foundation (Grant # MJFF-025378), GBA1 Canada and the Quebec consortium for drug discovery (CQDM). E.A.F. is supported by a Canada Research Chair (Tier 1) in Parkinson’s disease.

## CRediT Author Contributions

Conceptualization: W.E.R., T.M.D., A.I.K, C.H.; Methodology: W.E.R., T.P., C.H., A.I.K., T.M.D; Validation: W.E.R., A.I.K., T.M.D.; Data curation: W.E.R., A.I.K.; Investigation: W.E.R., A.I.K., C.H., E.N.-R., C.X.-Q.C., I.S., W.L.; Writing—first draft preparation: W.E.R., A.I.K.; Writing—review and editing: W.E.R., A.I.K.,T.M.D.; Supervision, T.M.D.; Project administration: W.E.R.,T.M.D.; Funding acquisition-resources: T.P., E.A.F, T.M.D. All authors have read and agreed to the published version of the manuscript.

