## Supplemental Data 1 for "A high-content imaging workflow to screen for molecules that reduce cellular uptake of α-synuclein preformed fibrils"

### Supplementary Data 1

Supplementary Figure S1:

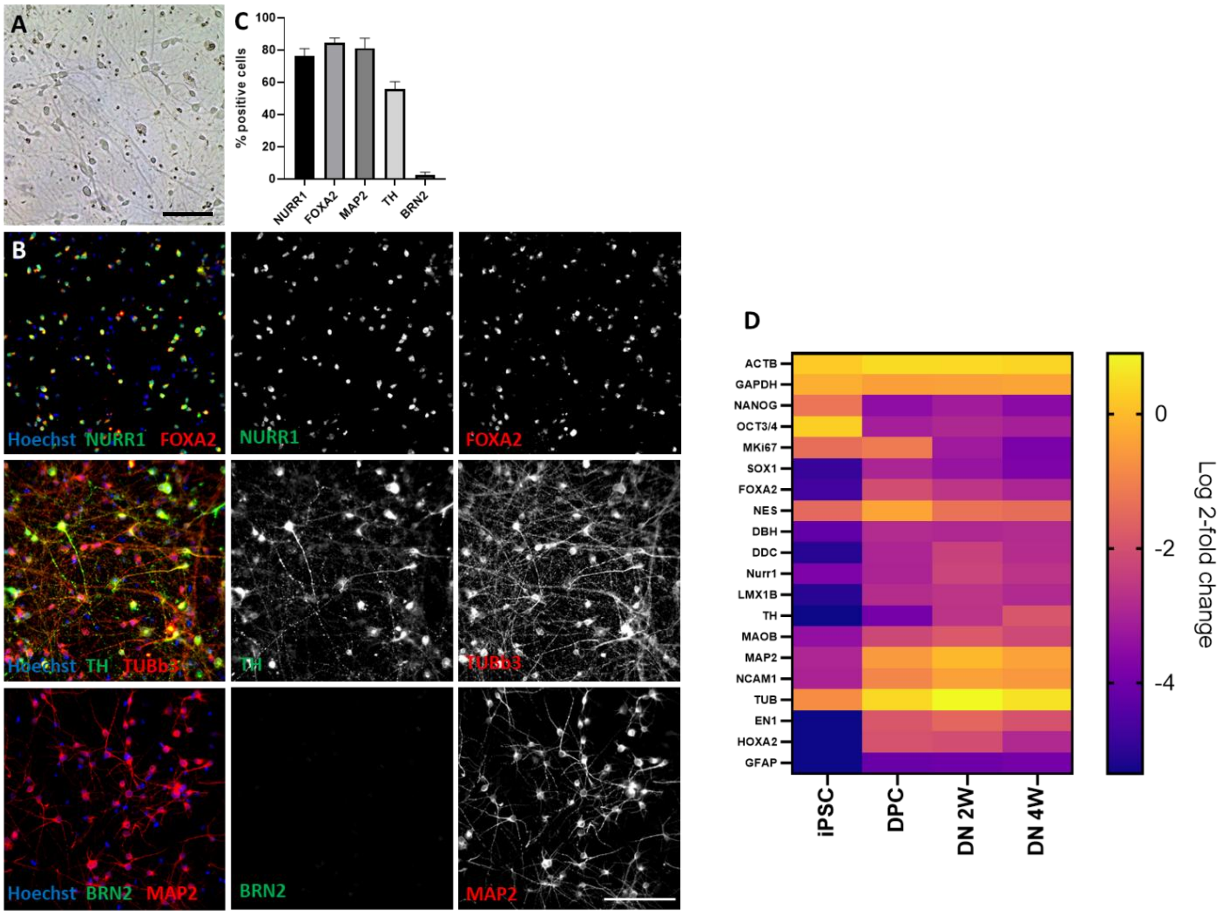


Supplementary Figure S1: Dopaminergic neurons (DN) after 2 weeks differentiation and developmental stage-dependent expression profiles. A) Brightfield image of 2week DN differentiated in 96 well microplates. Bar size: 100µm. B) Panel of immunofluorescence (IF) staining of 2-week DN with neural markers. Bar size: 100µm. C) Quantification of IF, percentage of marker –positive cells, mean of at least 5 independent experiments. Error bars: Standard deviation. D) Expression of marker genes at different stages of differentiation. Values are normalized against actin (ACTB) and GAPDH for each stage of differentiation.

Supplementary Figure S2:

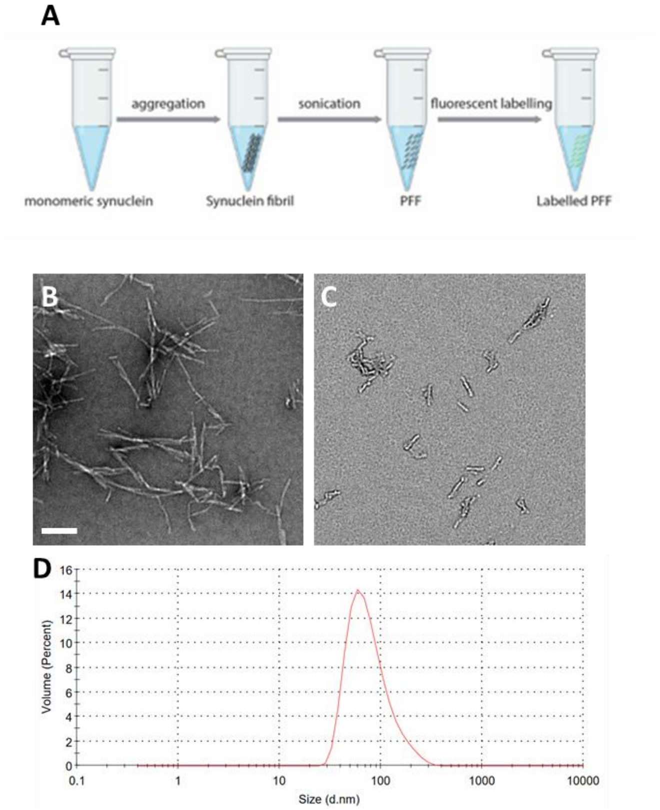


Supplementary Figure S2: PFF quality control for PFF batch used in screen. A) Schematic of PFF generation from monomeric synuclein. B-C) Electron microscopy of synuclein fibrils before (B) and after (C) sonication. Bar size: 200nm. D) Size distribution measurement with Dynamic Light Scattering (DLS) analysis, average size: 70nm.

Supplementary Figure S3:

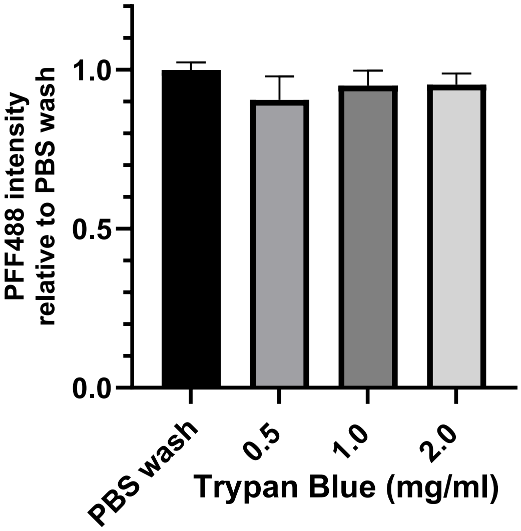


Supplementary Figure S3: PBS washes are as effective as Trypan Blue mediated quenching in reducing extracellular Alexa488-fluorophore signal. Error bars: standard deviation.

Supplementary Figure S4:

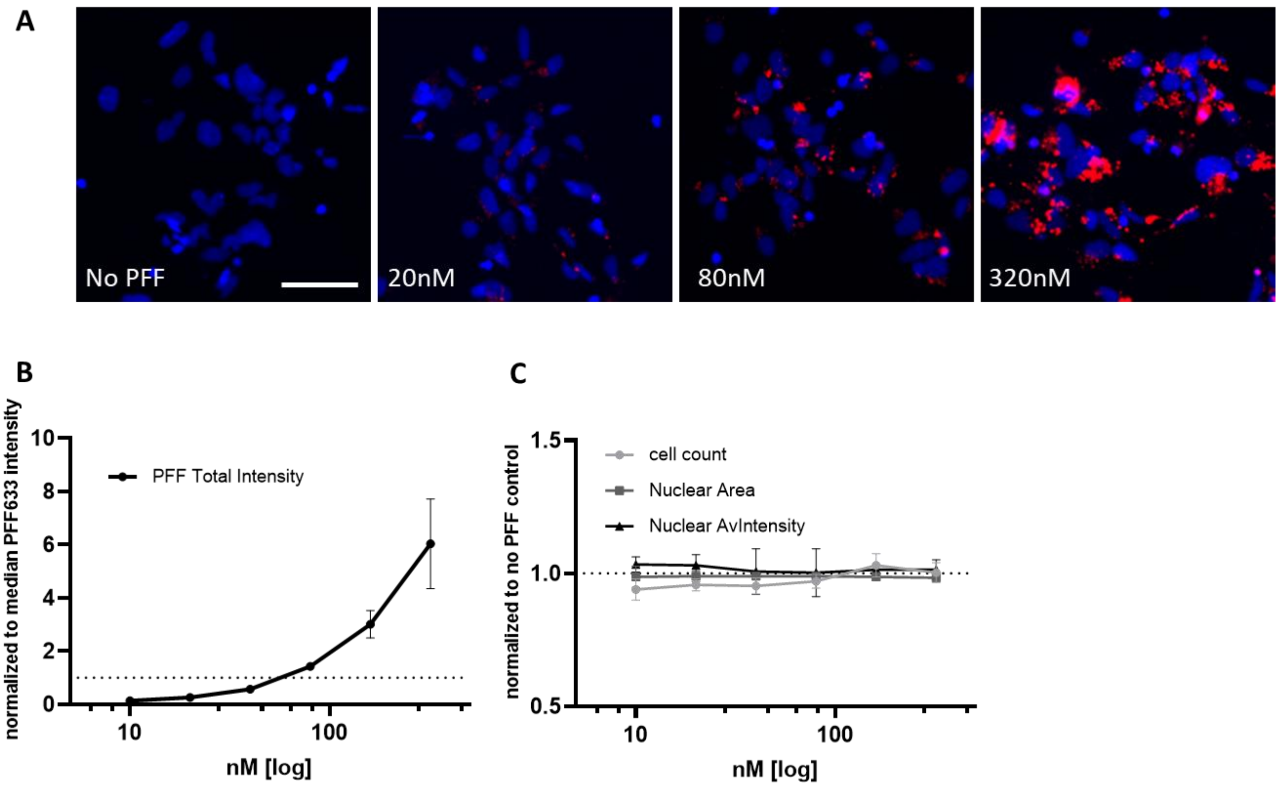


Supplementary Figure S4: Dose dependence of 24-hour Alexa633-fluorphore-labeled PFF uptake in DPCs. A) PFF uptake, overlay of Hoechst (blue) and PFF (red) channels. Size bar: 50µm. B) Quantification of cellular PFF load, normalized to median total intensity of PFF. C) Quantification of cell count, nuclear size and Hoechst intensity as indicators of cell health. There is no significant effect on cell health with increasing PFF concentration.  Graphs represent 3 independent experiments. Error bars: standard deviation.

Supplementary Figure S5:

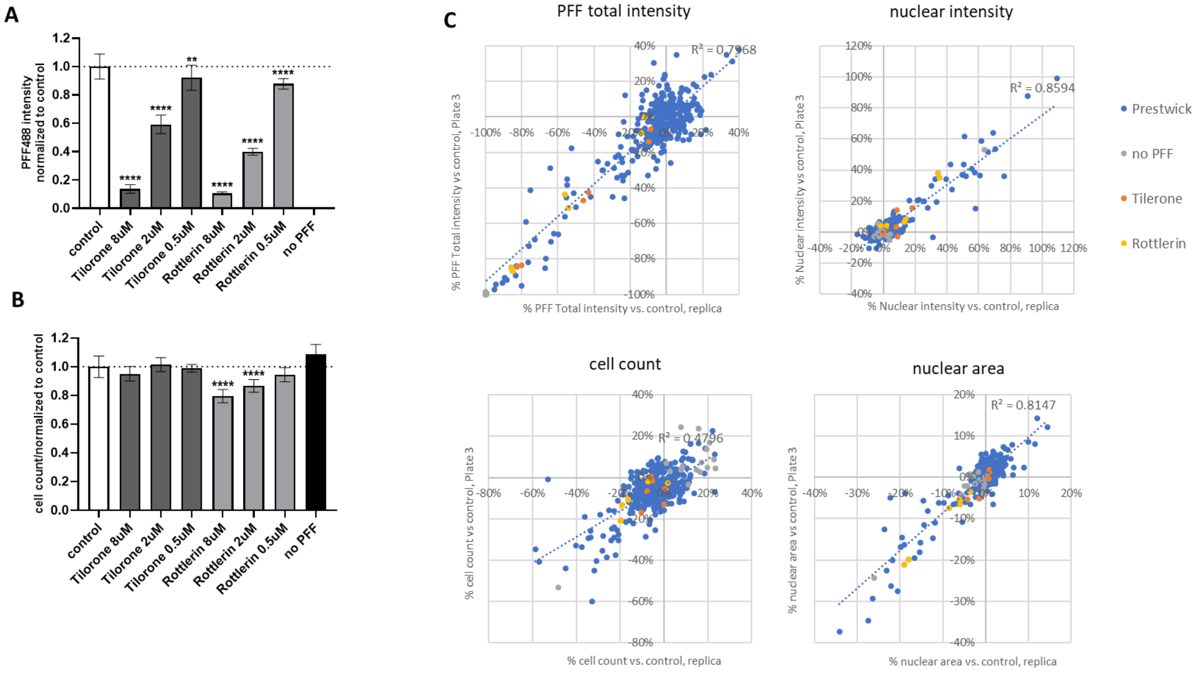


Supplementary Figure S5: PFF uptake screen, controls A) Dose dependent PFF reduction by compounds Rottlerin and Tilorone. B) Cell count/ well. Rottlerin reduces cell count at higher doses. A and B) Values are normalized to mean of vehicle (DMSO) only control. Error bars show standard deviation of values pooled from 5 screening plates. P-values: **** <0.0001, **<0.01, one-way ANOVA and Dunnett's corrected multiple comparisons test.

C) Alignment of replica plates for PFF intensity and cell health parameters. Values are normalized against the mean of PFF only controls. Each data point represents the identical well in the screening and replica plate. Linear regression line (blue line) and R-squared values are included in each graph.

Supplementary Figure S6:

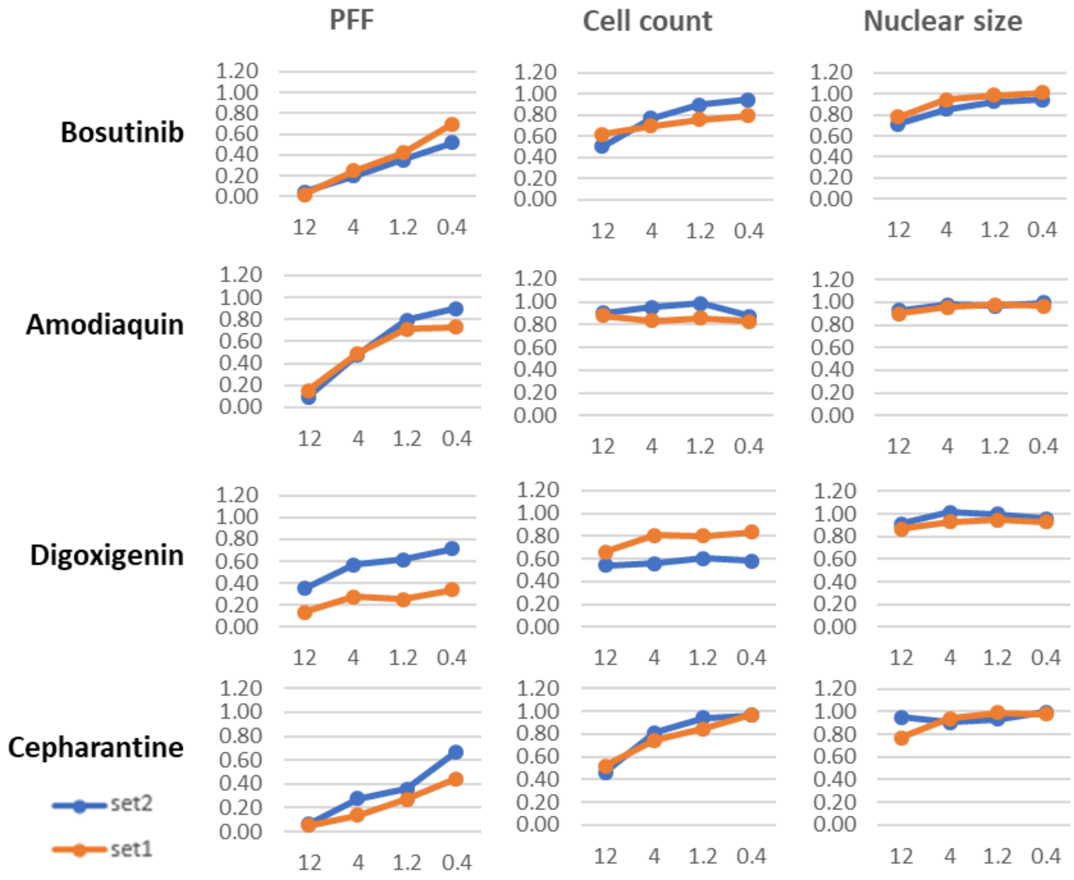


Supplementary Fig. S6: Dose response of active compounds. PFF reduction and cell health parameters. y-axis: change normalized against mean of vehicle-only control (mean control=1). x-axis: compound dose in µM. Each graph shows results from 2 independent experiments (set 1 (orange) and set 2 (blue), 2 replicas per set).
