## Supplemental Table 1 for "A high-content imaging workflow to screen for molecules that reduce cellular uptake of α-synuclein preformed fibrils"

| | | | | | % change relative to mean of control | | | | |
|---|---|---|---|---|---|---|---|---|---|
| plate # | # | row | col | Chemical name | change Mean Nuclear Area | change Mean Nuclear Intensity | change mean PFF_Total Intensity | change mean cell count | CAS number |
| Prestwick-P1 | co | A | 1 | DMSO | -9.3% | -23.6% | -20.3% | -4.5% | |
| Prestwick-P1 | co | B | 1 | DMSO | 2.4% | -0.9% | -0.6% | -5.3% | |
| Prestwick-P1 | co | C | 1 | DMSO | -6.6% | -12.8% | -13.9% | -5.4% | |
| Prestwick-P1 | co | D | 1 | DMSO | 4.2% | -0.5% | 14.5% | -4.7% | |
| Prestwick-P1 | co | E | 1 | DMSO | -9.9% | -21.6% | -14.9% | -13.2% | |
| Prestwick-P1 | co | F | 1 | DMSO | 3.6% | 2.1% | 12.8% | -6.9% | |
| Prestwick-P1 | co | G | 1 | DMSO | 2.5% | 2.1% | -7.9% | -8.0% | |
| Prestwick-P1 | co | H | 1 | DMSO | 5.5% | 13.5% | 0.5% | -6.2% | |
| Prestwick-P1 | co | I | 1 | DMSO | 4.2% | 5.4% | -8.6% | -4.4% | |
| Prestwick-P1 | co | J | 1 | DMSO | 5.0% | 14.4% | 15.1% | -1.7% | |
| Prestwick-P1 | co | K | 1 | DMSO | 1.5% | -9.8% | -10.9% | -1.5% | |
| Prestwick-P1 | co | L | 1 | DMSO | 4.6% | 20.5% | 3.1% | 5.2% | |
| Prestwick-P1 | co | M | 1 | DMSO | 2.8% | -2.1% | 2.8% | 1.3% | |
| Prestwick-P1 | co | N | 1 | DMSO | 4.3% | 17.6% | 26.7% | -1.5% | |
| Prestwick-P1 | co | O | 1 | DMSO | 6.9% | 26.3% | -4.7% | -5.0% | |
| Prestwick-P1 | co | P | 1 | DMSO | 4.1% | 38.7% | 3.2% | 18.7% | |
| Prestwick-P1 | co | A | 2 | DMSO | -15.4% | -21.7% | -12.1% | -4.7% | |
| Prestwick-P1 | co | B | 2 | DMSO | -1.1% | -13.8% | 9.4% | 0.3% | |
| Prestwick-P1 | co | C | 2 | DMSO | -5.4% | -7.1% | -13.9% | 1.8% | |
| Prestwick-P1 | co | D | 2 | DMSO | 1.8% | 2.2% | 2.9% | -3.1% | |
| Prestwick-P1 | co | E | 2 | DMSO | -3.0% | -8.2% | -14.0% | -0.2% | |
| Prestwick-P1 | co | F | 2 | DMSO | 2.5% | 2.8% | 3.9% | -8.8% | |
| Prestwick-P1 | co | G | 2 | DMSO | 1.2% | -8.0% | -9.8% | -5.2% | |
| Prestwick-P1 | co | H | 2 | DMSO | 3.5% | 1.6% | 1.2% | -13.9% | |
| Prestwick-P1 | co | I | 2 | Til2 | -6.7% | 15.1% | -81.4% | 4.6% | |
| Prestwick-P1 | co | J | 2 | Til2 | 1.4% | 36.6% | -69.3% | -2.1% | |
| Prestwick-P1 | co | K | 2 | Til0.5 | -1.3% | -8.9% | -37.9% | 2.3% | |
| Prestwick-P1 | co | L | 2 | Til0.5 | 5.9% | 18.9% | -25.5% | 1.0% | |
| Prestwick-P1 | co | M | 2 | Til0.13 | 1.5% | 1.8% | 5.0% | -6.4% | |
| Prestwick-P1 | co | N | 2 | Til0.13 | 4.9% | 15.5% | 17.6% | -4.2% | |
| Prestwick-P1 | co | O | 2 | DMSO | 3.8% | 11.9% | 10.7% | 0.0% | |
| Prestwick-P1 | co | P | 2 | DMSO | 7.9% | 40.5% | 11.7% | 5.2% | |
| Prestwick-P1 | 1 | A | 3 | Azaguanine-8 | -15.1% | -20.1% | -43.0% | -10.1% | 134-58-7 |
| Prestwick-P1 | 2 | B | 3 | Allantoin | 1.9% | -9.3% | 12.5% | 6.5% | 97-59-6 |
| Prestwick-P1 | 3 | C | 3 | Meticrane | -7.4% | -13.8% | -15.7% | -7.4% | 1084-65-7 |
| Prestwick-P1 | 4 | D | 3 | Benzonatate | 2.0% | -6.6% | 13.5% | 5.8% | 104-31-4 |
| Prestwick-P1 | 5 | E | 3 | Sulfaphenazole | 0.3% | 1.1% | -18.0% | -7.8% | 526-08-9 |
| Prestwick-P1 | 6 | F | 3 | (D)-Panthenol | 4.2% | 3.5% | 0.2% | -3.5% | 81-13-0 |
| Prestwick-P1 | 7 | G | 3 | Chloramphenicol | -4.4% | -13.2% | -23.4% | 6.5% | 56-75-7 |
| Prestwick-P1 | 8 | H | 3 | Epirizole | 3.7% | 8.6% | 8.0% | 0.0% | 18694-40-1 |
| Prestwick-P1 | 9 | I | 3 | Procaine hydrochloride | 6.2% | -0.5% | 40.9% | 1.8% | 51-05-8 |
| Prestwick-P1 | 10 | J | 3 | Moxisylyte hydrochoride | 7.2% | 10.9% | 23.7% | 7.2% | 964-52-3 |
| Prestwick-P1 | 11 | K | 3 | Mupirocin | -0.4% | -4.5% | -3.8% | 0.7% | 12650-69-0 |
| Prestwick-P1 | 12 | L | 3 | Carbamazepine | 8.7% | 15.1% | 16.3% | 1.0% | 298-46-4 |
| Prestwick-P1 | 13 | M | 3 | Morantel tartrate | 1.8% | -2.5% | 11.6% | -1.9% | 26155-31-7 |
| Prestwick-P1 | 14 | N | 3 | (R,S) Homatropine hydrobromide | 7.1% | 13.4% | 62.4% | -7.9% | 51-56-9 |
| Prestwick-P1 | 15 | O | 3 | Todralazine hydrochloride | 7.0% | 2.8% | 23.6% | -7.8% | 3778-76-5 |
| Prestwick-P1 | 16 | P | 3 | Imipramine hydrochloride | 13.4% | 24.3% | 101.1% | -1.4% | 113-52-0 |
| Prestwick-P1 | 17 | A | 4 | Acetazolamide | -10.0% | -18.8% | -26.0% | -13.2% | 59-66-5 |
| Prestwick-P1 | 18 | B | 4 | Metformin hydrochloride | -0.4% | -11.0% | 12.8% | 5.2% | 1115-70-4 |
| Prestwick-P1 | 19 | C | 4 | Hydroflumethiazide | -5.3% | -19.5% | -15.9% | -4.9% | 135-09-1 |
| Prestwick-P1 | 20 | D | 4 | Sulfacetamide sodic hydrate | 1.8% | -0.9% | 13.3% | -4.8% | 6209-17-2 |
| Prestwick-P1 | 21 | E | 4 | Sulfadiazine | -6.5% | -14.9% | -13.9% | -5.3% | 68-35-9 |
| Prestwick-P1 | 22 | F | 4 | Norethynodrel | 0.9% | -0.5% | -15.1% | -11.7% | 68-23-5 |
| Prestwick-P1 | 23 | G | 4 | Diprophylline | -2.7% | -4.7% | -17.1% | -6.1% | 479-18-5 |
| Prestwick-P1 | 24 | H | 4 | Triamterene | 2.2% | 4.1% | 10.5% | -2.1% | 396-01-0 |
| Prestwick-P1 | 25 | I | 4 | Betazole hydrochloride | 0.2% | -8.5% | -2.7% | -11.2% | 138-92-1 |
| Prestwick-P1 | 26 | J | 4 | Isoxicam | 4.9% | 5.1% | 15.2% | 9.3% | 34552-84-6 |
| Prestwick-P1 | 27 | K | 4 | Triflupromazine hydrochloride | -3.9% | 39.5% | -39.3% | -17.8% | 1098-60-8 |
| Prestwick-P1 | 28 | L | 4 | Mefenamic acid | 7.4% | 13.6% | 20.5% | 5.5% | 61-68-7 |
| Prestwick-P1 | 29 | M | 4 | Nifedipine | 4.7% | 4.1% | -7.2% | -24.5% | 21829-25-4 |
| Prestwick-P1 | 30 | N | 4 | Chlorpromazine hydrochloride | 7.6% | 23.8% | 5.7% | -13.5% | 69-09-0 |
| Prestwick-P1 | 31 | O | 4 | Sulindac | 1.4% | 1.8% | 4.4% | -6.7% | 38194-50-2 |
| Prestwick-P1 | 32 | P | 4 | Amitryptiline hydrochloride | 8.8% | 22.8% | 52.7% | -2.7% | 549-18-8 |
| Prestwick-P1 | 33 | A | 5 | Atracurium besylate | -9.0% | -24.2% | -12.7% | -13.8% | 64228-81-5 |
| Prestwick-P1 | 34 | B | 5 | Isoflupredone acetate | 0.5% | -9.8% | 15.0% | 14.9% | 338-98-7 |

| | | | | | | | | | |
|---|---|---|---|---|---|---|---|---|---|
| Prestwick-P1 | 35 | C | 5 | Heptaminol hydrochloride | -6.7% | -16.4% | -18.6% | -13.5% | 543-15-7 |
| Prestwick-P1 | 36 | D | 5 | Sulfathiazole | 2.4% | -1.1% | 2.8% | -6.7% | 72-14-0 |
| Prestwick-P1 | 37 | E | 5 | Thiamphenicol | -5.0% | -13.4% | -27.9% | -3.5% | 15318-45-3 |
| Prestwick-P1 | 38 | F | 5 | Cimetidine | 2.0% | 1.5% | -0.2% | 3.6% | 51481-61-9 |
| Prestwick-P1 | 39 | G | 5 | Dapsone | -2.5% | -8.4% | -21.4% | -4.3% | 80-08-0 |
| Prestwick-P1 | 40 | H | 5 | Troleandomycin | 2.5% | 4.5% | 5.0% | 3.7% | 2751-09-9 |
| Prestwick-P1 | 41 | I | 5 | (S)-Naproxen | -4.7% | -6.7% | -9.9% | -14.2% | 22204-53-1 |
| Prestwick-P1 | 42 | J | 5 | Naphazoline hydrochloride | 3.7% | 6.0% | 8.5% | -2.3% | 550-99-2 |
| Prestwick-P1 | 43 | K | 5 | Acetohexamide | 0.6% | -0.7% | -8.9% | -8.4% | 968-81-0 |
| Prestwick-P1 | 44 | L | 5 | Sulpiride | 6.5% | 13.2% | 11.7% | 1.2% | 15676-16-1 |
| Prestwick-P1 | 45 | M | 5 | Diphenhydramine hydrochloride | 6.2% | -0.5% | 6.1% | -15.0% | 147-24-0 |
| Prestwick-P1 | 46 | N | 5 | Minaprine dihydrochloride | 4.1% | 14.4% | 22.5% | 5.6% | 25953-17-7 |
| Prestwick-P1 | 47 | O | 5 | Adiphenine hydrochloride | 7.3% | 11.1% | 34.3% | -9.2% | 50-42-0 |
| Prestwick-P1 | 48 | P | 5 | Dibucaine | 6.8% | 21.7% | 20.0% | -3.8% | 85-79-0 |
| Prestwick-P1 | 49 | A | 6 | Amiloride hydrochloride dihydrate | -11.0% | -13.2% | -15.9% | -10.4% | 17440-83-4 |
| Prestwick-P1 | 50 | B | 6 | Amprolium hydrochloride | -4.1% | -8.8% | -7.3% | 16.4% | 137-88-2 |
| Prestwick-P1 | 51 | C | 6 | Levodopa | -5.4% | -12.6% | -22.3% | -14.7% | 59-92-7 |
| Prestwick-P1 | 52 | D | 6 | Idoxuridine | 2.9% | -3.0% | 18.9% | -2.5% | 54-42-2 |
| Prestwick-P1 | 53 | E | 6 | Doxylamine succinate | -4.5% | -15.1% | -14.1% | -10.7% | 562-10-7 |
| Prestwick-P1 | 54 | F | 6 | Ethambutol dihydrochloride | 2.4% | 3.8% | 1.4% | -4.5% | 1070-11-7 |
| Prestwick-P1 | 55 | G | 6 | Pyrimethamine | -1.6% | -4.8% | -25.7% | -7.1% | 58-14-0 |
| Prestwick-P1 | 56 | H | 6 | Hexamethonium dibromide dihydrate | 5.2% | 4.8% | 15.1% | -12.9% | 55-97-0 |
| Prestwick-P1 | 57 | I | 6 | Ticlopidine hydrochloride | -2.8% | -4.3% | -10.5% | -15.8% | 53885-35-1 |
| Prestwick-P1 | 58 | J | 6 | Dicyclomine hydrochloride | 0.4% | 44.8% | -11.4% | -28.2% | 67-92-5 |
| Prestwick-P1 | 59 | K | 6 | Benoxinate hydrochloride | 0.5% | -8.1% | -0.8% | -7.0% | 5987-82-6 |
| Prestwick-P1 | 60 | L | 6 | Oxethazaine | 8.1% | 27.8% | 26.0% | -3.2% | 126-27-2 |
| Prestwick-P1 | 61 | M | 6 | Miconazole | -0.5% | -8.5% | 13.0% | -16.3% | 22916-47-8 |
| Prestwick-P1 | 62 | N | 6 | Isoxsuprine hydrochloride | -17.3% | 114.2% | -62.9% | -27.7% | 579-56-6 |
| Prestwick-P1 | 63 | O | 6 | Prednisone | 3.9% | 10.5% | -0.2% | -8.7% | 53-03-2 |
| Prestwick-P1 | 64 | P | 6 | Thioridazine hydrochloride | 3.5% | 22.4% | 20.7% | 10.9% | 130-61-0 |
| Prestwick-P1 | 65 | A | 7 | Hydrochlorothiazide | -13.5% | -23.4% | -10.7% | -15.4% | 58-93-5 |
| Prestwick-P1 | 66 | B | 7 | Sulfaguanidine | -2.8% | -13.5% | -3.8% | 8.0% | 57-67-0 |
| Prestwick-P1 | 67 | C | 7 | Captopril | -8.7% | -14.4% | -10.2% | -11.2% | 62571-86-2 |
| Prestwick-P1 | 68 | D | 7 | Minoxidil | 2.6% | 7.0% | 1.2% | -4.7% | 38304-91-5 |
| Prestwick-P1 | 69 | E | 7 | Antipyrine | 0.4% | -9.7% | -19.4% | -13.5% | 60-80-0 |
| Prestwick-P1 | 70 | F | 7 | Antipyrine, 4-hydroxy | 1.7% | 4.5% | 3.0% | 13.1% | 1672-63-5 |
| Prestwick-P1 | 71 | G | 7 | Diflunisal | -1.9% | -7.2% | -11.8% | -11.7% | 22494-42-4 |
| Prestwick-P1 | 72 | H | 7 | Niclosamide | -12.0% | 23.6% | -84.3% | -18.6% | 50-65-7 |
| Prestwick-P1 | 73 | I | 7 | Amyleine hydrochloride | -13.6% | 68.5% | -40.6% | -36.9% | 532-59-2 |
| Prestwick-P1 | 74 | J | 7 | Lidocaine hydrochloride | 6.5% | 12.5% | 10.2% | 5.6% | 73-78-9 |
| Prestwick-P1 | 75 | K | 7 | Pheniramine maleate | -3.2% | 1.5% | -20.5% | -7.6% | 132-20-7 |
| Prestwick-P1 | 76 | L | 7 | Tolazoline hydrochloride | 6.8% | 13.3% | 14.5% | 0.4% | 59-97-2 |
| Prestwick-P1 | 77 | M | 7 | Acebutolol hydrochloride | 4.1% | -3.3% | 16.6% | -8.1% | 34381-68-5 |
| Prestwick-P1 | 78 | N | 7 | Tolnaftate | 8.8% | 13.2% | 5.3% | -6.6% | 2398-96-1 |
| Prestwick-P1 | 79 | O | 7 | Diphemanil methylsulfate | 5.0% | 48.8% | -66.0% | -18.3% | 62-97-5 |
| Prestwick-P1 | 80 | P | 7 | Trimethobenzamide hydrochloride | 7.4% | 25.0% | 16.0% | -9.2% | 554-92-7 |
| Prestwick-P1 | 81 | A | 8 | Metronidazole | -15.0% | -28.1% | -19.0% | -23.6% | 443-48-1 |
| Prestwick-P1 | 82 | B | 8 | Fulvestrant | -0.3% | 3.1% | -16.7% | 10.0% | 129453-61-8 |
| Prestwick-P1 | 83 | C | 8 | Khellin | -5.1% | -5.1% | -22.6% | -0.6% | 82-02-0 |
| Prestwick-P1 | 84 | D | 8 | Zimelidine dihydrochloride monohydrate | 0.6% | -0.7% | 12.9% | -4.6% | 61129-30-4 |
| Prestwick-P1 | 85 | E | 8 | R(-) Apomorphine hydrochloride hemihydrate | 0.2% | -7.6% | -21.0% | -18.6% | 41372-20-7 |
| Prestwick-P1 | 86 | F | 8 | Amoxapine | 1.0% | 5.5% | 13.3% | 2.8% | 14028-44-5 |
| Prestwick-P1 | 87 | G | 8 | Naloxone hydrochloride | 2.4% | -8.3% | -0.2% | -2.9% | 357-08-4 |
| Prestwick-P1 | 88 | H | 8 | Metolazone | 3.1% | 3.1% | 6.6% | -1.7% | 17560-51-9 |
| Prestwick-P1 | 89 | I | 8 | Bromocryptine mesylate | -1.6% | 1.4% | -30.4% | -15.8% | 22260-51-1 |
| Prestwick-P1 | 90 | J | 8 | Amfepramone hydrochloride | 4.8% | 8.0% | 11.5% | -0.2% | 90-84-6 |
| Prestwick-P1 | 91 | K | 8 | Glipizide | 2.9% | -8.3% | 2.7% | -0.2% | 29094-61-9 |
| Prestwick-P1 | 92 | L | 8 | Loxapine succinate | 11.6% | 5.5% | 52.6% | 3.2% | 27833-64-3 |
| Prestwick-P1 | 93 | M | 8 | Verapamil hydrochloride | 1.2% | -4.8% | -7.2% | -14.2% | 152-11-4 |
| Prestwick-P1 | 94 | N | 8 | Dipyridamole | 11.8% | 11.4% | 6.9% | -10.0% | 58-32-2 |
| Prestwick-P1 | 95 | O | 8 | Erythromycin | 6.6% | 3.4% | 5.2% | -15.5% | 114-07-8 |
| Prestwick-P1 | 96 | P | 8 | Chloroxine | -24.7% | 147.4% | -77.2% | -36.9% | 773-76-2 |
| Prestwick-P1 | 97 | A | 9 | Edrophonium chloride | -12.2% | -23.5% | -9.3% | 2.5% | 116-38-1 |
| Prestwick-P1 | 98 | B | 9 | Moroxidine hydrochloride | -4.5% | -13.9% | 2.7% | 5.2% | 3160-91-6 |
| Prestwick-P1 | 99 | C | 9 | Azacyclonol | -8.1% | -11.4% | -16.1% | -4.3% | 115-46-8 |
| Prestwick-P1 | 100 | D | 9 | Azathioprine | -2.4% | -6.0% | 3.7% | -3.7% | 446-86-6 |
| Prestwick-P1 | 101 | E | 9 | Cyproheptadine hydrochloride | 6.1% | 10.3% | 25.6% | -4.7% | 969-33-5 |
| Prestwick-P1 | 102 | F | 9 | Famotidine | 1.3% | 0.7% | -5.2% | -5.8% | 76824-35-6 |
| Prestwick-P1 | 103 | G | 9 | Ciprofloxacin hydrochloride monohydrate | -1.0% | -3.2% | -14.2% | 0.0% | 93107-08-5 |
| Prestwick-P1 | 104 | H | 9 | Ampicillin trihydrate | 1.4% | 0.1% | 5.3% | 0.3% | 7177-48-2 |
| Prestwick-P1 | 105 | I | 9 | Dehydrocholic acid | -0.8% | -5.7% | -1.4% | -5.5% | 81-23-2 |
| Prestwick-P1 | 106 | J | 9 | Tioconazole | -16.1% | 93.7% | -53.4% | -26.2% | 65899-73-2 |

| | | | | | | | | | |
|---|---|---|---|---|---|---|---|---|---|
| Prestwick-P1 | 107 | K | 9 | Hydroxyzine dihydrochloride | 5.9% | 17.4% | -10.5% | 10.9% | 2192-20-3 |
| Prestwick-P1 | 108 | L | 9 | Diltiazem hydrochloride | 8.3% | 13.6% | 17.4% | 0.0% | 33286-22-5 |
| Prestwick-P1 | 109 | M | 9 | Chlorhexidine | -5.6% | 45.2% | -14.3% | -3.3% | 55-56-1 |
| Prestwick-P1 | 110 | N | 9 | Loperamide hydrochloride | 4.9% | 19.6% | -13.9% | -5.9% | 34552-83-5 |
| Prestwick-P1 | 111 | O | 9 | Didanosine | 2.1% | -1.9% | 8.3% | 0.0% | 69655-05-6 |
| Prestwick-P1 | 112 | P | 9 | Josamycin | 4.8% | 12.1% | -2.9% | -9.1% | 16846-24-5 |
| Prestwick-P1 | 113 | A | 10 | Baclofen | -13.8% | -20.3% | -13.2% | 3.8% | 1134-47-0 |
| Prestwick-P1 | 114 | B | 10 | Acyclovir | -6.9% | -12.8% | -5.4% | 6.5% | 59277-89-3 |
| Prestwick-P1 | 115 | C | 10 | Lynestrenol | -24.9% | 131.4% | -85.6% | -16.4% | 52-76-6 |
| Prestwick-P1 | 116 | D | 10 | Guanabenz acetate | -1.5% | 1.2% | -3.3% | -6.6% | 23256-50-0 |
| Prestwick-P1 | 117 | E | 10 | Danazol | -1.4% | 33.4% | -44.0% | -14.3% | 17230-88-5 |
| Prestwick-P1 | 118 | F | 10 | Nicorandil | 3.7% | 1.2% | -5.0% | 2.7% | 65141-46-0 |
| Prestwick-P1 | 119 | G | 10 | Haloperidol | -3.5% | 2.1% | -15.0% | 2.2% | 52-86-8 |
| Prestwick-P1 | 120 | H | 10 | Naltrexone hydrochloride dihydrate | 0.7% | 1.4% | 4.5% | -2.9% | 16676-29-2 |
| Prestwick-P1 | 121 | I | 10 | Perphenazine | -3.9% | 8.8% | -47.9% | -4.8% | 58-39-9 |
| Prestwick-P1 | 122 | J | 10 | Mefloquine hydrochloride | -7.5% | 62.6% | -64.8% | -32.2% | 51773-92-3 |
| Prestwick-P1 | 123 | K | 10 | Azilsartan medoxomil | 4.4% | 10.8% | -0.4% | 2.6% | 863031-21-4 |
| Prestwick-P1 | 124 | L | 10 | Astemizole | -10.9% | 45.0% | -89.3% | -31.1% | 68844-77-9 |
| Prestwick-P1 | 125 | M | 10 | Chlortetracycline hydrochloride | -3.4% | 3.9% | 2.1% | 6.4% | 64-72-2 |
| Prestwick-P1 | 126 | N | 10 | Tamoxifen citrate | -15.7% | 110.1% | -69.7% | -31.1% | 54965-24-1 |
| Prestwick-P1 | 127 | O | 10 | Paclitaxel | 8.7% | 2.2% | -20.5% | -6.5% | 33069-62-4 |
| Prestwick-P1 | 128 | P | 10 | Ivermectin | -25.4% | 125.7% | -95.9% | -66.1% | 70288-86-7 |
| Prestwick-P1 | 129 | A | 11 | Diazoxide | -12.9% | -19.1% | -19.2% | -7.8% | 364-98-7 |
| Prestwick-P1 | 130 | B | 11 | Amidopyrine | -7.5% | -13.7% | -1.0% | 16.3% | 58-15-1 |
| Prestwick-P1 | 131 | C | 11 | Disulfiram | -27.5% | 50.1% | -98.0% | -32.6% | 97-77-8 |
| Prestwick-P1 | 132 | D | 11 | Acetylsalicylsalicylic acid | -1.3% | -2.0% | -0.4% | -11.8% | 530-75-6 |
| Prestwick-P1 | 133 | E | 11 | Pioglitazone | -1.4% | -4.6% | -11.3% | -8.8% | 111025-46-8 |
| Prestwick-P1 | 134 | F | 11 | Nomifensine maleate | 0.7% | 1.3% | 0.3% | 5.5% | 32795-47-4 |
| Prestwick-P1 | 135 | G | 11 | Chlorpheniramine maleate | -2.2% | -8.8% | -3.6% | -4.3% | 113-92-8 |
| Prestwick-P1 | 136 | H | 11 | Nalbuphine hydrochloride | 1.5% | -0.2% | -3.7% | -2.1% | 23277-43-2 |
| Prestwick-P1 | 137 | I | 11 | Isoconazole | -20.1% | 100.1% | -78.9% | -24.9% | 27523-40-6 |
| Prestwick-P1 | 138 | J | 11 | Spironolactone | 0.5% | -4.8% | 14.6% | -2.8% | 52-01-7 |
| Prestwick-P1 | 139 | K | 11 | Clindamycin hydrochloride | 2.6% | 2.3% | -3.3% | 3.7% | 21462-39-5 |
| Prestwick-P1 | 140 | L | 11 | Terfenadine | -24.4% | 145.3% | -93.3% | -33.1% | 50679-08-8 |
| Prestwick-P1 | 141 | M | 11 | Nicergoline | -3.1% | -5.1% | -17.0% | -4.5% | [illegible]7848-84-6 |
| Prestwick-P1 | 142 | N | 11 | Canrenoic acid potassium salt | 3.7% | 6.5% | 17.5% | 1.7% | 2181-04-6 |
| Prestwick-P1 | 143 | O | 11 | Gallamine triethiodide | -2.4% | -9.8% | 7.0% | 2.4% | 65-29-2 |
| Prestwick-P1 | 144 | P | 11 | Neomycin sulfate | 2.8% | 12.4% | 10.1% | -1.2% | 1405-10-3 |
| Prestwick-P1 | 145 | A | 12 | Busulfan | -9.9% | -21.4% | -13.0% | -2.9% | 55-98-1 |
| Prestwick-P1 | 146 | B | 12 | Pindolol | -4.4% | -12.9% | -2.3% | 16.1% | 13523-86-9 |
| Prestwick-P1 | 147 | C | 12 | Mianserine hydrochloride | -5.0% | -5.1% | 3.4% | -2.4% | 21535-47-7 |
| Prestwick-P1 | 148 | D | 12 | Nocodazole | 9.2% | 15.3% | 7.9% | -24.3% | 31430-18-9 |
| Prestwick-P1 | 149 | E | 12 | Aliskiren hemifumarate | -2.4% | -11.3% | -14.1% | 6.8% | 173334-57-1 |
| Prestwick-P1 | 150 | F | 12 | Oxandrolone | -1.1% | -6.3% | -4.0% | -6.4% | 53-39-4 |
| Prestwick-P1 | 151 | G | 12 | Picotamide monohydrate | -2.8% | -7.8% | -8.4% | -3.0% | 80530-63-8 |
| Prestwick-P1 | 152 | H | 12 | Triamcinolone | -2.4% | 0.9% | 0.1% | 7.0% | 124-94-7 |
| Prestwick-P1 | 153 | I | 12 | Pirenzepine dihydrochloride | -4.9% | -6.7% | -10.4% | -3.4% | 29868-97-1 |
| Prestwick-P1 | 154 | J | 12 | Dexamethasone acetate | 0.2% | 1.3% | 11.8% | -2.8% | 1177-87-3 |
| Prestwick-P1 | 155 | K | 12 | Cefotaxime sodium salt | 1.9% | 2.6% | 4.2% | 4.1% | 64485-93-4 |
| Prestwick-P1 | 156 | L | 12 | Tetracycline hydrochloride | 0.4% | 4.8% | 9.3% | -2.5% | 64-75-5 |
| Prestwick-P1 | 157 | M | 12 | Thioproperazine dimesylate | 1.0% | 6.0% | -37.7% | -6.3% | 2347-80-0 |
| Prestwick-P1 | 158 | N | 12 | Dihydroergotamine tartrate | 4.1% | 9.9% | 7.9% | -2.2% | 5989-77-5 |
| Prestwick-P1 | 159 | O | 12 | Dihydrostreptomycin sulfate | -0.1% | -6.8% | 12.7% | -8.4% | 5490-27-7 |
| Prestwick-P1 | 160 | P | 12 | Gentamicine sulfate | 4.5% | 16.4% | 18.9% | -3.3% | 1405-41-0 |
| Prestwick-P1 | 161 | A | 13 | Isoniazid | -11.0% | -19.8% | -13.6% | 5.7% | 54-85-3 |
| Prestwick-P1 | 162 | B | 13 | Pentylenetetrazole | -7.7% | -19.6% | 0.0% | 16.1% | 54-95-5 |
| Prestwick-P1 | 163 | C | 13 | Tranexamic acid | -9.5% | -18.7% | -6.9% | -2.6% | 1197-18-8 |
| Prestwick-P1 | 164 | D | 13 | Etofylline | -3.1% | -10.2% | 8.2% | -0.2% | 519-37-9 |
| Prestwick-P1 | 165 | E | 13 | Ampyrone | -5.1% | -7.3% | -11.3% | 1.2% | 83-07-8 |
| Prestwick-P1 | 166 | F | 13 | Levamisole hydrochloride | 1.2% | 0.7% | 47.9% | -2.4% | 16595-80-5 |
| Prestwick-P1 | 167 | G | 13 | Midodrine hydrochloride | -1.2% | -8.2% | -5.8% | 1.2% | 3092-17-9 |
| Prestwick-P1 | 168 | H | 13 | Thalidomide | 0.6% | 0.3% | 0.9% | 2.7% | 50-35-1 |
| Prestwick-P1 | 169 | I | 13 | Ranitidine hydrochloride | -3.1% | -0.7% | -7.6% | -9.7% | 66357-59-3 |
| Prestwick-P1 | 170 | J | 13 | Tiratricol, 3,3',5-triiodothyroacetic acid | -0.2% | 1.1% | 2.7% | -5.0% | 51-24-1 |
| Prestwick-P1 | 171 | K | 13 | Piroxicam | 6.4% | 17.6% | 1.9% | -1.3% | 36322-90-4 |
| Prestwick-P1 | 172 | L | 13 | Pyrantel tartrate | 2.8% | 1.2% | 9.8% | 0.7% | 33401-94-4 |
| Prestwick-P1 | 173 | M | 13 | Norfloxacin | -1.9% | -2.4% | 5.0% | 0.7% | 70458-96-7 |
| Prestwick-P1 | 174 | N | 13 | Antimycin A | -1.0% | 23.3% | 4.6% | -17.7% | 1397-94-0 |
| Prestwick-P1 | 175 | O | 13 | Etodolac | -0.3% | -2.8% | 4.6% | -10.6% | 41340-25-4 |
| Prestwick-P1 | 176 | P | 13 | Scopolamin-N-oxide hydrobromide | 3.0% | 11.5% | 11.8% | -5.3% | 6106-81-6 |
| Prestwick-P1 | 177 | A | 14 | Chlorzoxazone | -22.3% | 4.6% | -88.2% | -18.5% | 95-25-0 |
| Prestwick-P1 | 178 | B | 14 | Ornidazole | -7.3% | -23.7% | -6.7% | 9.2% | 16773-42-5 |

| | | | | | | | | | |
|---|---|---|---|---|---|---|---|---|---|
| Prestwick-P1 | 179 | C | 14 | Tranylcypromine hydrochloride | -4.8% | -8.1% | -7.3% | 1.0% | 1986-47-6 |
| Prestwick-P1 | 180 | D | 14 | Alverine citrate salt | -4.1% | 5.2% | -1.8% | 4.3% | 5560-59-8 |
| Prestwick-P1 | 181 | E | 14 | Pargyline hydrochloride | -5.0% | -7.8% | -15.8% | -3.1% | 306-07-0 |
| Prestwick-P1 | 182 | F | 14 | Methocarbamol | -2.6% | -8.9% | 1.7% | 9.2% | 532-03-6 |
| Prestwick-P1 | 183 | G | 14 | Oxolinic acid | -2.5% | -9.7% | -6.7% | -2.1% | 14698-29-4 |
| Prestwick-P1 | 184 | H | 14 | Nimesulide | -1.8% | -7.1% | 4.9% | 1.3% | 51803-78-2 |
| Prestwick-P1 | 185 | I | 14 | Flufenamic acid | -1.7% | -9.6% | -13.5% | -8.7% | 530-78-9 |
| Prestwick-P1 | 186 | J | 14 | Flumequine | -0.9% | -0.7% | 4.2% | -11.6% | 42835-25-6 |
| Prestwick-P1 | 187 | K | 14 | Fenspiride hydrochloride | 3.5% | 6.9% | -2.2% | -0.5% | 5053-08-7 |
| Prestwick-P1 | 188 | L | 14 | Gemfibrozil | 1.2% | 3.4% | 16.6% | 1.2% | 25812-30-0 |
| Prestwick-P1 | 189 | M | 14 | Xylometazoline hydrochloride | 0.6% | 1.4% | 24.0% | 0.1% | 1218-35-5 |
| Prestwick-P1 | 190 | N | 14 | Oxymetazoline hydrochloride | 3.2% | 9.3% | 23.2% | 0.5% | 2315-02-8 |
| Prestwick-P1 | 191 | O | 14 | (L) Hyoscyamine | 1.8% | -5.1% | -4.5% | -8.8% | 101-31-5 |
| Prestwick-P1 | 192 | P | 14 | Chlorphensin carbamate | 0.2% | 7.2% | 5.7% | -4.4% | 886-74-8 |
| Prestwick-P1 | 193 | A | 15 | Ethosuximide | -12.5% | -14.8% | -11.0% | 9.8% | 77-67-8 |
| Prestwick-P1 | 194 | B | 15 | Mafenide hydrochloride | -8.5% | -17.2% | -2.0% | 9.7% | 138-37-4 |
| Prestwick-P1 | 195 | C | 15 | Aceclofenac | -7.3% | -18.8% | -11.1% | 2.0% | 89796-99-6 |
| Prestwick-P1 | 196 | D | 15 | Iproniazide phosphate | -3.0% | -1.6% | -4.6% | 6.8% | 305-33-9 |
| Prestwick-P1 | 197 | E | 15 | Aztreonam | -2.2% | -10.6% | -6.5% | 13.2% | 78110-38-0 |
| Prestwick-P1 | 198 | F | 15 | Cloxacillin sodium salt | -2.1% | -11.8% | -7.1% | 6.8% | 642-78-4 |
| Prestwick-P1 | 199 | G | 15 | Asenapine maleate | 1.7% | 6.3% | 20.6% | -5.3% | 85650-56-2 |
| Prestwick-P1 | 200 | H | 15 | Pentoxifylline | 0.7% | 2.8% | 2.6% | 1.8% | 6493-05-6 |
| Prestwick-P1 | 201 | I | 15 | Tolfenamic acid | -3.7% | -9.0% | -11.0% | -2.4% | 13710-19-5 |
| Prestwick-P1 | 202 | J | 15 | Meclofenamic acid sodium salt monohydrate | 2.1% | -1.1% | 6.6% | -0.7% | 6385-02-0 |
| Prestwick-P1 | 203 | K | 15 | Mefexamide hydrochloride | 3.1% | 11.6% | 2.7% | 6.7% | 3413-64-7 |
| Prestwick-P1 | 204 | L | 15 | Tiapride hydrochloride | 1.2% | -1.1% | 6.7% | 6.4% | 51012-33-0 |
| Prestwick-P1 | 205 | M | 15 | Nifenazone | -0.7% | 1.3% | 16.0% | 3.6% | 2139-47-1 |
| Prestwick-P1 | 206 | N | 15 | Griseofulvin | 2.6% | 8.0% | 18.2% | 0.9% | 126-07-8 |
| Prestwick-P1 | 207 | O | 15 | Carmofur | -0.4% | -3.3% | -1.6% | -3.1% | 61422-45-5 |
| Prestwick-P1 | 208 | P | 15 | Dilazep dihydrochloride | 1.8% | 17.1% | -1.1% | -12.3% | 20153-98-4 |
| Prestwick-P1 | 209 | A | 16 | Riluzole hydrochloride | -12.1% | -22.0% | -12.1% | -3.6% | 850608-87-6 |
| Prestwick-P1 | 210 | B | 16 | Nitrofurantoin | -10.5% | -14.3% | 0.8% | 8.0% | 67-20-9 |
| Prestwick-P1 | 211 | C | 16 | Sulfamethoxazole | -4.9% | -7.7% | -4.1% | 5.5% | 723-46-6 |
| Prestwick-P1 | 212 | D | 16 | Mephenesin | -4.5% | -1.4% | -2.2% | 6.4% | 59-47-2 |
| Prestwick-P1 | 213 | E | 16 | Lacosamide | -3.6% | -13.4% | -14.8% | -1.4% | 175481-36-4 |
| Prestwick-P1 | 214 | F | 16 | Pentolinium bitartrate | -2.6% | -8.0% | -7.9% | 3.3% | 52-62-0 |
| Prestwick-P1 | 215 | G | 16 | Metaraminol bitartrate | -0.8% | -9.8% | -3.9% | 2.7% | 33402-03-8 |
| Prestwick-P1 | 216 | H | 16 | Salbutamol | -0.5% | -3.4% | 4.5% | 3.3% | 18559-94-9 |
| Prestwick-P1 | 217 | I | 16 | Tibolone | -1.1% | -0.8% | -3.7% | 5.8% | 5630-53-5 |
| Prestwick-P1 | 218 | J | 16 | Trimethoprim | -2.1% | -5.0% | -0.3% | -7.3% | 738-70-5 |
| Prestwick-P1 | 219 | K | 16 | Mebendazole | 10.7% | 24.4% | 7.3% | -18.0% | 31431-39-7 |
| Prestwick-P1 | 220 | L | 16 | Fenbufen | 1.5% | -2.0% | 12.9% | -1.8% | 36330-85-5 |
| Prestwick-P1 | 221 | M | 16 | Clemizole hydrochloride | 0.1% | 14.7% | -11.9% | -7.6% | 1163-36-6 |
| Prestwick-P1 | 222 | N | 16 | Tropicamide | -1.0% | 0.5% | 17.2% | -0.3% | 1508-75-4 |
| Prestwick-P1 | 223 | O | 16 | Ofloxacin | 3.1% | -5.0% | 10.7% | 1.0% | 82419-36-1 |
| Prestwick-P1 | 224 | P | 16 | Lomefloxacin hydrochloride | 1.5% | 17.2% | 5.7% | 1.8% | 98079-52-8 |
| Prestwick-P1 | 225 | A | 17 | Hydralazine hydrochloride | -13.0% | -19.3% | -17.8% | -2.0% | 304-20-1 |
| Prestwick-P1 | 226 | B | 17 | Phenelzine sulfate | -8.7% | -16.7% | -12.7% | 7.2% | 156-51-4 |
| Prestwick-P1 | 227 | C | 17 | Phenformin hydrochloride | -2.0% | -13.0% | -6.2% | 12.8% | 834-28-6 |
| Prestwick-P1 | 228 | D | 17 | Flutamide | -4.8% | -7.1% | -10.1% | 3.0% | 13311-84-7 |
| Prestwick-P1 | 229 | E | 17 | Aminopurine, 6-benzyl | -1.7% | -9.4% | -9.0% | 9.2% | 1214-39-7 |
| Prestwick-P1 | 230 | F | 17 | Tolbutamide | 0.8% | -7.7% | -6.2% | 0.9% | 64-77-7 |
| Prestwick-P1 | 231 | G | 17 | Prilocaine hydrochloride | -2.4% | -5.5% | -6.2% | 0.7% | 1786-81-8 |
| Prestwick-P1 | 232 | H | 17 | (S,+) Camptothecine | -4.8% | 43.6% | -30.2% | -56.5% | 7689-03-4 |
| Prestwick-P1 | 233 | I | 17 | Metoclopramide monohydrochloride | -0.5% | -3.9% | 4.1% | 4.4% | 7232-21-5 |
| Prestwick-P1 | 234 | J | 17 | Fenbendazole | -12.6% | 79.8% | -60.6% | -51.9% | 43210-67-9 |
| Prestwick-P1 | 235 | K | 17 | Ketoprofen | 6.4% | 5.9% | 4.0% | 8.3% | 22071-15-4 |
| Prestwick-P1 | 236 | L | 17 | Indapamide | 5.5% | -0.6% | 7.5% | -6.0% | 26807-65-8 |
| Prestwick-P1 | 237 | M | 17 | Nefopam hydrochloride | 3.6% | -1.0% | 6.4% | -6.4% | 23327-57-3 |
| Prestwick-P1 | 238 | N | 17 | Phentolamine hydrochloride | 4.2% | -2.5% | 15.8% | -3.2% | 73-05-2 |
| Prestwick-P1 | 239 | O | 17 | Orphenadrine hydrochloride | 8.1% | -0.5% | 21.8% | -4.4% | 341-69-5 |
| Prestwick-P1 | 240 | P | 17 | Proglumide | 1.0% | 13.1% | 12.2% | 4.7% | 6620-60-6 |
| Prestwick-P1 | 241 | A | 18 | Mexiletine hydrochloride | -12.6% | -19.0% | -14.6% | 12.5% | 5370-01-4 |
| Prestwick-P1 | 242 | B | 18 | Flavoxate hydrochloride | -10.5% | -17.4% | -20.2% | 12.4% | 3717-88-2 |
| Prestwick-P1 | 243 | C | 18 | Chlorothiazide | -3.1% | -14.0% | -8.9% | -1.1% | 58-94-6 |
| Prestwick-P1 | 244 | D | 18 | Diphenidol hydrochloride | 1.2% | 8.1% | -8.0% | -7.1% | 3254-89-5 |
| Prestwick-P1 | 245 | E | 18 | Ethisterone | 0.1% | -2.0% | -12.2% | 3.8% | 434-03-7 |
| Prestwick-P1 | 246 | F | 18 | Triprolidine hydrochloride | 3.1% | 1.1% | 24.8% | 11.7% | 550-70-9 |
| Prestwick-P1 | 247 | G | 18 | Vincamine | 5.7% | -0.3% | 57.3% | 10.9% | 1617-90-9 |
| Prestwick-P1 | 248 | H | 18 | Indomethacin | -0.8% | -1.2% | 3.8% | 1.0% | 53-86-1 |
| Prestwick-P1 | 249 | I | 18 | Fludrocortisone acetate | -4.4% | -5.2% | 3.2% | 3.2% | 514-36-3 |
| Prestwick-P1 | 250 | J | 18 | Fenoterol hydrobromide | 1.6% | -6.5% | 17.9% | -3.5% | 1944-12-3 |

| | | | | | | | | | |
|---|---|---|---|---|---|---|---|---|---|
| Prestwick-P1 | 251 | K | 18 | Dantrolene sodium salt | 4.0% | 4.1% | 13.8% | 7.2% | 14663-23-1 |
| Prestwick-P1 | 252 | L | 18 | Trazodone hydrochloride | 5.3% | -0.9% | 19.1% | 2.0% | 25332-39-2 |
| Prestwick-P1 | 253 | M | 18 | Lisinopril | 1.0% | -4.6% | 0.0% | 4.9% | 83915-83-7 |
| Prestwick-P1 | 254 | N | 18 | Lincomycin hydrochloride | 5.4% | -1.2% | 13.1% | 1.0% | 859-18-7 |
| Prestwick-P1 | 255 | O | 18 | Ifenprodil tartrate | 5.1% | 15.2% | 3.1% | -5.3% | 23210-58-4 |
| Prestwick-P1 | 256 | P | 18 | Flunarizine dihydrochloride | -7.9% | 52.4% | -31.2% | -44.6% | 30484-77-6 |
| Prestwick-P1 | 257 | A | 19 | Bufexamac | -10.9% | -17.7% | -19.0% | 7.8% | 2438-72-4 |
| Prestwick-P1 | 258 | B | 19 | Glutethimide, para-amino | -9.1% | -15.4% | -9.4% | 4.0% | 125-84-8 |
| Prestwick-P1 | 259 | C | 19 | Norethindrone | -4.7% | -9.2% | -13.9% | 12.7% | 68-22-4 |
| Prestwick-P1 | 260 | D | 19 | Nortriptyline hydrochloride | -3.1% | -1.7% | 4.8% | -1.8% | 894-71-3 |
| Prestwick-P1 | 261 | E | 19 | Doxepin hydrochloride | -1.6% | 1.6% | 8.9% | 13.0% | 1229-29-4 |
| Prestwick-P1 | 262 | F | 19 | Dyclonine hydrochloride | -2.1% | 3.0% | 5.5% | 1.1% | 536-43-6 |
| Prestwick-P1 | 263 | G | 19 | Cortisone | 0.2% | -5.5% | -6.7% | 6.9% | 53-06-5 |
| Prestwick-P1 | 264 | H | 19 | Prednisolone | 0.2% | 0.2% | -3.9% | 4.9% | 50-24-8 |
| Prestwick-P1 | 265 | I | 19 | Homochlorcyclizine dihydrochloride | 0.7% | 11.6% | 5.8% | 3.7% | 1982-36-1 |
| Prestwick-P1 | 266 | J | 19 | Diethylcarbamazine citrate | 0.1% | -9.1% | 3.9% | -0.8% | 1642-54-2 |
| Prestwick-P1 | 267 | K | 19 | Glafenine hydrochloride | 6.1% | 3.4% | -11.2% | 8.3% | 65513-72-6 |
| Prestwick-P1 | 268 | L | 19 | Pimethixene maleate | 13.3% | 5.2% | 87.6% | 14.9% | 13187-06-9 |
| Prestwick-P1 | 269 | M | 19 | Telenzepine dihydrochloride | 12.8% | 30.7% | 14.7% | -4.8% | 147416-96-4 |
| Prestwick-P1 | 270 | N | 19 | Econazole nitrate | -10.6% | 38.7% | -16.0% | -21.9% | 24169-02-6 |
| Prestwick-P1 | 271 | O | 19 | Trifluoperazine dihydrochloride | -3.6% | 48.1% | -54.6% | -16.6% | 440-17-5 |
| Prestwick-P1 | 272 | P | 19 | Enalapril maleate | -0.3% | 6.3% | 13.1% | -4.7% | 76095-16-4 |
| Prestwick-P1 | 273 | A | 20 | Dropropizine | -12.6% | -21.5% | -14.1% | 21.4% | 17692-31-8 |
| Prestwick-P1 | 274 | B | 20 | Pinacidil monohydrate | -9.2% | -16.2% | -5.3% | 22.5% | 85371-64-8 |
| Prestwick-P1 | 275 | C | 20 | Niflumic acid | -4.7% | -9.6% | -12.1% | 7.6% | 4394-00-7 |
| Prestwick-P1 | 276 | D | 20 | Isotretinoin | -1.1% | -4.4% | -3.9% | 0.8% | 4759-48-2 |
| Prestwick-P1 | 277 | E | 20 | Dimenhydrinate | -0.6% | -5.3% | 10.7% | 7.9% | 523-87-5 |
| Prestwick-P1 | 278 | F | 20 | Disopyramide | -3.3% | -4.1% | 7.5% | 10.3% | 3737-09-5 |
| Prestwick-P1 | 279 | G | 20 | Fenofibrate | -3.7% | 2.0% | -9.5% | -3.8% | 49562-28-9 |
| Prestwick-P1 | 280 | H | 20 | Bumetanide | -3.5% | -4.6% | -11.9% | -11.5% | 28395-03-1 |
| Prestwick-P1 | 281 | I | 20 | Chenodiol | 0.2% | 1.1% | -5.8% | -2.4% | 474-25-9 |
| Prestwick-P1 | 282 | J | 20 | Perhexiline maleate | -15.3% | 86.1% | -92.2% | -20.7% | 6724-53-4 |
| Prestwick-P1 | 283 | K | 20 | Pergolide mesylate | 5.6% | 9.7% | 8.1% | 1.4% | 66104-23-2 |
| Prestwick-P1 | 284 | L | 20 | Acemetacin | 6.8% | 7.7% | 15.4% | 1.2% | 53164-05-9 |
| Prestwick-P1 | 285 | M | 20 | Bupivacaine hydrochloride | 1.6% | -3.2% | 4.0% | -3.2% | 18010-40-7 |
| Prestwick-P1 | 286 | N | 20 | Clemastine fumarate | 3.4% | 10.1% | -25.5% | -2.5% | 14976-57-9 |
| Prestwick-P1 | 287 | O | 20 | Minocycline hydrochloride | 0.5% | -0.6% | 15.6% | -1.4% | 13614-98-7 |
| Prestwick-P1 | 288 | P | 20 | Glibenclamide | 0.3% | 8.0% | 12.7% | -2.4% | 10238-21-8 |
| Prestwick-P1 | 289 | A | 21 | Albendazole | -13.4% | 21.4% | -33.3% | -21.2% | 54965-21-8 |
| Prestwick-P1 | 290 | B | 21 | Clonidine hydrochloride | -7.0% | -21.5% | 0.3% | 11.0% | 4205-91-8 |
| Prestwick-P1 | 291 | C | 21 | Retinoic acid | -1.0% | -3.8% | -20.2% | -3.8% | 302-79-4 |
| Prestwick-P1 | 292 | D | 21 | Antazoline hydrochloride | -4.9% | -5.8% | 8.3% | -0.8% | 2508-72-7 |
| Prestwick-P1 | 293 | E | 21 | Clotrimazole | -1.1% | 18.8% | -33.2% | -6.5% | 23593-75-1 |
| Prestwick-P1 | 294 | F | 21 | Vinpocetine | -0.6% | 35.0% | 16.2% | -22.4% | 42971-09-5 |
| Prestwick-P1 | 295 | G | 21 | Labetalol hydrochloride | -1.1% | -5.4% | 1.2% | 8.6% | 32780-64-6 |
| Prestwick-P1 | 296 | H | 21 | Cinnarizine | -8.3% | 60.5% | -27.0% | -50.9% | 298-57-7 |
| Prestwick-P1 | 297 | I | 21 | Oxybutynin chloride | -5.2% | 34.4% | -20.9% | -10.7% | 1508-65-2 |
| Prestwick-P1 | 298 | J | 21 | Spiperone | -0.8% | -4.7% | 15.1% | -2.1% | 749-02-0 |
| Prestwick-P1 | 299 | K | 21 | Benzydamine hydrochloride | 9.5% | 18.1% | 32.3% | 8.6% | 132-69-4 |
| Prestwick-P1 | 300 | L | 21 | Fipexide hydrochloride | 6.5% | 5.9% | -4.7% | 4.3% | 34161-23-4 |
| Prestwick-P1 | 301 | M | 21 | Oxytetracycline dihydrate | 1.1% | -1.8% | 19.2% | 7.0% | 6153-64-6 |
| Prestwick-P1 | 302 | N | 21 | Pimozide | -7.2% | 38.7% | -59.2% | -17.5% | 2062-78-4 |
| Prestwick-P1 | 303 | O | 21 | Guanethidine sulfate | 5.5% | 1.9% | 9.7% | -2.1% | 60-02-6 |
| Prestwick-P1 | 304 | P | 21 | Quinacrine dihydrochloride hydrate | -9.0% | 32.6% | -90.6% | -24.9% | 69-05-6 |
| Prestwick-P1 | co | A | 22 | DMSO | -10.8% | -19.0% | -11.6% | 2.7% | |
| Prestwick-P1 | co | B | 22 | DMSO | -9.6% | -21.1% | -4.9% | 3.4% | |
| Prestwick-P1 | co | C | 22 | Rott2 | -13.0% | 59.6% | -84.5% | -25.8% | |
| Prestwick-P1 | co | D | 22 | Rott2 | -18.9% | 48.5% | -84.2% | -27.8% | |
| Prestwick-P1 | co | E | 22 | Rott0.5 | -10.6% | 9.3% | -51.8% | -17.4% | |
| Prestwick-P1 | co | F | 22 | Rott0.5 | -11.2% | 8.9% | -49.4% | -13.5% | |
| Prestwick-P1 | co | G | 22 | Rott0.13 | -6.6% | -4.2% | -15.9% | -2.8% | |
| Prestwick-P1 | co | H | 22 | Rott0.13 | -9.4% | -1.7% | -10.3% | -5.6% | |
| Prestwick-P1 | co | I | 22 | DMSO | 0.7% | -1.3% | -2.9% | 0.1% | |
| Prestwick-P1 | co | J | 22 | DMSO | 0.8% | -7.0% | -1.5% | 11.0% | |
| Prestwick-P1 | co | K | 22 | DMSO | 5.3% | 9.5% | -3.6% | 16.8% | |
| Prestwick-P1 | co | L | 22 | DMSO | 6.9% | 10.9% | 8.0% | 10.2% | |
| Prestwick-P1 | co | M | 22 | DMSO | -0.3% | -3.2% | 6.6% | 2.7% | |
| Prestwick-P1 | co | N | 22 | DMSO | 1.6% | -1.4% | 8.5% | -3.9% | |
| Prestwick-P1 | co | O | 22 | DMSO | 2.7% | -0.8% | 12.1% | -7.7% | |
| Prestwick-P1 | co | P | 22 | DMSO | 2.7% | 7.4% | 4.9% | -2.2% | |
| Prestwick-P1 | co | A | 23 | DMSO | -12.2% | -22.1% | 0.9% | 13.1% | |
| Prestwick-P1 | co | B | 23 | DMSO | -10.2% | -23.0% | 6.8% | 9.9% | |

| | | | | | | | | | |
|---|---|---|---|---|---|---|---|---|---|
| Prestwick-P1 | co | C | 23 | DMSO | -1.1% | -7.0% | 1.1% | -0.6% | |
| Prestwick-P1 | co | D | 23 | DMSO | -3.7% | -13.5% | -1.9% | 3.0% | |
| Prestwick-P1 | co | E | 23 | DMSO | -2.5% | -3.0% | -2.2% | 3.6% | |
| Prestwick-P1 | co | F | 23 | DMSO | -2.2% | -11.9% | -2.1% | 0.9% | |
| Prestwick-P1 | co | G | 23 | DMSO | -2.3% | 1.7% | -9.0% | 3.1% | |
| Prestwick-P1 | co | H | 23 | DMSO | -0.2% | 4.0% | 2.2% | -8.5% | |
| Prestwick-P1 | co | I | 23 | DMSO | 0.0% | 2.9% | -6.3% | 5.9% | |
| Prestwick-P1 | co | J | 23 | DMSO | 0.1% | -5.3% | -8.6% | 5.0% | |
| Prestwick-P1 | co | K | 23 | DMSO | 5.0% | 15.2% | 2.7% | 4.2% | |
| Prestwick-P1 | co | L | 23 | DMSO | 5.4% | 13.3% | -7.1% | 6.5% | |
| Prestwick-P1 | co | M | 23 | DMSO | 0.7% | -2.6% | -0.5% | 1.2% | |
| Prestwick-P1 | co | N | 23 | DMSO | 1.3% | 0.7% | 5.9% | -1.5% | |
| Prestwick-P1 | co | O | 23 | DMSO | 2.7% | 3.8% | 9.2% | -14.2% | |
| Prestwick-P1 | co | P | 23 | DMSO | -2.1% | 3.2% | 6.2% | 7.2% | |
| Prestwick-P1 | co | A | 24 | Ceph4 | -22.6% | 109.9% | -99.8% | -44.8% | |
| Prestwick-P1 | co | B | 24 | noPFF | -5.8% | -14.4% | -99.8% | 1.9% | |
| Prestwick-P1 | co | C | 24 | noPFF | -1.1% | -1.8% | -99.9% | 4.3% | |
| Prestwick-P1 | co | D | 24 | noPFF | -1.0% | -8.2% | -97.2% | 11.8% | |
| Prestwick-P1 | co | E | 24 | noPFF | -1.1% | -4.5% | -100.0% | 21.0% | |
| Prestwick-P1 | co | F | 24 | noPFF | 0.1% | -4.7% | -100.0% | -2.0% | |
| Prestwick-P1 | co | G | 24 | noPFF | 0.2% | 1.2% | -100.0% | 6.5% | |
| Prestwick-P1 | co | H | 24 | noPFF | 0.0% | -0.6% | -99.8% | -0.1% | |
| Prestwick-P1 | co | I | 24 | noPFF | -0.3% | 3.5% | -99.9% | 1.5% | |
| Prestwick-P1 | co | J | 24 | noPFF | 1.8% | 1.2% | -99.9% | 5.0% | |
| Prestwick-P1 | co | K | 24 | noPFF | 6.8% | 13.1% | -99.9% | 12.2% | |
| Prestwick-P1 | co | L | 24 | noPFF | 3.9% | 17.5% | -99.3% | 11.4% | |
| Prestwick-P1 | co | M | 24 | noPFF | 1.7% | 5.4% | -100.0% | 14.8% | |
| Prestwick-P1 | co | N | 24 | noPFF | -0.3% | 5.1% | -100.0% | 17.3% | |
| Prestwick-P1 | co | O | 24 | noPFF | 3.2% | 5.2% | -99.9% | 6.6% | |
| Prestwick-P1 | co | P | 24 | noPFF | -0.8% | 11.0% | -99.9% | 7.6% | |
| Prestwick-P2 | co | A | 1 | DMSO | -5.4% | 0.0% | 0.0% | 0.0% | |
| Prestwick-P2 | co | B | 1 | DMSO | 0.9% | -0.4% | -5.8% | -12.8% | |
| Prestwick-P2 | co | C | 1 | DMSO | -1.1% | -11.1% | 5.0% | -11.4% | |
| Prestwick-P2 | co | D | 1 | DMSO | 2.8% | 0.3% | 10.4% | -4.8% | |
| Prestwick-P2 | co | E | 1 | DMSO | -0.6% | -7.0% | -2.7% | 0.8% | |
| Prestwick-P2 | co | F | 1 | DMSO | 2.3% | 0.9% | -0.2% | -7.4% | |
| Prestwick-P2 | co | G | 1 | DMSO | -1.6% | -9.5% | -3.6% | 3.6% | |
| Prestwick-P2 | co | H | 1 | DMSO | 1.7% | 3.3% | 0.2% | -4.8% | |
| Prestwick-P2 | co | I | 1 | DMSO | -2.0% | -3.4% | -16.3% | 4.2% | |
| Prestwick-P2 | co | J | 1 | DMSO | 2.8% | 3.8% | -9.0% | -5.6% | |
| Prestwick-P2 | co | K | 1 | DMSO | -0.4% | 0.0% | -6.1% | 11.1% | |
| Prestwick-P2 | co | L | 1 | DMSO | 2.4% | 8.1% | -1.3% | 8.6% | |
| Prestwick-P2 | co | M | 1 | DMSO | -3.0% | 0.0% | -9.0% | 4.4% | |
| Prestwick-P2 | co | N | 1 | DMSO | 4.3% | 13.8% | 3.4% | -11.5% | |
| Prestwick-P2 | co | O | 1 | DMSO | 1.1% | 11.5% | -1.3% | 4.5% | |
| Prestwick-P2 | co | P | 1 | DMSO | 0.9% | 17.1% | -20.8% | -0.4% | |
| Prestwick-P2 | co | A | 2 | DMSO | -3.6% | -11.0% | 17.6% | -6.4% | |
| Prestwick-P2 | co | B | 2 | DMSO | 0.6% | -0.7% | 8.8% | -0.3% | |
| Prestwick-P2 | co | C | 2 | DMSO | -3.8% | -8.1% | 3.0% | -5.6% | |
| Prestwick-P2 | co | D | 2 | DMSO | -0.6% | 3.7% | -4.6% | -1.5% | |
| Prestwick-P2 | co | E | 2 | DMSO | -1.5% | -7.7% | -2.0% | -5.9% | |
| Prestwick-P2 | co | F | 2 | DMSO | 1.9% | 3.1% | -1.1% | -10.7% | |
| Prestwick-P2 | co | G | 2 | DMSO | -1.3% | -6.6% | -0.4% | -7.2% | |
| Prestwick-P2 | co | H | 2 | DMSO | 1.0% | 3.9% | -3.7% | -5.1% | |
| Prestwick-P2 | co | I | 2 | Til2 | -7.7% | 16.0% | -78.8% | -13.1% | |
| Prestwick-P2 | co | J | 2 | Til2 | -5.2% | 25.6% | -75.1% | -8.2% | |
| Prestwick-P2 | co | K | 2 | Til0.5 | 0.3% | 1.5% | -42.4% | 4.1% | |
| Prestwick-P2 | co | L | 2 | Til0.5 | 1.2% | 9.8% | -29.5% | 5.7% | |
| Prestwick-P2 | co | M | 2 | Til0.13 | -3.8% | 2.0% | -13.7% | 1.2% | |
| Prestwick-P2 | co | N | 2 | Til0.13 | 1.3% | 7.9% | -6.7% | 1.2% | |
| Prestwick-P2 | co | O | 2 | DMSO | 0.8% | 14.9% | 1.9% | 4.1% | |
| Prestwick-P2 | co | P | 2 | DMSO | 0.8% | 15.0% | 4.1% | 6.4% | |
| Prestwick-P2 | 305 | A | 3 | Bupropion hydrochloride | -2.7% | -8.8% | 6.7% | -15.1% | 31677-93-7 |
| Prestwick-P2 | 306 | B | 3 | Alprenolol hydrochloride | 1.0% | -3.7% | 14.2% | 7.6% | 13707-88-5 |
| Prestwick-P2 | 307 | C | 3 | Ethacrynic acid | -1.3% | -7.9% | 5.7% | -15.8% | 58-54-8 |
| Prestwick-P2 | 308 | D | 3 | Praziquantel | 1.2% | 0.6% | 2.4% | 0.3% | 55268-74-1 |
| Prestwick-P2 | 309 | E | 3 | Clomipramine hydrochloride | 1.4% | -3.3% | -16.8% | -11.8% | 17321-77-6 |
| Prestwick-P2 | 310 | F | 3 | Fendiline hydrochloride | -8.8% | 51.8% | -61.5% | -29.2% | 13636-18-5 |
| Prestwick-P2 | 311 | G | 3 | Methylprednisolone, 6-alpha | -1.1% | -1.8% | -2.9% | -5.2% | 83-43-2 |
| Prestwick-P2 | 312 | H | 3 | Quinidine hydrochloride monohydrate | -0.5% | 2.9% | 4.9% | -1.6% | 6151-40-2 |
| Prestwick-P2 | 313 | I | 3 | Pyrilamine maleate | 8.1% | -4.1% | 75.4% | -4.3% | 59-33-6 |
| Prestwick-P2 | 314 | J | 3 | Sulfinpyrazone | 2.8% | 10.2% | -6.5% | -9.2% | 57-96-5 |

| | | | | | | | | | |
|---|---|---|---|---|---|---|---|---|---|
| Prestwick-P2 | 315 | K | 3 | Mifepristone | 3.3% | -3.3% | -12.5% | 0.4% | 84371-65-3 |
| Prestwick-P2 | 316 | L | 3 | Diperodon hydrochloride | 1.6% | 5.5% | 2.4% | 10.5% | 537-12-2 |
| Prestwick-P2 | 317 | M | 3 | Amodiaquin dihydrochloride dihydrate | -4.6% | 4.7% | -64.9% | -9.6% | 6398-98-7 |
| Prestwick-P2 | 318 | N | 3 | Mebeverine hydrochloride | 0.2% | 10.0% | 7.8% | 8.7% | 2753-45-9 |
| Prestwick-P2 | 319 | O | 3 | Clofilium tosylate | -1.1% | 7.0% | -6.0% | -12.4% | 92953-10-1 |
| Prestwick-P2 | 320 | P | 3 | Fluphenazine dihydrochloride | -0.8% | 27.8% | -30.1% | -5.4% | 146-56-5 |
| Prestwick-P2 | 321 | A | 4 | Streptomycin sulfate | -2.6% | -10.7% | -6.1% | -8.2% | 3810-74-0 |
| Prestwick-P2 | 322 | B | 4 | Alfuzosin hydrochloride | 0.3% | -4.6% | -2.7% | -3.0% | 81403-68-1 |
| Prestwick-P2 | 323 | C | 4 | Lopinavir | -1.9% | -6.2% | -22.6% | -9.5% | 192725-17-0 |
| Prestwick-P2 | 324 | D | 4 | Practolol | -0.4% | 0.6% | 5.0% | 2.3% | 6673-35-4 |
| Prestwick-P2 | 325 | E | 4 | Furosemide | -3.5% | -12.0% | -14.8% | -8.2% | 54-31-9 |
| Prestwick-P2 | 326 | F | 4 | Methapyrilene hydrochloride | 4.7% | 5.9% | 12.4% | 3.2% | 135-23-9 |
| Prestwick-P2 | 327 | G | 4 | Chlorthalidone | -1.5% | -7.1% | -14.2% | -0.3% | 77-36-1 |
| Prestwick-P2 | 328 | H | 4 | Dobutamine hydrochloride | 0.0% | 3.2% | -10.5% | 2.6% | 49745-95-1 |
| Prestwick-P2 | 329 | I | 4 | Bambuterol hydrochloride | -0.9% | -6.1% | 2.5% | -7.3% | 81732-46-9 |
| Prestwick-P2 | 330 | J | 4 | Betamethasone | 2.4% | 2.7% | -0.4% | -1.6% | 378-44-9 |
| Prestwick-P2 | 331 | K | 4 | Ketotifen fumarate | 0.4% | -3.6% | -8.2% | 0.2% | 34580-14-8 |
| Prestwick-P2 | 332 | L | 4 | Debrisoquin sulfate | 2.4% | 1.9% | 4.7% | -10.2% | 581-88-4 |
| Prestwick-P2 | 333 | M | 4 | Lidoflazine | -6.9% | 22.5% | -47.2% | -23.6% | 3416-26-0 |
| Prestwick-P2 | 334 | N | 4 | Betaxolol hydrochloride | 3.0% | 6.9% | 8.6% | -1.5% | 63659-19-8 |
| Prestwick-P2 | 335 | O | 4 | Terbutaline hemisulfate | 1.7% | -5.1% | 12.3% | 0.9% | 23031-32-5 |
| Prestwick-P2 | 336 | P | 4 | Ketanserin tartrate hydrate | 1.8% | 7.9% | 12.5% | -4.2% | 83846-83-7 |
| Prestwick-P2 | 337 | A | 5 | Chlorpropamide | -1.5% | -17.1% | -7.4% | -12.4% | 94-20-2 |
| Prestwick-P2 | 338 | B | 5 | Phenylpropanolamine hydrochloride | 0.8% | -8.0% | -4.6% | 5.4% | 154-41-6 |
| Prestwick-P2 | 339 | C | 5 | Zidovudine | -1.4% | -13.4% | -20.5% | -7.5% | 30516-87-1 |
| Prestwick-P2 | 340 | D | 5 | Sulfisoxazole | 1.4% | 1.6% | 2.3% | -2.4% | 127-69-5 |
| Prestwick-P2 | 341 | E | 5 | Desipramine hydrochloride | -1.6% | -8.5% | -5.6% | -6.2% | 58-28-6 |
| Prestwick-P2 | 342 | F | 5 | Clorgyline hydrochloride | 0.7% | 11.9% | -16.8% | -6.2% | 17780-75-5 |
| Prestwick-P2 | 343 | G | 5 | Moclobemide | -0.7% | -5.7% | -8.5% | -6.9% | 71320-77-9 |
| Prestwick-P2 | 344 | H | 5 | Clopamide | 1.3% | 1.8% | 1.6% | 1.9% | 636-54-4 |
| Prestwick-P2 | 345 | I | 5 | Colchicine | 10.5% | 13.6% | 12.1% | -26.4% | 64-86-8 |
| Prestwick-P2 | 346 | J | 5 | Metergoline | 3.3% | -0.6% | -11.1% | -3.0% | 17692-51-2 |
| Prestwick-P2 | 347 | K | 5 | Amethopterin | 1.3% | -1.3% | -9.6% | 2.1% | 60388-53-6 |
| Prestwick-P2 | 348 | L | 5 | Methylergometrine maleate | 3.9% | 2.6% | 13.7% | -2.4% | 57432-61-8 |
| Prestwick-P2 | 349 | M | 5 | Nicardipine hydrochloride | -2.1% | 5.5% | -38.5% | -13.9% | 54527-84-3 |
| Prestwick-P2 | 350 | N | 5 | Probucol | -9.9% | 24.2% | -65.8% | -18.2% | 23288-49-5 |
| Prestwick-P2 | 351 | O | 5 | Hemicholinium bromide | 2.2% | -4.3% | 2.0% | 3.6% | 312-45-8 |
| Prestwick-P2 | 352 | P | 5 | Kanamycin A sulfate | 3.2% | 11.4% | 15.4% | -4.2% | 25389-94-0 |
| Prestwick-P2 | 353 | A | 6 | Ascorbic acid | -5.3% | -13.6% | -10.3% | -1.6% | 50-81-7 |
| Prestwick-P2 | 354 | B | 6 | (L) Methyldopa | -0.7% | -5.1% | -20.8% | 17.2% | 555-30-6 |
| Prestwick-P2 | 355 | C | 6 | Zaprinast | 1.5% | -6.6% | -12.9% | -10.7% | 37762-06-4 |
| Prestwick-P2 | 356 | D | 6 | Chlormezanone | 1.1% | -4.1% | 3.6% | 4.7% | 80-77-3 |
| Prestwick-P2 | 357 | E | 6 | Clenbuterol hydrochloride | -2.6% | -14.9% | -15.3% | -4.9% | 21898-19-1 |
| Prestwick-P2 | 358 | F | 6 | Maprotiline hydrochloride | 1.0% | 0.6% | -2.0% | -0.3% | 10347-81-6 |
| Prestwick-P2 | 359 | G | 6 | Hycanthone | 2.0% | -2.6% | -5.7% | -16.1% | 3105-97-3 |
| Prestwick-P2 | 360 | H | 6 | Adenosine 5'-monophosphate monohydrate | 3.1% | 3.5% | 11.6% | 0.2% | 18422-05-4 |
| Prestwick-P2 | 361 | I | 6 | Brinzolamide | -1.9% | -6.5% | -12.3% | -3.3% | 138890-62-7 |
| Prestwick-P2 | 362 | J | 6 | Ambroxol hydrochloride | 3.0% | 5.6% | -1.0% | 2.6% | 23828-92-4 |
| Prestwick-P2 | 363 | K | 6 | Cefditoren Pivoxil | -0.4% | -0.4% | -10.4% | -3.1% | 117467-28-4 |
| Prestwick-P2 | 364 | L | 6 | Clofazimine | -11.3% | 20.0% | -97.7% | -37.5% | 2030-63-9 |
| Prestwick-P2 | 365 | M | 6 | Mitoxantrone dihydrochloride | -16.5% | -29.1% | -97.6% | -61.6% | 70476-82-3 |
| Prestwick-P2 | 366 | N | 6 | GBR 12909 dihydrochloride | -20.8% | 79.9% | -77.8% | -44.8% | 67469-78-7 |
| Prestwick-P2 | 367 | O | 6 | Amikacin hydrate | 2.8% | 2.1% | -1.7% | -9.4% | 37517-28-5 |
| Prestwick-P2 | 368 | P | 6 | Etoposide | -4.1% | 23.9% | -15.6% | -42.5% | 33419-42-0 |
| Prestwick-P2 | 369 | A | 7 | Cefoperazone dihydrate | -3.0% | -6.7% | -8.8% | -9.5% | 62893-19-0 |
| Prestwick-P2 | 370 | B | 7 | Zoxazolamine | 1.0% | -3.9% | -5.3% | 1.7% | 61-80-3 |
| Prestwick-P2 | 371 | C | 7 | Procainamide hydrochloride | 1.1% | -4.1% | -22.6% | -7.0% | 614-39-1 |
| Prestwick-P2 | 372 | D | 7 | N6-methyladenosine | 1.1% | 4.0% | 0.2% | 0.6% | 1867-73-8 |
| Prestwick-P2 | 373 | E | 7 | Thioguanosine | -0.7% | 1.4% | -34.6% | -16.3% | 85-31-4 |
| Prestwick-P2 | 374 | F | 7 | Chlorprothixene hydrochloride | -2.2% | 20.0% | -24.1% | -14.2% | 6469-93-8 |
| Prestwick-P2 | 375 | G | 7 | Amoxicillin | 1.7% | -1.8% | -15.2% | -3.2% | 26787-78-0 |
| Prestwick-P2 | 376 | H | 7 | Pemirolast potassium | 2.1% | -2.3% | 4.0% | 7.1% | 100299-08-9 |
| Prestwick-P2 | 377 | I | 7 | Benfluorex | -3.8% | 13.9% | -33.9% | -23.2% | 23602-78-0 |
| Prestwick-P2 | 378 | J | 7 | Bepridil hydrochloride | -12.5% | 55.7% | -76.5% | -47.2% | 74764-40-2 |
| Prestwick-P2 | 379 | K | 7 | Nafronyl oxalate | 0.2% | -4.5% | -11.5% | 2.4% | 3200-06-4 |
| Prestwick-P2 | 380 | L | 7 | Bezafibrate | 4.1% | 5.0% | 17.4% | 8.7% | 41859-67-0 |
| Prestwick-P2 | 381 | M | 7 | Carbetapentane citrate | 3.2% | 10.6% | 2.2% | -1.7% | 23142-01-0 |
| Prestwick-P2 | 382 | N | 7 | Dequalinium dichloride | 2.0% | 3.5% | 3.5% | 0.4% | 522-51-0 |
| Prestwick-P2 | 383 | O | 7 | (Z,E) Clomiphene citrate | -8.5% | 55.1% | -80.9% | -38.3% | 50-41-9 |
| Prestwick-P2 | 384 | P | 7 | Oxantel pamoate | 2.7% | 7.3% | -4.3% | -8.3% | 68813-55-8 |
| Prestwick-P2 | 385 | A | 8 | Tacrine hydrochloride | -2.5% | -11.2% | -13.8% | -0.2% | 1684-40-8 |
| Prestwick-P2 | 386 | B | 8 | Bisoprolol hemifumarate | 4.5% | -8.4% | -10.0% | 8.6% | 104344-23-2 |

| | | | | | | | | | |
|---|---|---|---|---|---|---|---|---|---|
| Prestwick-P2 | 387 | C | 8 | Guanfacine hydrochloride | 1.8% | -4.7% | -27.1% | -15.1% | 29110-48-3 |
| Prestwick-P2 | 388 | D | 8 | Domperidone | 9.5% | -3.7% | 61.3% | 9.1% | 57808-66-9 |
| Prestwick-P2 | 389 | E | 8 | Ritodrine hydrochloride | -0.3% | -8.0% | -12.2% | -6.3% | 23239-51-2 |
| Prestwick-P2 | 390 | F | 8 | Clozapine | 1.1% | 5.0% | -1.7% | 0.4% | 5786-21-0 |
| Prestwick-P2 | 391 | G | 8 | Dextromethorphan hydrobromide monohydrate | 3.4% | -3.8% | -6.8% | -4.6% | 6700-34-1 |
| Prestwick-P2 | 392 | H | 8 | Droperidol | 2.4% | 6.4% | 11.8% | 7.5% | 548-73-2 |
| Prestwick-P2 | 393 | I | 8 | Meloxicam | -1.0% | -5.6% | -5.2% | 9.2% | 71125-38-7 |
| Prestwick-P2 | 394 | J | 8 | Benzbromarone | 3.0% | -1.2% | 3.3% | -1.4% | 3562-84-3 |
| Prestwick-P2 | 395 | K | 8 | Nefazodone hydrochloride | 0.6% | 6.3% | -32.4% | -4.5% | 82752-99-6 |
| Prestwick-P2 | 396 | L | 8 | Clebopride maleate | 4.1% | 0.8% | -7.5% | -0.8% | 84370-95-6 |
| Prestwick-P2 | 397 | M | 8 | Ketoconazole | 3.1% | 1.8% | -30.3% | -4.3% | 65277-42-1 |
| Prestwick-P2 | 398 | N | 8 | Fusidic acid sodium salt | 8.3% | -1.7% | 6.1% | -0.9% | 751-94-0 |
| Prestwick-P2 | 399 | O | 8 | Prochlorperazine dimaleate | 5.8% | 12.4% | -44.2% | -4.5% | 84-02-6 |
| Prestwick-P2 | 400 | P | 8 | Hesperidin | 3.1% | 9.7% | -7.0% | -5.9% | 520-26-3 |
| Prestwick-P2 | 401 | A | 9 | Testosterone propionate | -5.9% | 18.4% | -26.2% | -20.0% | 57-85-2 |
| Prestwick-P2 | 402 | B | 9 | Haloprogin | -19.7% | 51.4% | -81.2% | -19.0% | 777-11-7 |
| Prestwick-P2 | 403 | C | 9 | Dienogest | 1.5% | -5.5% | -16.7% | -6.8% | 65928-58-7 |
| Prestwick-P2 | 404 | D | 9 | Amifostine | 1.8% | -0.9% | 1.9% | 4.0% | 20537-88-6 |
| Prestwick-P2 | 405 | E | 9 | Suloctidil | -23.1% | 85.2% | -86.2% | -22.1% | 54063-56-8 |
| Prestwick-P2 | 406 | F | 9 | Zotepine | -3.8% | 46.2% | -21.4% | -18.4% | 26615-21-4 |
| Prestwick-P2 | 407 | G | 9 | Pefloxacine | 1.6% | -4.7% | -1.3% | 0.3% | 70458-92-3 |
| Prestwick-P2 | 408 | H | 9 | Corticosterone | 0.2% | -2.8% | -2.9% | 4.4% | 50-22-6 |
| Prestwick-P2 | 409 | I | 9 | Estradiol-17 beta | 1.7% | -0.3% | -9.8% | 7.9% | 50-28-2 |
| Prestwick-P2 | 410 | J | 9 | Clobutinol hydrochloride | 1.3% | 4.7% | 9.3% | 0.2% | 1215-83-4 |
| Prestwick-P2 | 411 | K | 9 | Alizapride hydrochloride | 1.3% | 5.3% | -5.9% | 6.3% | 59338-87-3 |
| Prestwick-P2 | 412 | L | 9 | Stanozolol | 1.7% | 13.1% | -15.6% | -15.0% | 10418-03-8 |
| Prestwick-P2 | 413 | M | 9 | Tropisetron hydrochloride | 5.4% | -4.0% | 33.6% | 5.6% | 105826-92-4 |
| Prestwick-P2 | 414 | N | 9 | Cefixime | 2.0% | 5.5% | 5.7% | 0.4% | 79350-37-1 |
| Prestwick-P2 | 415 | O | 9 | Vatalanib | -0.2% | 1.2% | -12.4% | -1.9% | 212141-54-3 |
| Prestwick-P2 | 416 | P | 9 | Itopride | 1.4% | 10.2% | -2.4% | -11.2% | 122898-67-3 |
| Prestwick-P2 | 417 | A | 10 | (L) Thyroxine | -4.2% | -7.2% | -4.1% | 4.0% | 51-48-9 |
| Prestwick-P2 | 418 | B | 10 | Idebenone | -1.7% | 3.6% | -11.9% | -2.4% | 58186-27-9 |
| Prestwick-P2 | 419 | C | 10 | Carbarsone | -2.1% | -13.3% | -16.2% | 3.1% | 121-59-5 |
| Prestwick-P2 | 420 | D | 10 | Amlodipine | 0.4% | 9.7% | -26.7% | -8.5% | 88150-42-9 |
| Prestwick-P2 | 421 | E | 10 | Carisoprodol | 1.2% | -7.7% | -5.8% | 6.2% | 78-44-4 |
| Prestwick-P2 | 422 | F | 10 | Cephalosporanic acid, 7-amino | -0.8% | -4.1% | -5.7% | 5.2% | 957-68-6 |
| Prestwick-P2 | 423 | G | 10 | Cyanocobalamin | -2.4% | -12.6% | 3.5% | -0.9% | 68-19-9 |
| Prestwick-P2 | 424 | H | 10 | Cefadroxil | 1.0% | -0.5% | 0.5% | -2.0% | 50370-12-2 |
| Prestwick-P2 | 425 | I | 10 | Nebivolol hydrochloride | -8.2% | 18.0% | -24.2% | -12.4% | 99200-09-6 |
| Prestwick-P2 | 426 | J | 10 | Oxcarbazepine | 2.4% | 6.1% | -4.4% | -7.9% | 28721-07-5 |
| Prestwick-P2 | 427 | K | 10 | Calcipotriene | 2.9% | 0.6% | -15.4% | 0.0% | 112965-21-6 |
| Prestwick-P2 | 428 | L | 10 | Linezolid | 0.5% | 0.0% | 11.0% | 5.5% | 165800-03-3 |
| Prestwick-P2 | 429 | M | 10 | Metrizamide | -0.2% | -6.1% | -3.8% | 3.4% | 31112-62-6 |
| Prestwick-P2 | 430 | N | 10 | Quetiapine hemifumarate | 1.3% | 11.3% | 4.8% | -1.0% | 111974-72-2 |
| Prestwick-P2 | 431 | O | 10 | Cefotetan | 4.0% | 4.7% | -13.2% | -13.4% | 69712-56-7 |
| Prestwick-P2 | 432 | P | 10 | Fentiazac | 1.8% | 7.5% | 14.1% | -5.8% | 18046-21-4 |
| Prestwick-P2 | 433 | A | 11 | Pepstatin A | -5.7% | -8.6% | -9.1% | 13.2% | 26305-03-3 |
| Prestwick-P2 | 434 | B | 11 | Delavirdine | 0.2% | -4.9% | 0.6% | 10.6% | 136817-59-9 |
| Prestwick-P2 | 435 | C | 11 | Modafinil | -3.0% | -12.4% | -15.8% | 3.2% | 68693-11-8 |
| Prestwick-P2 | 436 | D | 11 | Bacampicillin hydrochloride | -3.4% | 1.6% | -8.5% | 14.2% | 37661-08-8 |
| Prestwick-P2 | 437 | E | 11 | Linagliptin | 0.0% | -4.5% | -18.5% | -6.1% | 668270-12-0 |
| Prestwick-P2 | 438 | F | 11 | Buflomedil hydrochloride | 1.6% | 3.4% | -7.1% | -7.0% | 35543-24-9 |
| Prestwick-P2 | 439 | G | 11 | Cyclosporin A | -0.6% | -2.5% | -16.1% | -2.7% | 59865-13-3 |
| Prestwick-P2 | 440 | H | 11 | Digitoxigenin | -5.0% | -4.1% | -67.4% | -14.8% | 143-62-4 |
| Prestwick-P2 | 441 | I | 11 | Cyclobenzaprine hydrochloride | 1.9% | -2.8% | 21.1% | 1.7% | 6202-23-9 |
| Prestwick-P2 | 442 | J | 11 | Carteolol hydrochloride | -0.4% | 0.3% | -1.1% | -2.3% | 51781-21-6 |
| Prestwick-P2 | 443 | K | 11 | Mebhydroline 1,5-naphtalenedisulfonate | 2.1% | -6.5% | 9.1% | 2.7% | 6153-33-9 |
| Prestwick-P2 | 444 | L | 11 | Meclocycline sulfosalicylate | 0.9% | -0.5% | -5.8% | 3.1% | 73816-42-9 |
| Prestwick-P2 | 445 | M | 11 | Tosufloxacin hydrochloride | 2.0% | -8.4% | -9.4% | -2.2% | 100490-36-6 |
| Prestwick-P2 | 446 | N | 11 | Efavirenz | -11.7% | 60.0% | -53.1% | -28.7% | 154598-52-4 |
| Prestwick-P2 | 447 | O | 11 | Brompheniramine maleate | -1.6% | -7.1% | 2.6% | -0.3% | 980-71-2 |
| Prestwick-P2 | 448 | P | 11 | Primaquine diphosphate | 2.0% | 10.4% | -7.8% | -10.0% | 63-45-6 |
| Prestwick-P2 | 449 | A | 12 | Adamantamine fumarate | -4.9% | -12.8% | -7.8% | 8.8% | 80789-67-9 |
| Prestwick-P2 | 450 | B | 12 | Butoconazole nitrate | -6.6% | 18.1% | -24.3% | -10.2% | 32872-77-1 |
| Prestwick-P2 | 451 | C | 12 | Lamivudine | -2.1% | -8.9% | -5.2% | -8.4% | 134678-17-4 |
| Prestwick-P2 | 452 | D | 12 | Biotin | -0.4% | -3.6% | 3.3% | 6.0% | 58-85-5 |
| Prestwick-P2 | 453 | E | 12 | Dibenzepine hydrochloride | -1.6% | -2.9% | 13.2% | 6.3% | 315-80-0 |
| Prestwick-P2 | 454 | F | 12 | Roxatidine acetate hydrochloride | -1.0% | 1.5% | -2.3% | 7.2% | 93793-83-0 |
| Prestwick-P2 | 455 | G | 12 | Digoxin | -10.0% | -9.8% | -78.9% | -23.9% | 20830-75-5 |
| Prestwick-P2 | 456 | H | 12 | Doxorubicin hydrochloride | -54.5% | -55.3% | -73.4% | -87.0% | 25316-40-9 |
| Prestwick-P2 | 457 | I | 12 | Hydrocortisone base | -0.2% | -5.5% | -5.1% | -2.0% | 50-23-7 |
| Prestwick-P2 | 458 | J | 12 | Pitavastatin calcium | -0.6% | 19.1% | -12.7% | -3.8% | 147511-69-1 |

| | | | | | | | | |
|---|---|---|---|---|---|---|---|---|
| Prestwick-P2 | 459 | K | 12 | Meclozine dihydrochloride | -2.9% | 13.7% | -36.5% | -24.4% | 1104-22-9 |
| Prestwick-P2 | 460 | L | 12 | Melatonin | -0.2% | 2.3% | 4.1% | 4.2% | 73-31-4 |
| Prestwick-P2 | 461 | M | 12 | Rifapentine | -0.3% | 3.9% | 5.9% | -4.7% | 61379-65-5 |
| Prestwick-P2 | 462 | N | 12 | Neostigmine bromide | 2.9% | 5.5% | 4.4% | -0.3% | 114-80-7 |
| Prestwick-P2 | 463 | O | 12 | Progesterone | -0.1% | 7.4% | -2.7% | -7.5% | 57-83-0 |
| Prestwick-P2 | 464 | P | 12 | Felodipine | -7.5% | 34.0% | -24.4% | -14.9% | 72509-76-3 |
| Prestwick-P2 | 465 | A | 13 | Amiodarone hydrochloride | -26.0% | 62.1% | -94.6% | -59.9% | 19774-82-4 |
| Prestwick-P2 | 466 | B | 13 | Amphotericin B | -4.1% | 6.8% | -30.6% | -5.3% | 1397-89-3 |
| Prestwick-P2 | 467 | C | 13 | Bisacodyl | -2.4% | -9.0% | -13.5% | -1.5% | 603-50-9 |
| Prestwick-P2 | 468 | D | 13 | Erlotinib | 0.6% | 9.1% | -3.2% | -4.6% | 183321-74-6 |
| Prestwick-P2 | 469 | E | 13 | Valacyclovir hydrochloride | 0.3% | -5.6% | -25.9% | 6.1% | 124832-27-5 |
| Prestwick-P2 | 470 | F | 13 | Cisapride | 3.3% | 1.2% | 26.9% | 2.8% | 81098-60-4 |
| Prestwick-P2 | 471 | G | 13 | Carbimazole | -0.6% | -0.6% | -2.8% | -2.0% | 22232-54-8 |
| Prestwick-P2 | 472 | H | 13 | Epiandrosterone | 2.0% | 5.0% | 0.7% | -7.6% | 481-29-8 |
| Prestwick-P2 | 473 | I | 13 | Pilocarpine nitrate | 1.7% | -9.2% | -11.4% | -14.2% | 148-72-1 |
| Prestwick-P2 | 474 | J | 13 | Dicloxacillin sodium salt hydrate | 0.6% | -1.3% | 5.5% | 4.2% | 13412-64-1 |
| Prestwick-P2 | 475 | K | 13 | Butalbital | 1.2% | -0.5% | 13.6% | 1.1% | 77-26-9 |
| Prestwick-P2 | 476 | L | 13 | Dinoprost trometamol | 0.6% | -1.4% | 7.1% | 3.5% | 38362-01-5 |
| Prestwick-P2 | 477 | M | 13 | Niridazole | -0.6% | -1.7% | 9.7% | 0.0% | 61-57-4 |
| Prestwick-P2 | 478 | N | 13 | Ceforanide | -0.9% | 8.9% | 13.8% | 1.5% | 60925-61-3 |
| Prestwick-P2 | 479 | O | 13 | Raclopride | 2.1% | -0.3% | 2.8% | -5.3% | 84225-95-6 |
| Prestwick-P2 | 480 | P | 13 | Closantel | -5.9% | 26.5% | -26.6% | -16.8% | 57808-65-8 |
| Prestwick-P2 | 481 | A | 13 | Serotonin hydrochloride | -8.6% | -5.8% | -65.0% | -5.6% | 153-98-0 |
| Prestwick-P2 | 482 | B | 14 | Cefotiam hydrochloride | -0.3% | -7.4% | -2.8% | 5.7% | 61622-34-2 |
| Prestwick-P2 | 483 | C | 14 | Metixene hydrochloride | -0.9% | 6.1% | -34.1% | -6.5% | 1553-34-0 |
| Prestwick-P2 | 484 | D | 14 | Nitrofural | -1.5% | -0.9% | -4.5% | 1.2% | 59-87-0 |
| Prestwick-P2 | 485 | E | 14 | Vigabatrin hydrochloride | -1.7% | -8.9% | -2.2% | 9.3% | 1391054-02-6 |
| Prestwick-P2 | 486 | F | 14 | Biperiden hydrochloride | -1.6% | 4.7% | -2.7% | -4.7% | 1235-82-1 |
| Prestwick-P2 | 487 | G | 14 | Fluoxetine hydrochloride | -2.7% | 0.0% | 2.2% | -4.8% | 59333-67-4 |
| Prestwick-P2 | 488 | H | 14 | Iohexol | -0.4% | -2.0% | -7.9% | 3.9% | 66108-95-0 |
| Prestwick-P2 | 489 | I | 14 | Lofexidine | -0.5% | -2.6% | -3.6% | -5.4% | 31036-80-3 |
| Prestwick-P2 | 490 | J | 14 | Thiostrepton | -17.5% | 34.9% | -93.2% | -27.6% | 1393-48-2 |
| Prestwick-P2 | 491 | K | 14 | Pirenperone | 2.1% | -2.9% | 9.6% | -0.6% | 75444-65-4 |
| Prestwick-P2 | 492 | L | 14 | Grepafloxacin | 1.1% | -3.1% | 19.6% | -0.9% | 146863-02-7 |
| Prestwick-P2 | 493 | M | 14 | Ciclopirox ethanolamine | -29.9% | 97.9% | -99.2% | -29.6% | 41621-49-2 |
| Prestwick-P2 | 494 | N | 14 | Probenecid | 1.5% | 5.9% | 15.1% | 4.2% | 57-66-9 |
| Prestwick-P2 | 495 | O | 14 | Hexetidine | -30.6% | 102.7% | -93.5% | -27.0% | 141-94-6 |
| Prestwick-P2 | 496 | P | 14 | Selegiline hydrochloride | -2.8% | 5.5% | 5.6% | -5.3% | 14611-52-0 |
| Prestwick-P2 | 497 | A | 15 | Rofecoxib | -2.7% | -11.6% | -2.2% | 10.4% | 162011-90-7 |
| Prestwick-P2 | 498 | B | 15 | Benperidol | -2.3% | -4.4% | -6.9% | 12.9% | 2062-84-2 |
| Prestwick-P2 | 499 | C | 15 | Omeprazole | -3.2% | -7.5% | -10.8% | 5.0% | 73590-58-6 |
| Prestwick-P2 | 500 | D | 15 | Propylthiouracil | -0.8% | 0.7% | -4.2% | -0.3% | 51-52-5 |
| Prestwick-P2 | 501 | E | 15 | Cetirizine dihydrochloride | 1.0% | -5.1% | -18.0% | 5.3% | 83881-52-1 |
| Prestwick-P2 | 502 | F | 15 | Etifenin | 2.9% | 2.2% | -8.8% | 6.2% | 63245-28-3 |
| Prestwick-P2 | 503 | G | 15 | Norcyclobenzaprine | -1.9% | -2.9% | -4.1% | -0.5% | 303-50-4 |
| Prestwick-P2 | 504 | H | 15 | Pyrazinamide | 0.5% | 4.9% | 8.5% | 0.5% | 98-96-4 |
| Prestwick-P2 | 505 | I | 15 | Miglitol | -1.2% | -6.0% | -9.5% | 4.5% | 72432-03-2 |
| Prestwick-P2 | 506 | J | 15 | Tiabendazole | 0.7% | -0.4% | 10.1% | 0.0% | 148-79-8 |
| Prestwick-P2 | 507 | K | 15 | Phenacetin | 0.6% | 1.1% | 6.9% | 5.8% | 62-44-2 |
| Prestwick-P2 | 508 | L | 15 | Atovaquone | -1.2% | 6.0% | 11.1% | -1.5% | 95233-18-4 |
| Prestwick-P2 | 509 | M | 15 | Betahistine mesylate | 1.2% | -2.6% | 11.3% | -1.2% | 54856-23-4 |
| Prestwick-P2 | 510 | N | 15 | Tobramycin | 1.5% | 6.9% | 17.1% | -3.8% | 32986-56-4 |
| Prestwick-P2 | 511 | O | 15 | Pentamidine isethionate | 1.6% | 5.7% | -0.3% | -2.5% | 140-64-7 |
| Prestwick-P2 | 512 | P | 15 | Tolazamide | -1.5% | 7.3% | 14.7% | 2.1% | 1156-19-0 |
| Prestwick-P2 | 513 | A | 16 | Cefaclor hydrate | -2.1% | -10.3% | -1.8% | 11.3% | 70356-03-5 |
| Prestwick-P2 | 514 | B | 16 | Colistin sulfate | -3.1% | 2.5% | -1.6% | 6.8% | 1264-72-8 |
| Prestwick-P2 | 515 | C | 16 | Terconazole | -1.0% | 11.2% | -27.0% | 7.0% | 67915-31-5 |
| Prestwick-P2 | 516 | D | 16 | Tiaprofenic acid | 0.5% | -8.2% | 101.1% | 2.8% | 33005-95-7 |
| Prestwick-P2 | 517 | E | 16 | Metaproterenol sulfate | -1.9% | -3.3% | -4.4% | 12.9% | 5874-97-5 |
| Prestwick-P2 | 518 | F | 16 | Sisomicin sulfate | 0.4% | -2.0% | -11.1% | -3.4% | 53179-09-2 |
| Prestwick-P2 | 519 | G | 16 | Trimethadione | -0.6% | -6.9% | -4.2% | 11.9% | 127-48-0 |
| Prestwick-P2 | 520 | H | 16 | Lovastatin | 1.8% | -1.7% | 1.8% | 2.1% | 75330-75-5 |
| Prestwick-P2 | 521 | I | 16 | Rifampicin | -1.3% | 0.1% | -10.8% | 13.1% | 13292-46-1 |
| Prestwick-P2 | 522 | J | 16 | Ethionamide | 3.0% | -2.2% | -0.5% | -5.2% | 536-33-4 |
| Prestwick-P2 | 523 | K | 16 | Methoxamine hydrochloride | 1.9% | -0.5% | 17.8% | 2.0% | 61-16-5 |
| Prestwick-P2 | 524 | L | 16 | (S,-) Atenolol | -0.2% | -2.2% | 2.3% | 7.3% | 93379-54-5 |
| Prestwick-P2 | 525 | M | 16 | Tetramisole hydrochloride | -0.2% | -2.4% | 15.0% | 0.0% | 5086-74-8 |
| Prestwick-P2 | 526 | N | 16 | Pregnenolone | -1.3% | 1.0% | -31.8% | -11.3% | 145-13-1 |
| Prestwick-P2 | 527 | O | 16 | Nifuroxazide | 2.2% | 4.9% | 7.9% | -3.4% | 965-52-6 |
| Prestwick-P2 | 528 | P | 16 | Mirtazapine | -0.6% | 6.5% | 6.0% | -4.7% | 61337-67-5 |
| Prestwick-P2 | 529 | A | 17 | Daunorubicin hydrochloride | -45.1% | -60.6% | -81.0% | -72.9% | 23541-50-6 |
| Prestwick-P2 | 530 | B | 17 | Dosulepin hydrochloride | 1.7% | -7.2% | -4.6% | 6.5% | 897-15-4 |

| | | | | | | | | | |
|---|---|---|---|---|---|---|---|---|---|
| Prestwick-P2 | 531 | C | 17 | Vancomycin hydrochloride | 2.2% | -7.7% | -4.1% | 1.9% | 1404-93-9 |
| Prestwick-P2 | 532 | D | 17 | Artemisinin | -0.1% | -6.6% | -7.2% | 8.7% | 63968-64-9 |
| Prestwick-P2 | 533 | E | 17 | Sibutramine hydrochloride | -9.9% | 49.7% | -34.8% | -14.9% | 125494-59-9 |
| Prestwick-P2 | 534 | F | 17 | Acenocoumarol | 1.7% | -4.7% | 0.6% | 6.3% | 152-72-7 |
| Prestwick-P2 | 535 | G | 17 | Nystatine | 2.1% | -0.4% | 0.9% | 0.4% | 1400-61-9 |
| Prestwick-P2 | 536 | H | 17 | Budesonide | 6.4% | 2.5% | -9.8% | -9.5% | 51333-22-3 |
| Prestwick-P2 | 537 | I | 17 | Tenoxicam | 2.6% | -5.4% | 7.1% | 7.6% | 59804-37-4 |
| Prestwick-P2 | 538 | J | 17 | Triflusal | -0.8% | -2.3% | -1.0% | -6.4% | 322-79-2 |
| Prestwick-P2 | 539 | K | 17 | Piracetam | 4.2% | -1.6% | 4.6% | 7.7% | 7491-74-9 |
| Prestwick-P2 | 540 | L | 17 | Phenindione | 3.6% | -3.6% | -0.3% | -0.7% | 83-12-5 |
| Prestwick-P2 | 541 | M | 17 | Molsidomine | 3.5% | -0.5% | 2.0% | -8.9% | 25717-80-0 |
| Prestwick-P2 | 542 | N | 17 | Chloroquine diphosphate | 1.9% | 3.4% | -53.2% | 4.6% | 50-63-5 |
| Prestwick-P2 | 543 | O | 17 | Dirithromycin | 7.2% | -0.9% | -24.2% | -1.5% | 62013-04-1 |
| Prestwick-P2 | 544 | P | 17 | Gliclazide | 0.3% | 2.8% | 7.5% | -1.4% | 21187-98-4 |
| Prestwick-P2 | 545 | A | 18 | Ceftazidime pentahydrate | -2.1% | -10.9% | -2.9% | 13.8% | 78439-06-2 |
| Prestwick-P2 | 546 | B | 18 | Iobenguane sulfate | -3.2% | -9.3% | 0.3% | 22.1% | 103346-16-3 |
| Prestwick-P2 | 547 | C | 18 | Propafenone hydrochloride | -1.6% | 0.3% | -4.3% | 1.5% | 34183-22-7 |
| Prestwick-P2 | 548 | D | 18 | Ethamivan | -0.7% | 0.8% | -6.8% | 0.2% | 304-84-7 |
| Prestwick-P2 | 549 | E | 18 | Bromperidol | -1.6% | 2.1% | -10.6% | 4.0% | 10457-90-6 |
| Prestwick-P2 | 550 | F | 18 | Cyclizine hydrochloride | 0.5% | 2.6% | -3.8% | 0.4% | 303-25-3 |
| Prestwick-P2 | 551 | G | 18 | Imipenem | 1.1% | -1.4% | 1.7% | -1.7% | 74431-23-5 |
| Prestwick-P2 | 552 | H | 18 | Sulfasalazine | 0.3% | -9.3% | 9.0% | -3.0% | 599-79-1 |
| Prestwick-P2 | 553 | I | 18 | Mesoridazine besylate | 0.1% | -5.8% | -10.2% | -4.7% | 32672-69-8 |
| Prestwick-P2 | 554 | J | 18 | Trolox | 1.0% | -1.6% | 7.5% | 4.4% | 53188-07-1 |
| Prestwick-P2 | 555 | K | 18 | Thiocolchicoside | 4.1% | -0.6% | 3.1% | 11.4% | 602-41-5 |
| Prestwick-P2 | 556 | L | 18 | Clorsulon | 3.8% | -6.6% | 22.2% | 16.6% | 60200-06-8 |
| Prestwick-P2 | 557 | M | 18 | Trimetazidine dihydrochloride | 1.0% | -5.7% | 19.4% | -0.9% | 13171-25-0 |
| Prestwick-P2 | 558 | N | 18 | Ropivacaine hydrochloride | 2.5% | -2.1% | 0.1% | -6.3% | 84057-95-4 |
| Prestwick-P2 | 559 | O | 18 | Tazarotene | 4.7% | 0.5% | 9.9% | -7.3% | 118292-40-3 |
| Prestwick-P2 | 560 | P | 18 | Prenylamine lactate | -15.9% | 63.6% | -73.9% | -18.6% | 69-43-2 |
| Prestwick-P2 | 561 | A | 19 | Ziprasidone hydrochloride | -0.8% | 7.1% | -39.0% | 16.1% | 138982-67-9 |
| Prestwick-P2 | 562 | B | 19 | Pomalidomide | -0.4% | -8.5% | -2.2% | 11.0% | 19171-19-8 |
| Prestwick-P2 | 563 | C | 19 | Tetracaïne hydrochloride | 0.6% | -2.3% | 2.2% | 7.6% | 136-47-0 |
| Prestwick-P2 | 564 | D | 19 | Mometasone furoate | -11.1% | 32.4% | -71.1% | -54.0% | 83919-23-7 |
| Prestwick-P2 | 565 | E | 19 | Reboxetine mesylate | 0.9% | 1.2% | -5.2% | 2.7% | 98769-81-4 |
| Prestwick-P2 | 566 | F | 19 | Camylofine chlorhydrate | 1.3% | 4.0% | -11.3% | 3.4% | 54-30-8 |
| Prestwick-P2 | 567 | G | 19 | Emedastine difumarate | 2.5% | -4.3% | 8.1% | -3.0% | 87233-61-2 |
| Prestwick-P2 | 568 | H | 19 | Etofenamate | -11.4% | 46.8% | -23.6% | 0.9% | 30544-47-9 |
| Prestwick-P2 | 569 | I | 19 | Tranilast | 1.0% | -3.2% | 8.4% | 6.2% | 53902-12-8 |
| Prestwick-P2 | 570 | J | 19 | Tizanidine hydrochloride | 2.5% | 0.0% | -8.8% | -3.3% | 51322-75-9 |
| Prestwick-P2 | 571 | K | 19 | Loracarbef | 4.8% | -1.4% | 11.7% | 4.7% | 121961-22-6 |
| Prestwick-P2 | 572 | L | 19 | Fenipentol | 1.5% | -1.4% | 13.4% | 6.2% | 583-03-9 |
| Prestwick-P2 | 573 | M | 19 | Mizolastine | 2.5% | 3.0% | 12.7% | -0.7% | 108612-45-9 |
| Prestwick-P2 | 574 | N | 19 | Amisulpride | 4.9% | 1.7% | 14.4% | -13.3% | 71675-85-9 |
| Prestwick-P2 | 575 | O | 19 | Alendronate sodium | 2.3% | 1.8% | 7.5% | -5.1% | 121268-17-5 |
| Prestwick-P2 | 576 | P | 19 | Dipivefrin hydrochloride | -0.2% | 10.1% | 16.4% | -0.8% | 64019-93-8 |
| Prestwick-P2 | 577 | A | 20 | Pyridostigmine iodide | -2.8% | -10.3% | 3.8% | 13.8% | 4685-03-4 |
| Prestwick-P2 | 578 | B | 20 | Pentobarbital | -3.1% | -6.8% | -6.3% | 10.6% | 76-74-4 |
| Prestwick-P2 | 579 | C | 20 | Troglitazone | -0.5% | -1.4% | -13.6% | -4.5% | 97322-87-7 |
| Prestwick-P2 | 580 | D | 20 | Dacarbazine | 0.3% | -2.4% | 5.3% | -1.5% | 4342-03-4 |
| Prestwick-P2 | 581 | E | 20 | Papaverine hydrochloride | -3.4% | -1.6% | -4.7% | 2.2% | 61-25-6 |
| Prestwick-P2 | 582 | F | 20 | Yohimbine hydrochloride | -1.6% | -7.6% | -0.7% | 0.2% | 65-19-0 |
| Prestwick-P2 | 583 | G | 20 | Zaleplon | -1.1% | -4.0% | -7.3% | -2.0% | 151319-34-5 |
| Prestwick-P2 | 584 | H | 20 | Diclofenac sodium | -1.0% | -6.4% | -2.3% | 1.2% | 15307-79-6 |
| Prestwick-P2 | 585 | I | 20 | Zafirlukast | -0.2% | 0.8% | -2.0% | 1.2% | 107753-78-6 |
| Prestwick-P2 | 586 | J | 20 | Butenafine hydrochloride | -33.0% | 90.0% | -87.1% | -60.1% | 101828-21-1 |
| Prestwick-P2 | 587 | K | 20 | Diosmin | 3.2% | -2.1% | 13.7% | 11.7% | 520-27-4 |
| Prestwick-P2 | 588 | L | 20 | Carbidopa | 3.7% | 0.5% | 11.8% | 4.9% | 28860-95-9 |
| Prestwick-P2 | 589 | M | 20 | Pyridoxine hydrochloride | 1.6% | -0.5% | 13.4% | 0.3% | 58-56-0 |
| Prestwick-P2 | 590 | N | 20 | Mercaptopurine | 0.0% | -1.1% | 14.5% | -3.0% | 50-44-2 |
| Prestwick-P2 | 591 | O | 20 | Thiorphan | 3.5% | -2.2% | 9.1% | -6.1% | 76721-89-6 |
| Prestwick-P2 | 592 | P | 20 | Tomoxetine hydrochloride | -0.3% | 6.8% | 9.1% | -3.4% | 82248-59-7 |
| Prestwick-P2 | 593 | A | 21 | Atropine sulfate monohydrate | -4.5% | -10.8% | 1.7% | 16.1% | 5908-99-6 |
| Prestwick-P2 | 594 | B | 21 | Eserine hemisulfate salt | 0.6% | -8.9% | 6.2% | 14.5% | 64-47-1 |
| Prestwick-P2 | 595 | C | 21 | Tenatoprazole | 0.0% | -2.8% | -2.9% | 3.2% | 113712-98-4 |
| Prestwick-P2 | 596 | D | 21 | Acetopromazine maleate salt | 0.3% | -2.0% | -14.0% | 2.7% | 3598-37-6 |
| Prestwick-P2 | 597 | E | 21 | Voriconazole | 1.6% | -0.9% | 1.9% | 8.5% | 137234-62-9 |
| Prestwick-P2 | 598 | F | 21 | Alfacalcidol | -10.6% | 32.7% | -50.3% | -24.9% | 41294-56-8 |
| Prestwick-P2 | 599 | G | 21 | Exemestane | 2.0% | -2.6% | -5.5% | -6.4% | 107868-30-4 |
| Prestwick-P2 | 600 | H | 21 | Fomepizole | -1.4% | -1.3% | -10.9% | -4.6% | 7554-65-6 |
| Prestwick-P2 | 601 | I | 21 | Carbadox | 1.8% | -0.2% | 2.1% | 5.5% | 6804-07-5 |
| Prestwick-P2 | 602 | J | 21 | Rimantadine hydrochloride | 1.0% | 3.7% | 12.7% | 3.3% | 13392-28-4 |

| | | | | | | | | | |
|---|---|---|---|---|---|---|---|---|---|
| Prestwick-P2 | 603 | K | 21 | (-) Emtricitabine | 3.4% | -2.7% | 5.7% | 7.2% | 143491-57-0 |
| Prestwick-P2 | 604 | L | 21 | Demecarium bromide | 3.5% | 2.8% | 1.7% | 3.6% | 56-94-0 |
| Prestwick-P2 | 605 | M | 21 | Cytarabine | 5.2% | 1.3% | 15.8% | -11.1% | 147-94-4 |
| Prestwick-P2 | 606 | N | 21 | Racecadotril | 5.4% | 0.3% | 5.4% | -5.4% | 81110-73-8 |
| Prestwick-P2 | 607 | O | 21 | Aceclidine hydrochloride | 1.2% | 0.3% | 6.7% | -1.6% | 6109-70-2 |
| Prestwick-P2 | 608 | P | 21 | Penciclovir | 0.0% | 5.4% | 6.2% | -7.0% | 39809-25-1 |
| Prestwick-P2 | co | A | 22 | DMSO | -2.3% | -8.6% | -4.5% | 19.0% | |
| Prestwick-P2 | co | B | 22 | DMSO | -2.9% | -7.7% | -2.4% | 11.5% | |
| Prestwick-P2 | co | C | 22 | Rott2 | -18.9% | 50.5% | -83.7% | -18.5% | |
| Prestwick-P2 | co | D | 22 | Rott2 | -20.1% | 50.1% | -82.3% | -18.5% | |
| Prestwick-P2 | co | E | 22 | Rott0.5 | -9.9% | 12.3% | -50.3% | -10.4% | |
| Prestwick-P2 | co | F | 22 | Rott0.5 | -12.7% | 8.5% | -53.6% | -19.5% | |
| Prestwick-P2 | co | G | 22 | Rott0.13 | -6.8% | 1.2% | -14.9% | -13.9% | |
| Prestwick-P2 | co | H | 22 | Rott0.13 | -7.3% | 0.2% | -9.2% | -10.3% | |
| Prestwick-P2 | co | I | 22 | DMSO | 1.5% | 3.4% | 0.9% | 0.5% | |
| Prestwick-P2 | co | J | 22 | DMSO | 0.0% | -4.9% | -7.5% | -5.1% | |
| Prestwick-P2 | co | K | 22 | DMSO | 1.8% | 3.1% | 8.7% | 4.3% | |
| Prestwick-P2 | co | L | 22 | DMSO | 0.7% | 1.8% | 10.9% | 11.9% | |
| Prestwick-P2 | co | M | 22 | DMSO | 0.1% | 1.0% | 9.1% | 5.9% | |
| Prestwick-P2 | co | N | 22 | DMSO | 1.3% | -3.3% | 4.6% | 2.2% | |
| Prestwick-P2 | co | O | 22 | DMSO | 1.5% | 2.0% | 7.9% | -7.0% | |
| Prestwick-P2 | co | P | 22 | DMSO | -2.0% | 3.3% | 9.2% | -2.9% | |
| Prestwick-P2 | co | A | 23 | DMSO | -3.5% | -6.8% | -11.5% | 18.5% | |
| Prestwick-P2 | co | B | 23 | DMSO | 0.0% | -7.5% | 6.9% | 10.0% | |
| Prestwick-P2 | co | C | 23 | DMSO | -1.8% | -2.4% | 0.3% | -2.7% | |
| Prestwick-P2 | co | D | 23 | DMSO | -1.8% | -4.2% | -3.6% | -0.8% | |
| Prestwick-P2 | co | E | 23 | DMSO | -2.4% | 2.8% | -7.3% | 0.3% | |
| Prestwick-P2 | co | F | 23 | DMSO | -0.9% | -6.3% | 1.1% | -1.9% | |
| Prestwick-P2 | co | G | 23 | DMSO | 1.1% | 1.5% | 10.6% | 0.2% | |
| Prestwick-P2 | co | H | 23 | DMSO | -0.6% | 0.9% | -6.0% | -3.7% | |
| Prestwick-P2 | co | I | 23 | DMSO | 0.7% | 1.9% | -1.2% | 6.8% | |
| Prestwick-P2 | co | J | 23 | DMSO | 1.5% | -0.7% | -4.5% | -6.2% | |
| Prestwick-P2 | co | K | 23 | DMSO | 3.6% | -0.3% | 6.6% | 5.0% | |
| Prestwick-P2 | co | L | 23 | DMSO | 0.4% | 0.4% | 0.6% | 18.0% | |
| Prestwick-P2 | co | M | 23 | DMSO | 0.8% | -0.9% | -7.1% | 2.5% | |
| Prestwick-P2 | co | N | 23 | DMSO | 3.1% | 1.3% | 2.2% | -6.6% | |
| Prestwick-P2 | co | O | 23 | DMSO | -0.4% | 4.8% | 8.2% | -15.1% | |
| Prestwick-P2 | co | P | 23 | DMSO | 1.3% | 1.7% | 19.3% | 2.8% | |
| Prestwick-P2 | co | A | 24 | noPFF | -5.9% | -5.8% | -99.7% | 6.1% | |
| Prestwick-P2 | co | B | 24 | Ceph4 | -23.9% | 76.6% | -99.8% | -53.6% | |
| Prestwick-P2 | co | C | 24 | noPFF | 2.1% | -2.9% | -99.9% | 2.0% | |
| Prestwick-P2 | co | D | 24 | noPFF | 0.7% | -7.1% | -99.2% | 10.2% | |
| Prestwick-P2 | co | E | 24 | noPFF | -1.1% | -8.2% | -99.5% | 15.6% | |
| Prestwick-P2 | co | F | 24 | noPFF | 1.1% | 1.1% | -99.3% | -1.4% | |
| Prestwick-P2 | co | G | 24 | noPFF | 0.5% | -2.3% | -100.0% | 8.0% | |
| Prestwick-P2 | co | H | 24 | noPFF | -0.8% | 0.5% | -99.6% | 0.7% | |
| Prestwick-P2 | co | I | 24 | noPFF | 1.5% | 3.0% | -99.7% | -0.8% | |
| Prestwick-P2 | co | J | 24 | noPFF | -0.7% | -1.0% | -100.0% | 11.3% | |
| Prestwick-P2 | co | K | 24 | noPFF | 1.7% | -3.0% | -99.9% | 12.3% | |
| Prestwick-P2 | co | L | 24 | noPFF | 0.4% | -1.4% | -98.4% | 10.5% | |
| Prestwick-P2 | co | M | 24 | noPFF | 3.1% | 4.3% | -99.8% | 12.8% | |
| Prestwick-P2 | co | N | 24 | noPFF | 1.0% | -1.9% | -99.4% | -15.1% | |
| Prestwick-P2 | co | O | 24 | noPFF | -1.8% | 3.5% | -99.9% | 7.5% | |
| Prestwick-P2 | co | P | 24 | noPFF | -2.2% | 1.3% | -100.0% | 7.2% | |
| Prestwick-P3 | co | A | 1 | DMSO | -6.1% | -7.4% | -0.3% | -19.5% | |
| Prestwick-P3 | co | B | 1 | DMSO | 1.4% | 1.0% | 41.1% | -17.6% | |
| Prestwick-P3 | co | C | 1 | DMSO | 0.9% | -12.5% | 25.6% | -21.1% | |
| Prestwick-P3 | co | D | 1 | DMSO | 3.0% | 1.2% | 24.7% | -13.5% | |
| Prestwick-P3 | co | E | 1 | DMSO | 0.6% | -8.0% | 12.0% | -10.5% | |
| Prestwick-P3 | co | F | 1 | DMSO | 2.8% | 2.7% | 15.4% | -14.4% | |
| Prestwick-P3 | co | G | 1 | DMSO | 0.6% | -3.4% | -5.1% | -5.8% | |
| Prestwick-P3 | co | H | 1 | DMSO | 1.1% | 6.9% | -15.8% | -17.3% | |
| Prestwick-P3 | co | I | 1 | DMSO | 1.1% | 0.2% | -17.1% | -11.2% | |
| Prestwick-P3 | co | J | 1 | DMSO | 2.0% | 4.4% | -8.4% | -9.9% | |
| Prestwick-P3 | co | K | 1 | DMSO | -1.1% | -0.1% | -20.4% | -1.4% | |
| Prestwick-P3 | co | L | 1 | DMSO | 1.0% | 4.0% | -0.8% | 1.0% | |
| Prestwick-P3 | co | M | 1 | DMSO | 1.1% | -3.9% | 3.5% | -3.1% | |
| Prestwick-P3 | co | N | 1 | DMSO | 3.1% | -1.0% | 12.3% | 2.3% | |
| Prestwick-P3 | co | O | 1 | DMSO | 2.5% | 5.2% | -1.9% | -5.1% | |
| Prestwick-P3 | co | P | 1 | DMSO | -0.1% | 16.3% | -11.6% | -9.1% | |
| Prestwick-P3 | co | A | 2 | DMSO | -2.6% | -5.2% | 27.0% | -10.0% | |
| Prestwick-P3 | co | B | 2 | DMSO | -1.1% | 4.2% | 23.3% | -4.7% | |

| | | | | | | | | | |
|---|---|---|---|---|---|---|---|---|---|
| Prestwick-P3 | co | C | 2 | DMSO | 0.4% | -2.5% | 11.5% | -10.5% | |
| Prestwick-P3 | co | D | 2 | DMSO | -0.8% | 3.2% | 3.6% | -4.9% | |
| Prestwick-P3 | co | E | 2 | DMSO | -1.1% | -3.7% | -2.1% | -11.1% | |
| Prestwick-P3 | co | F | 2 | DMSO | 2.9% | -0.7% | 4.9% | -14.7% | |
| Prestwick-P3 | co | G | 2 | DMSO | -0.8% | -6.0% | -13.9% | -16.2% | |
| Prestwick-P3 | co | H | 2 | DMSO | 3.6% | 2.7% | 4.4% | -8.6% | |
| Prestwick-P3 | co | I | 2 | Til2 | -4.2% | 8.7% | -82.4% | -10.4% | |
| Prestwick-P3 | co | J | 2 | Til2 | -1.5% | 18.4% | -80.2% | -5.3% | |
| Prestwick-P3 | co | K | 2 | Til0.5 | 0.7% | 1.8% | -45.9% | -7.6% | |
| Prestwick-P3 | co | L | 2 | Til0.5 | 0.6% | 7.9% | -42.9% | -5.9% | |
| Prestwick-P3 | co | M | 2 | Til0.13 | 0.8% | -0.2% | -9.8% | 0.5% | |
| Prestwick-P3 | co | N | 2 | Til0.13 | 1.0% | 8.8% | -8.6% | 0.0% | |
| Prestwick-P3 | co | O | 2 | DMSO | -0.1% | 4.8% | -0.1% | 1.2% | |
| Prestwick-P3 | co | P | 2 | DMSO | 0.1% | 18.7% | 10.6% | 4.4% | |
| Prestwick-P3 | 609 | A | 3 | Itraconazole | -2.9% | 7.5% | -29.7% | -9.8% | 84625-61-6 |
| Prestwick-P3 | 610 | B | 3 | Acarbose | 1.2% | 3.0% | 16.7% | -3.9% | 56180-94-0 |
| Prestwick-P3 | 611 | C | 3 | Escitalopram oxalate | -1.9% | -0.8% | 9.1% | -10.6% | 128196-01-0 |
| Prestwick-P3 | 612 | D | 3 | Ropinirole hydrochloride | 0.7% | 2.8% | 18.1% | -6.9% | 91374-20-8 |
| Prestwick-P3 | 613 | E | 3 | Cilostazol | 0.6% | 0.5% | -3.2% | -11.0% | 73963-72-1 |
| Prestwick-P3 | 614 | F | 3 | Galanthamine hydrobromide | 2.9% | 4.2% | 3.1% | -12.3% | 1953-04-4 |
| Prestwick-P3 | 615 | G | 3 | Temozolomide | 0.7% | 1.1% | 6.2% | -8.8% | 85622-93-1 |
| Prestwick-P3 | 616 | H | 3 | Xylazine | 0.4% | 5.9% | 7.5% | -3.5% | 7361-61-7 |
| Prestwick-P3 | 617 | I | 3 | Tirofiban hydrochloride | 3.7% | 1.5% | 33.3% | -18.4% | 150915-40-5 |
| Prestwick-P3 | 618 | J | 3 | Oxibendazol | -14.6% | 60.6% | -55.0% | -30.8% | 20559-55-1 |
| Prestwick-P3 | 619 | K | 3 | Tolvaptan | 6.6% | -6.5% | -12.3% | -3.0% | 150683-30-0 |
| Prestwick-P3 | 620 | L | 3 | Acipimox | 4.7% | 2.0% | 7.3% | 2.3% | 51037-30-0 |
| Prestwick-P3 | 621 | M | 3 | Folic acid | 0.2% | -3.1% | 2.1% | -11.2% | 59-30-3 |
| Prestwick-P3 | 622 | N | 3 | Benazepril hydrochloride | 2.4% | 5.5% | 16.9% | -8.6% | 86541-74-4 |
| Prestwick-P3 | 623 | O | 3 | Levetiracetam | 2.4% | -3.1% | 13.6% | -4.3% | 102767-28-2 |
| Prestwick-P3 | 624 | P | 3 | Dexfenfluramine hydrochloride | 1.4% | 12.1% | 46.4% | -10.3% | 3239-45-0 |
| Prestwick-P3 | 625 | A | 4 | Entacapone | 0.5% | -5.9% | -1.4% | -18.7% | 130929-57-6 |
| Prestwick-P3 | 626 | B | 4 | Nicotinamide | -0.3% | 3.7% | 11.0% | -8.7% | 98-92-0 |
| Prestwick-P3 | 627 | C | 4 | Lacidipine | -9.7% | 33.3% | -56.3% | -36.0% | 103890-78-4 |
| Prestwick-P3 | 628 | D | 4 | Argatroban | 1.8% | 4.0% | 7.3% | -2.8% | 74863-84-6 |
| Prestwick-P3 | 629 | E | 4 | Azelastine hydrochloride | -1.0% | -7.7% | -28.8% | -8.1% | 79307-93-0 |
| Prestwick-P3 | 630 | F | 4 | Etretinate | 3.4% | 8.0% | -1.9% | -14.3% | 54350-48-0 |
| Prestwick-P3 | 631 | G | 4 | Celiprolol hydrochloride | 0.6% | -2.2% | -5.3% | 0.3% | 57470-78-7 |
| Prestwick-P3 | 632 | H | 4 | Zopiclone | 1.2% | 3.2% | 3.3% | -6.9% | 43200-80-2 |
| Prestwick-P3 | 633 | I | 4 | Ipsapirone | 1.3% | -0.9% | -0.4% | -10.6% | 95847-70-4 |
| Prestwick-P3 | 634 | J | 4 | Hydroxychloroquine sulfate | 0.4% | 5.5% | -19.3% | -0.4% | 747-36-4 |
| Prestwick-P3 | 635 | K | 4 | Diflorasone Diacetate | 2.7% | -3.5% | -16.6% | -9.3% | 33564-31-7 |
| Prestwick-P3 | 636 | L | 4 | Acamprosate calcium | 2.9% | 3.0% | 4.5% | -3.5% | 77337-73-6 |
| Prestwick-P3 | 637 | M | 4 | Aniracetam | 9.0% | -16.2% | 0.1% | -10.7% | 72432-10-1 |
| Prestwick-P3 | 638 | N | 4 | Dimethisoquin hydrochloride | 1.8% | 27.8% | 47.2% | -14.1% | 2773-92-4 |
| Prestwick-P3 | 639 | O | 4 | Etoricoxib | 0.5% | -3.0% | 8.2% | -8.6% | 202409-33-4 |
| Prestwick-P3 | 640 | P | 4 | Sertindole | -15.4% | 52.2% | -88.1% | -40.0% | 106516-24-9 |
| Prestwick-P3 | 641 | A | 5 | Sulmazole | -1.8% | -5.6% | -4.7% | -4.0% | 73384-60-8 |
| Prestwick-P3 | 642 | B | 5 | Gefitinib | 3.1% | 3.7% | -34.9% | -3.9% | 184475-35-2 |
| Prestwick-P3 | 643 | C | 5 | Glimepiride | 0.0% | -3.2% | -13.5% | -9.1% | 93479-97-1 |
| Prestwick-P3 | 644 | D | 5 | Picrotoxinin | 4.0% | 0.4% | 11.3% | -3.9% | 17617-45-7 |
| Prestwick-P3 | 645 | E | 5 | Pranlukast | 0.0% | -0.7% | 8.2% | -13.3% | 103177-37-3 |
| Prestwick-P3 | 646 | F | 5 | D,L-Penicillamine | 3.8% | 5.4% | -1.9% | -8.1% | 52-66-4 |
| Prestwick-P3 | 647 | G | 5 | Dydrogesterone | 1.8% | -2.4% | -4.4% | -1.6% | 152-62-5 |
| Prestwick-P3 | 648 | H | 5 | Sumatriptan succinate | 1.4% | 7.7% | 5.1% | -1.9% | 103628-48-4 |
| Prestwick-P3 | 649 | I | 5 | Pranoprofen | 2.3% | -2.8% | 2.3% | -14.5% | 52549-17-4 |
| Prestwick-P3 | 650 | J | 5 | Secnidazole | 0.7% | 2.3% | 3.7% | -6.8% | 3366-95-8 |
| Prestwick-P3 | 651 | K | 5 | Tylosin | 0.8% | -1.6% | -3.6% | -4.2% | 1401-69-0 |
| Prestwick-P3 | 652 | L | 5 | Citalopram hydrobromide | 3.7% | 2.7% | 5.2% | -6.7% | 59729-32-7 |
| Prestwick-P3 | 653 | M | 5 | Trihexyphenidyl-D,L hydrochloride | -0.1% | 5.4% | -3.2% | -10.7% | 58947-95-8 |
| Prestwick-P3 | 654 | N | 5 | Succinylsulfathiazole | -0.5% | 10.4% | 2.1% | -3.5% | 116-43-8 |
| Prestwick-P3 | 655 | O | 5 | Sulfabenzamide | 3.2% | -3.4% | 11.4% | -10.2% | 127-71-9 |
| Prestwick-P3 | 656 | P | 5 | Benzocaine | 0.7% | 10.9% | 7.1% | -6.8% | 94-09-7 |
| Prestwick-P3 | 657 | A | 6 | Flunisolide | -0.2% | -8.0% | -3.2% | -15.9% | 3385-03-3 |
| Prestwick-P3 | 658 | B | 6 | N-Acetyl-DL-homocysteine Thiolactone | 3.4% | -2.0% | 12.8% | -4.2% | 1195-16-0 |
| Prestwick-P3 | 659 | C | 6 | Mepenzolate bromide | 3.7% | -8.3% | -0.1% | -13.9% | 76-90-4 |
| Prestwick-P3 | 660 | D | 6 | Benfotiamine | 3.4% | 3.2% | 2.9% | -6.1% | 22457-89-2 |
| Prestwick-P3 | 661 | E | 6 | Zileuton | 3.4% | -5.6% | -7.7% | -7.1% | 111406-87-2 |
| Prestwick-P3 | 662 | F | 6 | Loratadine | 2.7% | 6.1% | -18.8% | -15.0% | 79794-75-5 |
| Prestwick-P3 | 663 | G | 6 | Opipramol dihydrochloride | -0.3% | 5.5% | -10.3% | -6.9% | 909-39-7 |
| Prestwick-P3 | 664 | H | 6 | Nalidixic acid sodium salt | 1.2% | 3.6% | 2.1% | -0.1% | 3374-05-8 |
| Prestwick-P3 | 665 | I | 6 | Tadalafil | 2.3% | -1.4% | -7.2% | -7.0% | 171596-29-5 |
| Prestwick-P3 | 666 | J | 6 | Mirabegron | -0.9% | 4.5% | -0.1% | 1.0% | 223673-61-8 |

| | | | | | | | | | |
|---|---|---|---|---|---|---|---|---|---|
| Prestwick-P3 | 667 | K | 6 | Promazine hydrochloride | 6.2% | -2.2% | 39.9% | -12.3% | 53-60-1 |
| Prestwick-P3 | 668 | L | 6 | Sulfamerazine | 3.1% | 1.4% | 18.7% | -5.0% | 127-79-7 |
| Prestwick-P3 | 669 | M | 6 | Famprofazone | -22.4% | 58.0% | -75.3% | -58.7% | 22881-35-2 |
| Prestwick-P3 | 670 | N | 6 | Bromopride | 1.4% | 6.9% | 5.8% | -8.7% | 4093-35-0 |
| Prestwick-P3 | 671 | O | 6 | Dipyrone sodium salt | 3.5% | 1.8% | -4.9% | -15.2% | 5907-38-0 |
| Prestwick-P3 | 672 | P | 6 | Isosorbide dinitrate | 2.3% | 5.7% | 14.6% | -7.5% | 87-33-2 |
| Prestwick-P3 | 673 | A | 7 | Flurandrenolide | -2.2% | -3.1% | 8.2% | -9.3% | 1524-88-5 |
| Prestwick-P3 | 674 | B | 7 | Oxiconazole nitrate | -20.0% | 55.6% | -62.2% | -44.9% | 64211-46-7 |
| Prestwick-P3 | 675 | C | 7 | Halcinonide | -0.4% | 9.3% | -22.9% | -17.2% | 3093-35-4 |
| Prestwick-P3 | 676 | D | 7 | Lanatoside C | -5.4% | 5.3% | -76.2% | -24.2% | 17575-22-3 |
| Prestwick-P3 | 677 | E | 7 | Tetraethylenepentamine pentahydrochloride | 3.2% | -3.6% | -7.6% | -22.9% | 4961-41-5 |
| Prestwick-P3 | 678 | F | 7 | Nisoldipine | -0.2% | 13.6% | -28.1% | -34.6% | 63675-72-9 |
| Prestwick-P3 | 679 | G | 7 | Oxacillin sodium | 0.6% | -2.5% | -2.2% | 3.3% | 1173-88-2 |
| Prestwick-P3 | 680 | H | 7 | Beta-Escin | -27.4% | 90.9% | -80.0% | -33.6% | 11072-93-8 |
| Prestwick-P3 | 681 | I | 7 | Ibutilide fumarate | 2.0% | -1.1% | -6.9% | -13.3% | 122647-32-9 |
| Prestwick-P3 | 682 | J | 7 | Tigecycline | 0.6% | 6.5% | -11.7% | -9.6% | 220620-09-7 |
| Prestwick-P3 | 683 | K | 7 | Venlafaxine | 1.5% | -3.2% | -7.0% | 1.8% | 93413-69-5 |
| Prestwick-P3 | 684 | L | 7 | Ethotoin | 3.4% | 1.9% | 4.3% | -1.4% | 86-35-1 |
| Prestwick-P3 | 685 | M | 7 | Methyl benzethonium chloride | -23.1% | 70.5% | -89.5% | -28.2% | 25155-18-4 |
| Prestwick-P3 | 686 | N | 7 | Chlorcyclizine hydrochloride | 0.9% | 12.2% | -8.0% | -6.4% | 1620-21-9 |
| Prestwick-P3 | 687 | O | 7 | Sulfachloropyridazine | 1.7% | 0.7% | -37.8% | -15.4% | 80-32-0 |
| Prestwick-P3 | 688 | P | 7 | Pramoxine hydrochloride | -1.1% | 18.4% | 12.4% | -12.5% | 637-58-1 |
| Prestwick-P3 | 689 | A | 8 | Rebamipide | 1.0% | -11.3% | -6.4% | -11.6% | 90098-04-7 |
| Prestwick-P3 | 690 | B | 8 | Nilvadipine | 0.2% | 9.9% | -27.2% | -12.7% | 75530-68-6 |
| Prestwick-P3 | 691 | C | 8 | Benzamil hydrochloride | 5.1% | -7.2% | -16.1% | -10.2% | 2898-76-2 |
| Prestwick-P3 | 692 | D | 8 | Suxibuzone | 4.4% | -4.5% | 1.3% | -0.2% | 27470-51-5 |
| Prestwick-P3 | 693 | E | 8 | Acefylline | 0.8% | -8.2% | -1.7% | -5.0% | 652-37-9 |
| Prestwick-P3 | 694 | F | 8 | Acitretin | 4.3% | 6.8% | -15.5% | -15.9% | 55079-83-9 |
| Prestwick-P3 | 695 | G | 8 | Thiamine hydrochloride | 3.3% | -5.3% | -0.6% | -0.3% | 67-03-8 |
| Prestwick-P3 | 696 | H | 8 | Tazobactam | 1.0% | 3.1% | -1.1% | 4.8% | 89786-04-9 |
| Prestwick-P3 | 697 | I | 8 | Tramadol hydrochloride | 3.1% | -4.0% | -0.2% | -3.9% | 27203-92-5 |
| Prestwick-P3 | 698 | J | 8 | Estropipate piperazine salt | 3.0% | 6.3% | 7.6% | -4.1% | 7280-37-7 |
| Prestwick-P3 | 699 | K | 8 | Dasabuvir | 1.8% | 3.6% | 8.0% | 4.5% | 1132935-63-7 |
| Prestwick-P3 | 700 | L | 8 | Tetrahydrozoline hydrochloride | 1.6% | -0.6% | -4.2% | 5.3% | 522-48-5 |
| Prestwick-P3 | 701 | M | 8 | Diphenylpyraline hydrochloride | 0.1% | 3.7% | 4.2% | 0.9% | 132-18-3 |
| Prestwick-P3 | 702 | N | 8 | Benzethonium chloride | -14.5% | 43.1% | -59.7% | -26.2% | 121-54-0 |
| Prestwick-P3 | 703 | O | 8 | Finasteride | 4.4% | -2.8% | -5.0% | -7.1% | 98319-26-7 |
| Prestwick-P3 | 704 | P | 8 | Fluorometholone | -2.6% | 11.5% | -5.4% | -15.0% | 426-13-1 |
| Prestwick-P3 | 705 | A | 9 | Etanidazole | -0.4% | -2.2% | -0.8% | 2.4% | 22668-01-5 |
| Prestwick-P3 | 706 | B | 9 | Pinaverium bromide | -1.7% | 7.3% | 4.4% | 0.4% | 53251-94-8 |
| Prestwick-P3 | 707 | C | 9 | 6-Furfurylaminopurine | 1.1% | -0.7% | -2.5% | -12.1% | 525-79-1 |
| Prestwick-P3 | 708 | D | 9 | Avermectin B1 | -12.4% | 32.5% | -66.8% | -31.8% | 71751-41-2 |
| Prestwick-P3 | 709 | E | 9 | Zonisamide | 1.0% | -3.5% | -5.0% | -7.2% | 68291-97-4 |
| Prestwick-P3 | 710 | F | 9 | Irsogladine maleate | 3.0% | -0.9% | 6.1% | -9.0% | 84504-69-8 |
| Prestwick-P3 | 711 | G | 9 | Ibandronate sodium | 0.4% | -5.1% | -3.7% | 6.6% | 114084-78-5 |
| Prestwick-P3 | 712 | H | 9 | Warfarin | -1.0% | 2.9% | 2.8% | 3.3% | 81-81-2 |
| Prestwick-P3 | 713 | I | 9 | N-Butylscopolammonium bromide | -0.6% | 1.5% | -11.1% | -3.4% | 149-64-4 |
| Prestwick-P3 | 714 | J | 9 | Irinotecan hydrochloride trihydrate | 0.9% | 4.0% | -5.7% | -22.8% | 136572-09-3 |
| Prestwick-P3 | 715 | K | 9 | Hexestrol | -2.4% | 16.4% | -24.0% | -20.9% | 84-16-2 |
| Prestwick-P3 | 716 | L | 9 | Cefmetazole sodium salt | 2.6% | -0.2% | 7.3% | -1.0% | 56796-39-5 |
| Prestwick-P3 | 717 | M | 9 | Trioxsalen | -1.1% | 11.1% | 12.5% | -12.1% | 3902-71-4 |
| Prestwick-P3 | 718 | N | 9 | Doxofylline | -0.8% | 4.6% | 6.3% | -2.4% | 69975-86-6 |
| Prestwick-P3 | 719 | O | 9 | Cephalothin sodium salt | -1.0% | -2.5% | 2.3% | -5.1% | 58-71-9 |
| Prestwick-P3 | 720 | P | 9 | Cefuroxime sodium salt | -1.2% | 8.0% | 15.7% | -9.2% | 56238-63-2 |
| Prestwick-P3 | 721 | A | 10 | Althiazide | -3.4% | -4.5% | 9.6% | 3.3% | 5588-16-9 |
| Prestwick-P3 | 722 | B | 10 | Isopyrin hydrochloride | -1.4% | 0.9% | 9.7% | 3.5% | 18342-39-7 |
| Prestwick-P3 | 723 | C | 10 | Sulfaquinoxaline sodium salt | -1.3% | -6.0% | -2.5% | -7.3% | 967-80-6 |
| Prestwick-P3 | 724 | D | 10 | Streptozotocin | 1.9% | 1.1% | 5.7% | -13.0% | 18883-66-4 |
| Prestwick-P3 | 725 | E | 10 | Bimatoprost | 0.9% | -4.5% | -1.4% | -9.3% | 155206-00-1 |
| Prestwick-P3 | 726 | F | 10 | Sulfamethizole | 1.6% | 1.5% | 6.9% | -6.6% | 144-82-1 |
| Prestwick-P3 | 727 | G | 10 | Terazosin hydrochloride | 1.0% | -6.1% | 1.2% | -3.5% | 63590-64-7 |
| Prestwick-P3 | 728 | H | 10 | Phenazopyridine hydrochloride | 1.9% | 0.9% | -3.4% | -1.5% | 136-40-3 |
| Prestwick-P3 | 729 | I | 10 | Butamben | -0.6% | -5.8% | -11.9% | -7.0% | 94-25-7 |
| Prestwick-P3 | 730 | J | 10 | Sulfapyridine | 1.1% | 0.7% | 2.0% | -2.2% | 144-83-2 |
| Prestwick-P3 | 731 | K | 10 | Alclometasone dipropionate | 1.6% | 3.8% | -8.7% | -12.2% | 66734-13-2 |
| Prestwick-P3 | 732 | L | 10 | Leflunomide | -0.4% | 0.7% | -0.7% | -0.6% | 75706-12-6 |
| Prestwick-P3 | 733 | M | 10 | Clobetasol propionate | -0.9% | 2.1% | -2.8% | -10.0% | 25122-46-7 |
| Prestwick-P3 | 734 | N | 10 | Podophyllotoxin | 14.4% | 7.0% | 21.1% | -23.9% | 518-28-5 |
| Prestwick-P3 | 735 | O | 10 | (R) Naproxen sodium salt | 1.4% | -3.4% | 16.9% | -12.3% | 23979-41-1 |
| Prestwick-P3 | 736 | P | 10 | Propidium iodide | -3.6% | 0.9% | 13.0% | -6.6% | 25535-16-4 |
| Prestwick-P3 | 737 | A | 11 | Phenethicillin potassium salt | -0.1% | -6.4% | 3.4% | -4.7% | 132-93-4 |
| Prestwick-P3 | 738 | B | 11 | Sulfamethoxypyridazine | 1.5% | -0.2% | 10.9% | -8.3% | 80-35-3 |

| | | | | | | | | | |
|---|---|---|---|---|---|---|---|---|---|
| Prestwick-P3 | 739 | C | 11 | Metoprolol-(+)-tartrate salt | 0.1% | -12.6% | -3.5% | -3.7% | 56392-17-7 |
| Prestwick-P3 | 740 | D | 11 | Flumethasone | 0.8% | 1.0% | 4.1% | -7.4% | 2135-17-3 |
| Prestwick-P3 | 741 | E | 11 | Medrysone | -0.4% | -5.2% | -5.1% | -9.4% | 2668-66-8 |
| Prestwick-P3 | 742 | F | 11 | Flunixin meglumine | 2.8% | 3.4% | -0.1% | -10.0% | 42461-84-7 |
| Prestwick-P3 | 743 | G | 11 | Demeclocycline hydrochloride | -0.8% | 2.7% | 15.0% | 5.3% | 64-73-3 |
| Prestwick-P3 | 744 | H | 11 | Fenoprofen calcium salt dihydrate | 1.4% | 3.1% | 4.4% | 0.2% | 53746-45-5 |
| Prestwick-P3 | 745 | I | 11 | Meclofenoxate hydrochloride | 0.6% | -3.4% | -9.0% | -6.6% | 3685-84-5 |
| Prestwick-P3 | 746 | J | 11 | Furaltadone hydrochloride | 0.3% | -0.6% | 9.3% | -1.5% | 3759-92-0 |
| Prestwick-P3 | 747 | K | 11 | (D-) Norgestrel | 3.0% | -3.4% | 6.7% | -1.3% | 797-63-7 |
| Prestwick-P3 | 748 | L | 11 | Fluocinonide | 2.6% | -0.7% | -10.6% | -1.8% | 356-12-7 |
| Prestwick-P3 | 749 | M | 11 | Clofibric acid | 2.0% | -5.4% | 4.7% | 0.0% | 882-09-7 |
| Prestwick-P3 | 750 | N | 11 | Bendroflumethiazide | 0.2% | 4.9% | 7.4% | 2.5% | 73-48-3 |
| Prestwick-P3 | 751 | O | 11 | Cloperastine hydrochloride | -1.8% | 12.3% | -3.0% | -15.5% | 14984-68-0 |
| Prestwick-P3 | 752 | P | 11 | Eprosartan mesylate | 1.6% | 5.8% | 7.3% | -18.6% | 133040-01-4 |
| Prestwick-P3 | 753 | A | 12 | Deferoxamine mesylate | -2.4% | -5.4% | -19.2% | 1.3% | 138-14-7 |
| Prestwick-P3 | 754 | B | 12 | Mephentermine hemisulfate | -3.1% | 6.7% | -2.2% | -5.7% | 1212-72-2 |
| Prestwick-P3 | 755 | C | 12 | Flecainide acetate | 0.3% | -7.0% | -11.8% | -1.3% | 54143-56-5 |
| Prestwick-P3 | 756 | D | 12 | Cefazolin sodium salt | 2.2% | -0.4% | 11.2% | -8.2% | 27164-46-1 |
| Prestwick-P3 | 757 | E | 12 | Spiramycin | -0.3% | 1.6% | -23.1% | -4.8% | 8025-81-8 |
| Prestwick-P3 | 758 | F | 12 | Glycopyrrolate bromide | -0.9% | -1.5% | -5.5% | -3.9% | 596-51-0 |
| Prestwick-P3 | 759 | G | 12 | Piperacillin sodium salt | -0.8% | -8.1% | 12.1% | 5.8% | 59703-84-3 |
| Prestwick-P3 | 760 | H | 12 | Diethylstilbestrol | -18.9% | 22.6% | -52.3% | -57.1% | 56-53-1 |
| Prestwick-P3 | 761 | I | 12 | Ethoxyquin | 0.5% | 3.3% | -21.0% | -8.8% | 91-53-2 |
| Prestwick-P3 | 762 | J | 12 | Tinidazole | -0.5% | 7.5% | 3.5% | -3.6% | 19387-91-8 |
| Prestwick-P3 | 763 | K | 12 | Sulfamethazine sodium salt | -0.3% | -3.8% | 6.0% | 9.6% | 1981-58-4 |
| Prestwick-P3 | 764 | L | 12 | Guaifenesin | 1.0% | -3.7% | -0.7% | -3.1% | 93-14-1 |
| Prestwick-P3 | 765 | M | 12 | Dicumarol | -0.6% | 1.2% | 1.8% | 3.1% | 66-76-2 |
| Prestwick-P3 | 766 | N | 12 | Methimazole | 0.1% | 3.4% | 12.3% | 9.1% | 60-56-0 |
| Prestwick-P3 | 767 | O | 12 | Isocarboxazid | 0.0% | 1.3% | 4.1% | -7.3% | 59-63-2 |
| Prestwick-P3 | 768 | P | 12 | Lithocholic acid | -1.3% | 7.4% | 6.5% | -13.3% | 434-13-9 |
| Prestwick-P3 | 769 | A | 13 | Liranaftate | -13.5% | 6.2% | -1.6% | -9.6% | 88678-31-3 |
| Prestwick-P3 | 770 | B | 13 | Sulfadimethoxine | 0.9% | -2.4% | 22.0% | 2.6% | 122-11-2 |
| Prestwick-P3 | 771 | C | 13 | Trimetozine | -0.2% | -7.8% | -2.0% | 5.0% | 635-41-6 |
| Prestwick-P3 | 772 | D | 13 | Folinic acid calcium salt | 0.6% | 0.1% | 13.9% | 6.7% | 6035-45-6 |
| Prestwick-P3 | 773 | E | 13 | Aprepitant | -23.7% | 76.0% | -64.3% | -22.9% | 170729-80-3 |
| Prestwick-P3 | 774 | F | 13 | Monensin sodium salt | -8.7% | 13.8% | -94.4% | -27.7% | 22373-78-0 |
| Prestwick-P3 | 775 | G | 13 | Chlorotrianisene | -2.9% | 1.5% | -5.9% | -0.6% | 569-57-3 |
| Prestwick-P3 | 776 | H | 13 | Ribostamycin sulfate salt | 2.3% | -1.3% | 8.2% | -9.1% | 53797-35-6 |
| Prestwick-P3 | 777 | I | 13 | Guanadrel sulfate | 1.1% | -0.1% | -15.2% | -4.3% | 22195-34-2 |
| Prestwick-P3 | 778 | J | 13 | Vidarabine | 4.2% | 4.3% | -0.2% | -1.5% | 5536-17-4 |
| Prestwick-P3 | 779 | K | 13 | Alexidine dihydrochloride | -26.4% | 69.0% | -90.7% | 15.7% | 22573-93-9 |
| Prestwick-P3 | 780 | L | 13 | Alogliptin benzoate | 1.3% | -1.7% | -6.2% | -4.2% | 850649-62-6 |
| Prestwick-P3 | 781 | M | 13 | Merbromin disodium salt | 1.8% | -2.6% | 24.7% | -15.2% | 129-16-8 |
| Prestwick-P3 | 782 | N | 13 | Hexylcaine hydrochloride | 1.2% | 3.8% | -0.9% | -9.5% | 532-76-3 |
| Prestwick-P3 | 783 | O | 13 | Methotrimeprazine maleate salt | 11.5% | 1.9% | 57.0% | -10.8% | 7104-38-3 |
| Prestwick-P3 | 784 | P | 13 | Dienestrol | -5.4% | 25.8% | -35.3% | -25.0% | 84-17-3 |
| Prestwick-P3 | 785 | A | 13 | Sulfanilamide | -0.3% | -9.0% | -21.7% | -5.5% | 63-74-1 |
| Prestwick-P3 | 786 | B | 14 | Balsalazide disodium salt | -0.3% | -4.4% | 0.3% | -0.2% | 80573-04-2 |
| Prestwick-P3 | 787 | C | 14 | Levonordefrin | 0.1% | -3.4% | 0.8% | -5.1% | 829-74-3 |
| Prestwick-P3 | 788 | D | 14 | Amprenavir | 1.2% | 0.6% | 18.8% | -5.9% | 161814-49-9 |
| Prestwick-P3 | 789 | E | 14 | Isoetharine mesylate salt | 1.2% | -2.5% | -8.4% | 0.5% | 7279-75-6 |
| Prestwick-P3 | 790 | F | 14 | Dronedarone hydrochloride | -34.2% | 109.2% | -95.3% | -37.6% | 141626-36-0 |
| Prestwick-P3 | 791 | G | 14 | Methacholine chloride | -0.4% | -1.0% | 4.1% | 2.0% | 62-51-1 |
| Prestwick-P3 | 792 | H | 14 | Pipenzolate bromide | 0.3% | -0.5% | 10.7% | 3.1% | 125-51-9 |
| Prestwick-P3 | 793 | I | 14 | Sulfameter | 1.8% | -2.0% | 2.3% | -1.0% | 651-06-9 |
| Prestwick-P3 | 794 | J | 14 | Isopropamide iodide | 0.4% | -2.0% | -2.4% | -2.8% | 71-81-8 |
| Prestwick-P3 | 795 | K | 14 | Zomepirac sodium salt | 1.9% | -2.4% | 19.5% | 6.1% | 64092-48-4 |
| Prestwick-P3 | 796 | L | 14 | Cinoxacin | 1.6% | -4.3% | 7.7% | 5.4% | 28657-80-9 |
| Prestwick-P3 | 797 | M | 14 | Drofenine hydrochloride | -11.9% | 39.3% | -32.7% | -20.7% | 548-66-3 |
| Prestwick-P3 | 798 | N | 14 | Cycloheximide | 2.2% | -8.8% | -83.0% | -4.7% | 66-81-9 |
| Prestwick-P3 | 799 | O | 14 | Pridinol methanesulfonate salt | -0.1% | -1.7% | 0.5% | -3.1% | 6856-31-1 |
| Prestwick-P3 | 800 | P | 14 | Amrinone | -0.9% | 6.7% | 9.4% | -11.3% | 60719-84-8 |
| Prestwick-P3 | 801 | A | 15 | Carbinoxamine maleate salt | -1.8% | -1.0% | 12.3% | 7.1% | 3505-38-2 |
| Prestwick-P3 | 802 | B | 15 | Methazolamide | 0.5% | -5.2% | 3.9% | 8.5% | 554-57-4 |
| Prestwick-P3 | 803 | C | 15 | Auranofin | -16.6% | 42.3% | -99.5% | -33.6% | 34031-32-8 |
| Prestwick-P3 | 804 | D | 15 | Cromolyn disodium salt | 1.8% | 1.1% | 2.7% | -1.4% | 15826-37-6 |
| Prestwick-P3 | 805 | E | 15 | Azlocillin sodium salt | 1.2% | -0.8% | 4.4% | 6.3% | 37091-65-9 |
| Prestwick-P3 | 806 | F | 15 | Clidinium bromide | -0.1% | 2.2% | -10.3% | -6.4% | 3485-62-9 |
| Prestwick-P3 | 807 | G | 15 | Butacaine | -2.8% | 8.0% | -9.2% | 13.6% | 149-16-6 |
| Prestwick-P3 | 808 | H | 15 | Cefoxitin sodium salt | 4.2% | 3.5% | 7.9% | -3.2% | 33564-30-6 |
| Prestwick-P3 | 809 | I | 15 | Olanzapine | 0.0% | 4.7% | -8.0% | -2.9% | 132539-06-1 |
| Prestwick-P3 | 810 | J | 15 | Trimeprazine tartrate | 10.0% | 7.0% | 67.6% | -4.6% | 4330-99-8 |

| | | | | | | | | | |
|---|---|---|---|---|---|---|---|---|---|
| Prestwick-P3 | 811 | K | 15 | Paroxetine hydrochloride | -2.2% | 9.9% | -23.4% | -0.1% | 110429-35-1 |
| Prestwick-P3 | 812 | L | 15 | Nylidrin | -1.0% | 8.2% | 2.1% | 18.6% | 447-41-6 |
| Prestwick-P3 | 813 | M | 15 | Gabapentin | 1.8% | 0.2% | 3.5% | -4.4% | 60142-96-3 |
| Prestwick-P3 | 814 | N | 15 | Raloxifene hydrochloride | 0.5% | 16.6% | -83.4% | -8.3% | 82640-04-8 |
| Prestwick-P3 | 815 | O | 15 | Iopamidol | 0.9% | 2.8% | 4.1% | -4.8% | 60166-93-0 |
| Prestwick-P3 | 816 | P | 15 | Iopromide | 0.9% | 5.1% | 9.4% | -12.8% | 73334-07-3 |
| Prestwick-P3 | 817 | A | 16 | Pyrithyldione | 0.0% | -4.9% | 2.2% | -7.6% | 77-04-3 |
| Prestwick-P3 | 818 | B | 16 | Spectinomycin dihydrochloride | -1.3% | -3.2% | -5.0% | 5.0% | 21736-83-4 |
| Prestwick-P3 | 819 | C | 16 | Bucladesine sodium salt | -1.0% | -3.6% | 2.3% | -0.3% | 16980-89-5 |
| Prestwick-P3 | 820 | D | 16 | Cefsulodin sodium salt | -0.3% | 4.5% | -6.7% | -5.2% | 52152-93-9 |
| Prestwick-P3 | 821 | E | 16 | Sulfamonomethoxine | 1.3% | -4.3% | 1.5% | 2.4% | 1220-83-3 |
| Prestwick-P3 | 822 | F | 16 | Benzthiazide | -0.4% | 4.7% | -5.5% | 9.1% | 91-33-8 |
| Prestwick-P3 | 823 | G | 16 | Eszopiclone | 1.2% | 2.7% | 14.1% | 23.2% | 138729-47-2 |
| Prestwick-P3 | 824 | H | 16 | Novobiocin sodium salt | 0.4% | 2.2% | 8.3% | -2.5% | 1476-53-5 |
| Prestwick-P3 | 825 | I | 16 | Nafcillin sodium salt monohydrate | 1.3% | 2.2% | -1.4% | -5.6% | 7177-50-6 |
| Prestwick-P3 | 826 | J | 16 | Procyclidine hydrochloride | -0.8% | 1.9% | -10.2% | 8.4% | 1508-76-5 |
| Prestwick-P3 | 827 | K | 16 | Liothyronine | -1.5% | 4.3% | 4.2% | 4.6% | 6893-02-3 |
| Prestwick-P3 | 828 | L | 16 | Roxithromycin | -0.2% | 0.0% | -10.0% | 6.2% | 80214-83-1 |
| Prestwick-P3 | 829 | M | 16 | Ciclesonide | -7.6% | 12.6% | -51.4% | -13.0% | 126544-47-6 |
| Prestwick-P3 | 830 | N | 16 | Aclidinium bromide | 1.5% | 5.1% | 15.5% | -11.1% | 320345-99-1 |
| Prestwick-P3 | 831 | O | 16 | Theophylline monohydrate | 0.9% | 1.8% | -0.6% | -2.0% | 5967-84-0 |
| Prestwick-P3 | 832 | P | 16 | Theobromine | 0.3% | 1.8% | 7.4% | -16.0% | 83-67-0 |
| Prestwick-P3 | 833 | A | 17 | Piromidic acid | -6.1% | 31.4% | -26.5% | -52.8% | 19562-30-2 |
| Prestwick-P3 | 834 | B | 17 | Trimipramine maleate salt | 12.0% | -4.5% | 72.2% | -2.0% | 521-78-8 |
| Prestwick-P3 | 835 | C | 17 | Fosfosal | 4.7% | -7.6% | 0.3% | 7.1% | 6064-83-1 |
| Prestwick-P3 | 836 | D | 17 | Suprofen | 2.3% | -1.9% | -8.2% | -3.5% | 40828-46-4 |
| Prestwick-P3 | 837 | E | 17 | Trichlormethiazide | 4.3% | 1.7% | 0.6% | 1.5% | 133-67-5 |
| Prestwick-P3 | 838 | F | 17 | Oxolamine citrate salt | -1.0% | 0.8% | -4.7% | 5.9% | 1949-20-8 |
| Prestwick-P3 | 839 | G | 17 | Zolmitriptan | 2.3% | -4.1% | 1.9% | 4.2% | 139264-17-8 |
| Prestwick-P3 | 840 | H | 17 | Indoprofen | 3.0% | -1.6% | 3.9% | 9.2% | 31842-01-0 |
| Prestwick-P3 | 841 | I | 17 | Amiprilose hydrochloride | 0.3% | 1.8% | -8.2% | 5.2% | 60414-06-4 |
| Prestwick-P3 | 842 | J | 17 | Ethynylestradiol 3-methyl ether | -19.6% | 57.6% | -54.7% | -29.1% | 72-33-3 |
| Prestwick-P3 | 843 | K | 17 | Beclomethasone dipropionate | 3.5% | -1.4% | -17.2% | 4.1% | 5534-09-8 |
| Prestwick-P3 | 844 | L | 17 | Tolmetin sodium salt dihydrate | -1.0% | 2.4% | -6.9% | 4.0% | 64490-92-2 |
| Prestwick-P3 | 845 | M | 17 | Simvastatin | -21.9% | 65.9% | -75.9% | -5.2% | 79902-63-9 |
| Prestwick-P3 | 846 | N | 17 | Azacytidine-5 | 4.7% | -3.9% | -5.0% | -7.8% | 320-67-2 |
| Prestwick-P3 | 847 | O | 17 | Reserpine | 6.5% | 5.0% | -49.9% | -12.1% | 50-55-5 |
| Prestwick-P3 | 848 | P | 17 | Bicalutamide | 1.0% | 3.4% | 10.5% | -14.5% | 90357-06-5 |
| Prestwick-P3 | 849 | A | 18 | Chloropyramine hydrochloride | 1.6% | -6.1% | 36.3% | 12.9% | 6170-42-9 |
| Prestwick-P3 | 850 | B | 18 | Furazolidone | 0.6% | -7.7% | -0.9% | 11.4% | 67-45-8 |
| Prestwick-P3 | 851 | C | 18 | Deflazacort | 1.8% | -8.1% | 0.2% | 1.2% | 14484-47-0 |
| Prestwick-P3 | 852 | D | 18 | Nadolol | 1.2% | -2.9% | -3.7% | 4.2% | 42200-33-9 |
| Prestwick-P3 | 853 | E | 18 | Propantheline bromide | 0.7% | -2.2% | 3.0% | -2.9% | 50-34-0 |
| Prestwick-P3 | 854 | F | 18 | Viloxazine hydrochloride | -2.6% | 2.0% | -9.6% | 1.5% | 35604-67-2 |
| Prestwick-P3 | 855 | G | 18 | Carbenoxolone disodium salt | 0.6% | -2.7% | 4.1% | 6.6% | 7421-40-1 |
| Prestwick-P3 | 856 | H | 18 | Iocetamic acid | -0.7% | 0.7% | -7.8% | 0.2% | 16034-77-8 |
| Prestwick-P3 | 857 | I | 18 | (-) Levobunolol hydrochloride | 0.8% | -1.2% | -8.9% | -7.8% | 27912-14-7 |
| Prestwick-P3 | 858 | J | 18 | Iodixanol | 1.9% | 3.3% | 2.7% | -15.0% | 92339-11-2 |
| Prestwick-P3 | 859 | K | 18 | (+) Levobunolol hydrochloride | 0.3% | 3.5% | -18.2% | 5.0% | 47141-41-3 |
| Prestwick-P3 | 860 | L | 18 | Doxazosin mesylate | 0.7% | 0.4% | -26.5% | 5.1% | 77883-43-3 |
| Prestwick-P3 | 861 | M | 18 | Paromomycin sulfate | 0.8% | -2.9% | -5.0% | 3.8% | 1263-89-4 |
| Prestwick-P3 | 862 | N | 18 | Acetaminophen | 2.0% | -1.5% | -3.7% | -0.6% | 103-90-2 |
| Prestwick-P3 | 863 | O | 18 | Scopolamine hydrochloride | -0.3% | 0.9% | -6.0% | -6.3% | 55-16-3 |
| Prestwick-P3 | 864 | P | 18 | Ioversol | 1.7% | 1.6% | 13.0% | -12.3% | 87771-40-2 |
| Prestwick-P3 | 865 | A | 19 | Dichlorphenamide | 1.0% | -5.9% | -7.0% | 23.0% | 120-97-8 |
| Prestwick-P3 | 866 | B | 19 | Sulconazole nitrate | -22.2% | 61.8% | -63.6% | -20.8% | 61318-91-0 |
| Prestwick-P3 | 867 | C | 19 | Moxalactam disodium salt | 2.0% | -6.6% | -10.9% | 3.9% | 64953-12-4 |
| Prestwick-P3 | 868 | D | 19 | Aminophylline | 0.0% | -3.0% | -1.5% | 2.4% | 317-34-0 |
| Prestwick-P3 | 869 | E | 19 | Dimethadione | 1.1% | -4.9% | 2.5% | -1.4% | 695-53-4 |
| Prestwick-P3 | 870 | F | 19 | Ethaverine hydrochloride | -6.4% | 4.3% | -25.0% | -11.7% | 985-13-7 |
| Prestwick-P3 | 871 | G | 19 | Ganciclovir | 2.4% | -1.4% | -3.1% | -2.9% | 82410-32-0 |
| Prestwick-P3 | 872 | H | 19 | Ethopropazine hydrochloride | 5.9% | 6.7% | 66.4% | -6.7% | 1094-08-2 |
| Prestwick-P3 | 873 | I | 19 | Clinafloxacin | -0.8% | 1.2% | -15.8% | -0.1% | 105956-97-6 |
| Prestwick-P3 | 874 | J | 19 | Equilin | 2.0% | 6.1% | -13.2% | -8.2% | 474-86-2 |
| Prestwick-P3 | 875 | K | 19 | Fluvastatin sodium salt | -0.1% | 9.6% | -5.0% | 8.1% | 93957-55-2 |
| Prestwick-P3 | 876 | L | 19 | Octoclothepin maleate salt | -15.3% | 50.5% | -55.5% | -0.2% | 4789-68-8 |
| Prestwick-P3 | 877 | M | 19 | Phthalylsulfathiazole | -0.9% | 0.9% | -15.5% | 7.3% | 85-73-4 |
| Prestwick-P3 | 878 | N | 19 | Luteolin | -3.8% | 8.1% | -9.3% | 7.7% | 491-70-3 |
| Prestwick-P3 | 879 | O | 19 | Rabeprazole sodium salt | -0.6% | 3.0% | 0.4% | 2.2% | 117976-89-3 |
| Prestwick-P3 | 880 | P | 19 | Carbachol chloride | -4.2% | 6.2% | -3.0% | 5.5% | 51-83-2 |
| Prestwick-P3 | 881 | A | 20 | Niacin | -1.0% | -3.9% | -8.4% | 18.7% | 59-67-6 |
| Prestwick-P3 | 882 | B | 20 | Bemegride | -2.1% | -6.2% | -13.5% | 9.6% | 64-65-3 |

| | | | | | | | | | |
|---|---|---|---|---|---|---|---|---|---|
| Prestwick-P3 | 883 | C | 20 | Cortisol acetate | 0.9% | -2.8% | -11.5% | -0.8% | 50-03-3 |
| Prestwick-P3 | 884 | D | 20 | Flubendazol | -6.6% | 30.4% | -25.0% | -21.6% | 31430-15-6 |
| Prestwick-P3 | 885 | E | 20 | Hymecromone | 1.2% | -1.3% | -1.7% | 3.2% | 90-33-5 |
| Prestwick-P3 | 886 | F | 20 | Abacavir sulfate | 0.2% | 1.0% | -5.7% | 1.1% | 188062-50-2 |
| Prestwick-P3 | 887 | G | 20 | (+) Isoproterenol-(+)-bitartrate salt | 1.8% | -0.9% | -6.1% | 8.4% | 14638-70-1 |
| Prestwick-P3 | 888 | H | 20 | Monobenzone | 4.2% | -3.2% | 3.2% | -12.2% | 103-16-2 |
| Prestwick-P3 | 889 | I | 20 | Nizatidine | 0.4% | 1.7% | -16.7% | -14.9% | 76963-41-2 |
| Prestwick-P3 | 890 | J | 20 | Thioperamide maleate | -2.5% | 2.8% | -11.3% | 0.2% | 106243-16-7 |
| Prestwick-P3 | 891 | K | 20 | Propofol | -1.8% | 7.0% | -25.5% | 8.3% | 2078-54-8 |
| Prestwick-P3 | 892 | L | 20 | (S,-) Eticlopride hydrochloride | -4.8% | 21.3% | -24.3% | -9.6% | 97612-24-3 |
| Prestwick-P3 | 893 | M | 20 | Pentetic acid | 1.6% | -0.6% | -8.4% | 1.9% | 67-43-6 |
| Prestwick-P3 | 894 | N | 20 | Bretylium tosylate | 0.1% | -0.5% | -2.3% | 3.3% | 61-75-6 |
| Prestwick-P3 | 895 | O | 20 | Crotamiton | 1.2% | -2.3% | -4.3% | 2.1% | 483-63-6 |
| Prestwick-P3 | 896 | P | 20 | Toremifene | -20.7% | 51.1% | -67.0% | -33.0% | 89778-26-7 |
| Prestwick-P3 | 897 | A | 21 | Digoxigenin | -11.0% | -4.3% | -72.9% | -4.2% | 1672-46-4 |
| Prestwick-P3 | 898 | B | 21 | Meglumine | 1.2% | -7.8% | -5.2% | 11.2% | 6284-40-8 |
| Prestwick-P3 | 899 | C | 21 | Felbinac | -0.1% | -2.4% | -10.2% | 8.6% | 5728-52-9 |
| Prestwick-P3 | 900 | D | 21 | Butylparaben | -0.3% | -3.5% | -13.8% | 10.3% | 94-26-8 |
| Prestwick-P3 | 901 | E | 21 | Diloxanide furoate | 3.6% | 0.4% | -5.2% | -1.1% | 3736-81-0 |
| Prestwick-P3 | 902 | F | 21 | Metyrapone | 1.9% | -2.4% | 0.3% | -0.3% | 54-36-4 |
| Prestwick-P3 | 903 | G | 21 | 2-Aminobenzenesulfonamide | 0.4% | -3.8% | -13.1% | 8.0% | 3306-62-5 |
| Prestwick-P3 | 904 | H | 21 | Estrone | 0.9% | -2.9% | -3.5% | -5.3% | 53-16-7 |
| Prestwick-P3 | 905 | I | 21 | Xamoterol hemifumarate | -0.4% | 2.1% | -6.0% | -2.6% | 73210-73-8 |
| Prestwick-P3 | 906 | J | 21 | Posaconazole hydrate | -2.8% | 8.6% | -41.9% | -6.1% | 171228-49-2 |
| Prestwick-P3 | 907 | K | 21 | Primidone | 2.2% | 0.3% | -6.7% | -5.7% | 125-33-7 |
| Prestwick-P3 | 908 | L | 21 | Flucytosine | 0.6% | 0.3% | -5.7% | 8.9% | 2022-85-7 |
| Prestwick-P3 | 909 | M | 21 | Pralidoxime chloride | 0.2% | -1.0% | 11.1% | 4.9% | 51-15-0 |
| Prestwick-P3 | 910 | N | 21 | Phenoxybenzamine hydrochloride | -19.0% | 49.8% | -77.5% | -22.2% | 63-92-3 |
| Prestwick-P3 | 911 | O | 21 | (R,+) Atenolol | 0.5% | 4.1% | -2.3% | 11.6% | 56715-13-0 |
| Prestwick-P3 | 912 | P | 21 | Tyloxapol | -2.1% | 13.7% | -23.7% | 4.1% | 25301-02-4 |
| Prestwick-P3 | co | A | 22 | DMSO | -3.7% | -8.9% | -0.2% | 22.1% | |
| Prestwick-P3 | co | B | 22 | DMSO | -1.3% | -5.1% | -7.6% | 13.1% | |
| Prestwick-P3 | co | C | 22 | Rott2 | -19.0% | 35.5% | -85.0% | -16.2% | |
| Prestwick-P3 | co | D | 22 | Rott2 | -18.0% | 34.6% | -85.6% | -19.8% | |
| Prestwick-P3 | co | E | 22 | Rott0.5 | -8.6% | 13.3% | -56.2% | -16.4% | |
| Prestwick-P3 | co | F | 22 | Rott0.5 | -6.0% | 14.0% | -53.9% | -19.0% | |
| Prestwick-P3 | co | G | 22 | Rott0.13 | -3.6% | 1.9% | -13.9% | 1.6% | |
| Prestwick-P3 | co | H | 22 | Rott0.13 | -6.1% | -1.8% | -11.8% | -7.4% | |
| Prestwick-P3 | co | I | 22 | DMSO | -1.1% | -0.4% | -9.3% | -1.1% | |
| Prestwick-P3 | co | J | 22 | DMSO | -1.0% | 3.5% | -2.1% | -1.6% | |
| Prestwick-P3 | co | K | 22 | DMSO | 1.7% | 2.0% | 0.1% | 12.1% | |
| Prestwick-P3 | co | L | 22 | DMSO | -0.3% | 0.1% | -2.4% | 12.9% | |
| Prestwick-P3 | co | M | 22 | DMSO | -0.1% | 3.8% | -1.6% | 12.0% | |
| Prestwick-P3 | co | N | 22 | DMSO | -0.6% | 0.3% | -1.8% | 10.4% | |
| Prestwick-P3 | co | O | 22 | DMSO | 0.1% | 2.9% | -0.8% | 6.5% | |
| Prestwick-P3 | co | P | 22 | DMSO | 0.1% | -2.2% | 7.9% | 8.7% | |
| Prestwick-P3 | co | A | 23 | DMSO | -2.4% | -7.0% | -0.1% | 17.7% | |
| Prestwick-P3 | co | B | 23 | DMSO | -2.1% | -6.5% | -6.7% | 14.2% | |
| Prestwick-P3 | co | C | 23 | DMSO | -0.2% | -0.5% | -9.1% | 6.8% | |
| Prestwick-P3 | co | D | 23 | DMSO | 1.2% | -4.9% | -12.3% | 5.3% | |
| Prestwick-P3 | co | E | 23 | DMSO | 1.4% | -1.3% | -15.9% | 5.0% | |
| Prestwick-P3 | co | F | 23 | DMSO | 2.9% | -3.7% | 1.1% | 4.8% | |
| Prestwick-P3 | co | G | 23 | DMSO | -1.9% | -0.1% | -16.6% | 8.8% | |
| Prestwick-P3 | co | H | 23 | DMSO | -2.7% | -1.8% | -15.2% | 1.1% | |
| Prestwick-P3 | co | I | 23 | DMSO | -1.8% | 1.6% | -6.5% | -0.1% | |
| Prestwick-P3 | co | J | 23 | DMSO | 1.2% | -0.2% | -6.6% | 8.0% | |
| Prestwick-P3 | co | K | 23 | DMSO | 1.6% | -0.7% | -2.9% | 10.3% | |
| Prestwick-P3 | co | L | 23 | DMSO | -0.3% | 0.1% | -3.8% | 8.3% | |
| Prestwick-P3 | co | M | 23 | DMSO | -0.5% | 1.0% | -5.5% | 13.4% | |
| Prestwick-P3 | co | N | 23 | DMSO | -1.0% | 1.2% | -7.3% | 9.9% | |
| Prestwick-P3 | co | O | 23 | DMSO | -4.3% | 4.4% | -4.4% | 12.1% | |
| Prestwick-P3 | co | P | 23 | DMSO | 0.5% | 1.2% | 7.1% | 10.7% | |
| Prestwick-P3 | co | A | 24 | noPFF | -4.3% | -2.0% | -99.9% | 14.6% | |
| Prestwick-P3 | co | B | 24 | noPFF | -0.1% | -6.0% | -99.7% | 11.2% | |
| Prestwick-P3 | co | C | 24 | Ceph4 | -26.1% | 63.5% | -100.0% | -48.1% | |
| Prestwick-P3 | co | D | 24 | noPFF | -0.2% | -1.3% | -99.9% | 7.6% | |
| Prestwick-P3 | co | E | 24 | noPFF | -1.5% | -3.3% | -100.0% | 8.4% | |
| Prestwick-P3 | co | F | 24 | noPFF | -1.7% | -2.6% | -100.0% | 17.3% | |
| Prestwick-P3 | co | G | 24 | noPFF | -3.7% | -3.5% | -100.0% | 23.0% | |
| Prestwick-P3 | co | H | 24 | noPFF | -2.9% | -2.2% | -100.0% | 19.9% | |
| Prestwick-P3 | co | I | 24 | noPFF | -0.5% | -3.7% | -100.0% | 15.7% | |
| Prestwick-P3 | co | J | 24 | noPFF | -5.0% | -3.4% | -100.0% | 20.9% | |

| | | | | | | | | | |
|---|---|---|---|---|---|---|---|---|---|
| Prestwick-P3 | co | K | 24 | noPFF | -2.5% | 3.2% | -100.0% | 19.3% | |
| Prestwick-P3 | co | L | 24 | noPFF | -0.4% | 0.0% | -100.0% | 7.8% | |
| Prestwick-P3 | co | M | 24 | noPFF | -2.4% | 4.3% | -100.0% | 23.4% | |
| Prestwick-P3 | co | N | 24 | noPFF | 0.2% | 4.0% | -100.0% | 1.1% | |
| Prestwick-P3 | co | O | 24 | noPFF | -0.3% | 2.1% | -100.0% | 2.8% | |
| Prestwick-P3 | co | P | 24 | noPFF | -5.7% | -1.3% | -100.0% | 16.4% | |
| Prestwick-P4 | co | A | 1 | DMSO | -1.3% | -6.3% | 3.1% | -13.3% | |
| Prestwick-P4 | co | B | 1 | DMSO | 0.3% | -1.5% | 16.1% | -10.2% | |
| Prestwick-P4 | co | C | 1 | DMSO | -0.3% | -5.5% | 5.3% | -6.2% | |
| Prestwick-P4 | co | D | 1 | DMSO | 0.8% | 1.7% | 13.2% | -7.8% | |
| Prestwick-P4 | co | E | 1 | DMSO | -1.2% | 0.0% | 7.2% | -7.1% | |
| Prestwick-P4 | co | F | 1 | DMSO | 0.2% | -2.0% | 14.4% | 0.3% | |
| Prestwick-P4 | co | G | 1 | DMSO | 0.4% | -5.5% | -15.2% | -6.4% | |
| Prestwick-P4 | co | H | 1 | DMSO | 1.7% | 4.7% | -7.1% | -15.7% | |
| Prestwick-P4 | co | I | 1 | DMSO | 0.7% | -5.8% | -9.8% | -2.3% | |
| Prestwick-P4 | co | J | 1 | DMSO | 2.8% | 0.9% | 2.8% | -7.8% | |
| Prestwick-P4 | co | K | 1 | DMSO | 2.8% | -6.3% | -10.7% | -2.1% | |
| Prestwick-P4 | co | L | 1 | DMSO | 0.4% | 3.7% | -6.2% | 1.4% | |
| Prestwick-P4 | co | M | 1 | DMSO | -0.1% | -3.8% | 20.7% | 3.5% | |
| Prestwick-P4 | co | N | 1 | DMSO | 3.5% | -1.4% | -7.9% | 1.6% | |
| Prestwick-P4 | co | O | 1 | DMSO | -4.6% | 5.8% | -23.3% | 6.7% | |
| Prestwick-P4 | co | P | 1 | DMSO | -0.4% | 9.4% | -8.1% | 4.9% | |
| Prestwick-P4 | co | A | 2 | DMSO | -2.3% | -6.1% | 3.8% | -6.1% | |
| Prestwick-P4 | co | B | 2 | DMSO | 0.6% | -0.5% | 22.0% | -7.0% | |
| Prestwick-P4 | co | C | 2 | DMSO | -0.1% | -8.3% | 19.4% | -5.7% | |
| Prestwick-P4 | co | D | 2 | DMSO | 0.0% | 1.9% | 7.2% | -2.6% | |
| Prestwick-P4 | co | E | 2 | DMSO | -1.0% | -3.8% | 5.1% | -6.1% | |
| Prestwick-P4 | co | F | 2 | DMSO | 1.7% | 5.5% | 0.3% | -14.2% | |
| Prestwick-P4 | co | G | 2 | DMSO | 0.0% | -3.4% | -7.2% | -3.5% | |
| Prestwick-P4 | co | H | 2 | DMSO | 2.5% | 3.6% | 11.5% | -9.7% | |
| Prestwick-P4 | co | I | 2 | Til2 | -8.6% | 18.4% | -83.2% | -1.2% | |
| Prestwick-P4 | co | J | 2 | Til2 | -5.7% | 21.4% | -83.0% | -7.3% | |
| Prestwick-P4 | co | K | 2 | Til0.5 | 0.1% | 2.6% | -43.1% | 6.0% | |
| Prestwick-P4 | co | L | 2 | Til0.5 | 1.4% | 3.2% | -34.4% | 1.3% | |
| Prestwick-P4 | co | M | 2 | Til0.13 | 0.4% | -2.8% | -19.8% | 0.7% | |
| Prestwick-P4 | co | N | 2 | Til0.13 | 1.6% | 5.7% | -10.0% | -2.9% | |
| Prestwick-P4 | co | O | 2 | DMSO | -0.7% | 2.4% | -1.1% | 1.0% | |
| Prestwick-P4 | co | P | 2 | DMSO | -1.6% | 11.8% | -6.6% | 2.4% | |
| Prestwick-P4 | 913 | A | 3 | Dolasetron mesilate | 4.0% | -9.3% | 12.6% | -17.7% | 115956-13-3 |
| Prestwick-P4 | 914 | B | 3 | Clioquinol | -14.2% | 56.6% | -30.8% | -30.5% | 130-26-7 |
| Prestwick-P4 | 915 | C | 3 | Aminohippuric acid | 0.1% | -2.7% | 11.6% | -1.5% | 61-78-9 |
| Prestwick-P4 | 916 | D | 3 | N-Acetyl-L-leucine | 1.2% | 2.8% | 13.9% | -3.4% | 1188-21-2 |
| Prestwick-P4 | 917 | E | 3 | Urapidil hydrochloride | -1.4% | 0.3% | -7.3% | -7.6% | 64887-14-5 |
| Prestwick-P4 | 918 | F | 3 | Fluspirilen | -5.8% | 26.3% | -51.8% | -19.6% | 1841-19-6 |
| Prestwick-P4 | 919 | G | 3 | Lorglumide sodium salt | -0.4% | 0.4% | -8.7% | 3.6% | 97964-56-2 |
| Prestwick-P4 | 920 | H | 3 | Nitrendipine | 2.5% | 2.8% | -22.1% | -9.0% | 39562-70-4 |
| Prestwick-P4 | 921 | I | 3 | Thonzonium bromide | -21.9% | 66.5% | -96.1% | -15.0% | 553-08-2 |
| Prestwick-P4 | 922 | J | 3 | Idazoxan hydrochloride | 5.1% | 4.2% | -5.5% | -0.5% | 79944-56-2 |
| Prestwick-P4 | 923 | K | 3 | Paliperidone | 1.4% | -3.7% | -3.9% | 3.2% | 144598-75-4 |
| Prestwick-P4 | 924 | L | 3 | Bephenium hydroxynaphthoate | 3.6% | 2.7% | 10.1% | 1.2% | 3818-50-6 |
| Prestwick-P4 | 925 | M | 3 | Salmeterol | 1.3% | -1.5% | -3.5% | 1.5% | 89365-50-4 |
| Prestwick-P4 | 926 | N | 3 | Altretamine | 1.1% | 3.1% | 22.1% | 3.7% | 645-05-6 |
| Prestwick-P4 | 927 | O | 3 | Florfenicol | 1.0% | -3.2% | 11.6% | -3.3% | 73231-34-2 |
| Prestwick-P4 | 928 | P | 3 | Megestrol acetate | -0.2% | 10.0% | -6.7% | -7.6% | 595-33-5 |
| Prestwick-P4 | 929 | A | 4 | Oxybenzone | 0.0% | 0.9% | -7.6% | -22.8% | 131-57-7 |
| Prestwick-P4 | 930 | B | 4 | Promethazine hydrochloride | 7.7% | -0.9% | 80.1% | -0.9% | 58-33-3 |
| Prestwick-P4 | 931 | C | 4 | Pipemidic acid | 1.3% | -4.8% | 3.1% | -8.2% | 51940-44-4 |
| Prestwick-P4 | 932 | D | 4 | Dioxybenzone | 2.6% | 4.2% | 2.7% | -0.8% | 131-53-3 |
| Prestwick-P4 | 933 | E | 4 | (S,+) Ibuprofen | -1.4% | -6.8% | -14.0% | -14.7% | 51146-56-6 |
| Prestwick-P4 | 934 | F | 4 | Ethynodiol diacetate | 1.8% | 6.9% | 3.8% | -1.2% | 297-76-7 |
| Prestwick-P4 | 935 | G | 4 | Flurbiprofen | 0.3% | -0.1% | -10.7% | -4.6% | 5104-49-4 |
| Prestwick-P4 | 936 | H | 4 | Nimodipine | -1.5% | 11.6% | -30.1% | -20.1% | 66085-59-4 |
| Prestwick-P4 | 937 | I | 4 | Quinapril hydrochloride | 0.2% | -1.1% | -7.1% | -8.3% | 82586-55-8 |
| Prestwick-P4 | 938 | J | 4 | Nilutamide | 3.9% | 1.6% | 13.7% | -2.9% | 63612-50-0 |
| Prestwick-P4 | 939 | K | 4 | Dehydroisoandosterone 3-acetate | -11.3% | 37.7% | -44.8% | -28.8% | 853-23-6 |
| Prestwick-P4 | 940 | L | 4 | Benserazide hydrochloride | 3.6% | -1.1% | 0.5% | 4.1% | 14919-77-8 |
| Prestwick-P4 | 941 | M | 4 | Prazosin hydrochloride | 2.1% | 2.1% | -35.9% | -10.8% | 19237-84-4 |
| Prestwick-P4 | 942 | N | 4 | Timolol maleate salt | 0.7% | 1.9% | 11.5% | 4.3% | 26921-17-5 |
| Prestwick-P4 | 943 | O | 4 | Deoxycorticosterone | -0.2% | -3.1% | 6.6% | -5.5% | 64-85-7 |
| Prestwick-P4 | 944 | P | 4 | Urosiol | 2.9% | 1.7% | 11.4% | -5.9% | 128-13-2 |
| Prestwick-P4 | 945 | A | 5 | Diacerein | 2.6% | -1.1% | 6.0% | -13.4% | 13739-02-1 |
| Prestwick-P4 | 946 | B | 5 | Esmolol hydrochloride | 3.1% | -0.4% | -3.7% | 1.6% | 81161-17-3 |

| | | | | | | | | | |
|---|---|---|---|---|---|---|---|---|---|
| Prestwick-P4 | 947 | C | 5 | Adrenosterone | 3.2% | 0.2% | -2.2% | -15.2% | 382-45-6 |
| Prestwick-P4 | 948 | D | 5 | Methylatropine nitrate | 1.6% | 2.6% | 0.4% | -7.9% | 52-88-0 |
| Prestwick-P4 | 949 | E | 5 | Nabumetone | 2.6% | 3.8% | -6.3% | -9.3% | 42924-53-8 |
| Prestwick-P4 | 950 | F | 5 | Nisoxetine hydrochloride | 2.0% | -1.7% | 13.8% | 4.1% | 57754-86-6 |
| Prestwick-P4 | 951 | G | 5 | Bacitracin | 1.9% | -1.1% | -6.7% | -11.5% | 1405-87-4 |
| Prestwick-P4 | 952 | H | 5 | Gemifloxacin mesylate | 3.5% | 2.1% | 3.8% | 6.6% | 204519-65-3 |
| Prestwick-P4 | 953 | I | 5 | Ketorolac tromethamine | -0.1% | 2.6% | -12.3% | -8.8% | 74103-07-4 |
| Prestwick-P4 | 954 | J | 5 | Protriptyline hydrochloride | 5.4% | 3.3% | -2.8% | -3.7% | 1225-55-4 |
| Prestwick-P4 | 955 | K | 5 | Iodipamide | 3.7% | -0.9% | -6.5% | -10.4% | 606-17-7 |
| Prestwick-P4 | 956 | L | 5 | Allopurinol | 4.6% | -0.8% | 9.2% | 1.1% | 315-30-0 |
| Prestwick-P4 | 957 | M | 5 | Octopamine hydrochloride | 4.5% | -2.5% | -2.1% | -16.6% | 770-05-8 |
| Prestwick-P4 | 958 | N | 5 | Stavudine | 1.7% | 0.5% | 11.2% | -1.5% | 3056-17-5 |
| Prestwick-P4 | 959 | O | 5 | Proparacaine hydrochloride | 2.0% | 5.4% | -8.3% | -4.5% | 5875-06-9 |
| Prestwick-P4 | 960 | P | 5 | Aminocaproic acid | 1.1% | 5.9% | 11.9% | -2.0% | 60-32-2 |
| Prestwick-P4 | 961 | A | 6 | Denatonium benzoate | 0.6% | -1.2% | -8.7% | -19.7% | 3734-33-6 |
| Prestwick-P4 | 962 | B | 6 | Canrenone | 0.6% | 0.6% | 0.3% | 12.2% | 976-71-6 |
| Prestwick-P4 | 963 | C | 6 | Remoxipride hydrochloride | 1.7% | -2.9% | -15.1% | -20.7% | 73220-03-8 |
| Prestwick-P4 | 964 | D | 6 | Rosuvastatin | 0.3% | 9.1% | 9.8% | -5.3% | 287714-41-4 |
| Prestwick-P4 | 965 | E | 6 | Nitrocaramiphen hydrochloride | -1.8% | 8.0% | -17.6% | -18.8% | 98636-73-8 |
| Prestwick-P4 | 966 | F | 6 | Nandrolone | 2.4% | 3.8% | 0.6% | -3.9% | 434-22-0 |
| Prestwick-P4 | 967 | G | 6 | Gliquidone | 1.4% | 4.9% | -13.7% | -13.8% | 33342-05-1 |
| Prestwick-P4 | 968 | H | 6 | Pizotifen malate | 8.0% | 0.1% | 54.4% | 2.5% | 5189-11-7 |
| Prestwick-P4 | 969 | I | 6 | Alfadolone acetate | -0.4% | 3.2% | -28.2% | -10.3% | 23930-37-2 |
| Prestwick-P4 | 970 | J | 6 | Alfaxalone | 4.2% | -0.2% | -6.0% | -3.9% | 23930-19-0 |
| Prestwick-P4 | 971 | K | 6 | Flucloxacillin sodium | 3.3% | 7.3% | -5.9% | -14.0% | 1847-24-1 |
| Prestwick-P4 | 972 | L | 6 | Trapidil | 2.8% | 0.4% | 16.1% | 3.4% | 15421-84-8 |
| Prestwick-P4 | 973 | M | 6 | Isradipine | -11.8% | 31.8% | -54.3% | -56.7% | 75695-93-1 |
| Prestwick-P4 | 974 | N | 6 | Nifekalant | 3.6% | 2.5% | 6.8% | -1.1% | 130636-43-0 |
| Prestwick-P4 | 975 | O | 6 | Halofantrine hydrochloride | -17.2% | 44.4% | -98.6% | -57.6% | 36167-63-2 |
| Prestwick-P4 | 976 | P | 6 | Articaine hydrochloride | 0.2% | 4.1% | 12.0% | 2.5% | 23964-57-0 |
| Prestwick-P4 | 977 | A | 7 | Enilconazole | -4.6% | 6.6% | -28.3% | -10.6% | 35554-44-0 |
| Prestwick-P4 | 978 | B | 7 | Methacycline hydrochloride | -0.5% | 1.5% | -17.3% | 11.1% | 3963-95-9 |
| Prestwick-P4 | 979 | C | 7 | Pirlindole mesylate | 1.9% | 2.0% | -11.0% | -16.3% | 60762-57-4 |
| Prestwick-P4 | 980 | D | 7 | Pronethalol hydrochloride | 1.1% | 5.4% | 2.1% | -5.4% | 51-02-5 |
| Prestwick-P4 | 981 | E | 7 | Dimaprit dihydrochloride | 4.4% | 5.9% | 0.5% | -19.6% | 23256-33-9 |
| Prestwick-P4 | 982 | F | 7 | Oxfendazole | 0.9% | -3.1% | -3.3% | -1.2% | 53716-50-0 |
| Prestwick-P4 | 983 | G | 7 | Ribavirin | 3.2% | -0.8% | -14.8% | -6.7% | 36791-04-5 |
| Prestwick-P4 | 984 | H | 7 | Cyclopenthiazide | 1.7% | 2.0% | 2.2% | -3.1% | 742-20-1 |
| Prestwick-P4 | 985 | I | 7 | Azapropazone | 2.6% | 0.8% | -11.3% | -9.6% | 13539-59-8 |
| Prestwick-P4 | 986 | J | 7 | Meptazinol hydrochloride | 3.9% | 1.0% | 0.3% | -4.5% | 59263-76-2 |
| Prestwick-P4 | 987 | K | 7 | Deptropine citrate | 6.8% | 1.1% | 10.0% | -6.2% | 2169-75-7 |
| Prestwick-P4 | 988 | L | 7 | Sertraline | -2.8% | 20.4% | -32.9% | -13.5% | 79617-96-2 |
| Prestwick-P4 | 989 | M | 7 | Isometheptene mucate | 2.2% | 3.0% | -5.3% | -3.7% | 7492-31-1 |
| Prestwick-P4 | 990 | N | 7 | Nifurtimox | 3.7% | -4.2% | 33.0% | -10.4% | 23256-30-6 |
| Prestwick-P4 | 991 | O | 7 | Nomegestrol acetate | 4.5% | 3.6% | -30.0% | -14.5% | 58652-20-3 |
| Prestwick-P4 | 992 | P | 7 | Pancuronium bromide | 0.9% | 8.2% | 11.7% | -7.5% | 15500-66-0 |
| Prestwick-P4 | 993 | A | 8 | Floxuridine | -1.9% | -5.5% | -13.1% | -6.1% | 50-91-9 |
| Prestwick-P4 | 994 | B | 8 | Sotalol hydrochloride | 3.0% | -4.5% | -2.1% | 15.8% | 959-24-0 |
| Prestwick-P4 | 995 | C | 8 | Naftopidil dihydrochloride | 0.6% | 1.3% | -29.7% | -25.1% | 57149-08-3 |
| Prestwick-P4 | 996 | D | 8 | Tracazolate hydrochloride | -1.5% | 14.9% | -1.9% | -7.0% | 41094-88-6 |
| Prestwick-P4 | 997 | E | 8 | Guaiacol | 1.7% | -5.5% | -14.8% | -6.7% | 90-05-1 |
| Prestwick-P4 | 998 | F | 8 | Capecitabine | 2.7% | 1.9% | 11.3% | 0.5% | 154361-50-9 |
| Prestwick-P4 | 999 | G | 8 | Fluvoxamine maleate | 1.6% | 2.4% | 3.9% | -10.6% | 61718-82-9 |
| Prestwick-P4 | 1000 | H | 8 | Prothionamide | 2.8% | -3.0% | 5.1% | 3.0% | 14222-60-7 |
| Prestwick-P4 | 1001 | I | 8 | Apramycin | 1.0% | -2.5% | -7.8% | -3.5% | 37321-09-8 |
| Prestwick-P4 | 1002 | J | 8 | Darunavir | 3.8% | 2.0% | -3.2% | -2.6% | 635728-49-3 |
| Prestwick-P4 | 1003 | K | 8 | Ethamsylate diethylammonium salt | 5.2% | -2.0% | 5.8% | -3.9% | 2624-44-4 |
| Prestwick-P4 | 1004 | L | 8 | Moxonidine | 6.3% | -3.2% | 6.6% | 0.4% | 75438-57-2 |
| Prestwick-P4 | 1005 | M | 8 | Letrozole | 3.9% | -4.8% | 1.0% | -8.2% | 112809-51-5 |
| Prestwick-P4 | 1006 | N | 8 | Levofloxacin | 4.5% | 0.2% | 3.5% | 1.3% | 100986-85-4 |
| Prestwick-P4 | 1007 | O | 8 | Molindone hydrochloride | 6.9% | -5.8% | -18.2% | -11.9% | 15622-65-8 |
| Prestwick-P4 | 1008 | P | 8 | Alcuronium chloride | 1.9% | 1.8% | -7.2% | -1.6% | 15180-03-7 |
| Prestwick-P4 | 1009 | A | 9 | Gestrinone | 0.6% | -1.8% | -10.8% | 5.2% | 16320-04-0 |
| Prestwick-P4 | 1010 | B | 9 | Decamethonium bromide | 1.5% | 3.6% | 4.2% | 2.7% | 541-22-0 |
| Prestwick-P4 | 1011 | C | 9 | Zardaverine | -0.3% | 0.6% | -15.3% | -4.5% | 101975-10-4 |
| Prestwick-P4 | 1012 | D | 9 | Memantine hydrochloride | -1.2% | 8.6% | 4.5% | 3.7% | 41100-52-1 |
| Prestwick-P4 | 1013 | E | 9 | Pramipexole dihydrochloride | 2.5% | 1.9% | -10.6% | -2.5% | 104632-25-9 |
| Prestwick-P4 | 1014 | F | 9 | Norgestimate | -3.2% | 17.2% | -12.9% | -11.7% | 35189-28-7 |
| Prestwick-P4 | 1015 | G | 9 | Fluticasone propionate | -0.3% | 5.9% | -14.2% | -17.5% | 80474-14-2 |
| Prestwick-P4 | 1016 | H | 9 | Zuclopenthixol dihydrochloride | -0.8% | 7.7% | -31.4% | -1.2% | 633-59-0 |
| Prestwick-P4 | 1017 | I | 9 | Fursultiamine hydrochloride | 0.8% | 0.2% | -4.5% | 6.6% | 2105-43-3 |
| Prestwick-P4 | 1018 | J | 9 | Gabexate mesilate | 3.4% | 2.9% | 7.4% | -6.6% | 56974-61-9 |

| | | | | | | | | | |
|---|---|---|---|---|---|---|---|---|---|
| Prestwick-P4 | 1019 | K | 9 | Etilefrine hydrochloride | 2.1% | 1.0% | 15.3% | 5.2% | 534-87-2 |
| Prestwick-P4 | 1020 | L | 9 | Alprostadil | 1.5% | -1.1% | -0.5% | -0.7% | 745-65-3 |
| Prestwick-P4 | 1021 | M | 9 | Tocainide hydrochloride | 0.7% | -1.5% | -3.9% | 4.1% | 71395-14-7 |
| Prestwick-P4 | 1022 | N | 9 | Benzathine benzylpenicillin | 2.0% | 5.5% | 7.4% | 0.9% | 5928-84-7 |
| Prestwick-P4 | 1023 | O | 9 | Zalcitabine | 1.4% | -3.6% | 0.2% | -11.0% | 7481-89-2 |
| Prestwick-P4 | 1024 | P | 9 | Methyldopate hydrochloride | -1.1% | 0.8% | -5.4% | -0.7% | 2508-79-4 |
| Prestwick-P4 | 1025 | A | 10 | Darifenacin hydrobromide | -4.6% | -0.6% | -38.5% | 7.3% | 133099-07-7 |
| Prestwick-P4 | 1026 | B | 10 | Indatraline hydrochloride | -4.0% | 17.6% | -25.5% | 6.2% | 86939-10-8 |
| Prestwick-P4 | 1027 | C | 10 | Ozagrel hydrochloride | 0.1% | -3.1% | -11.2% | -10.1% | 78712-43-3 |
| Prestwick-P4 | 1028 | D | 10 | Piribedil | -0.3% | 3.6% | 0.6% | 7.7% | 3605-01-4 |
| Prestwick-P4 | 1029 | E | 10 | Chlormadinone acetate | -1.2% | -2.2% | -20.5% | -7.9% | 302-22-7 |
| Prestwick-P4 | 1030 | F | 10 | Phenylbutazone | 0.9% | 4.6% | 4.1% | 2.5% | 50-33-9 |
| Prestwick-P4 | 1031 | G | 10 | Proguanil hydrochloride | 2.0% | -1.4% | -6.9% | 1.3% | 637-32-1 |
| Prestwick-P4 | 1032 | H | 10 | Lymecycline | -0.3% | 0.1% | 7.9% | -0.7% | 992-21-2 |
| Prestwick-P4 | 1033 | I | 10 | Pivampicillin | -0.2% | -3.0% | -45.1% | -6.5% | 33817-20-8 |
| Prestwick-P4 | 1034 | J | 10 | Lodoxamide | 0.6% | 0.4% | 1.3% | -3.8% | 53882-12-5 |
| Prestwick-P4 | 1035 | K | 10 | Tribenoside | -3.1% | 9.4% | -20.4% | -6.1% | 10310-32-4 |
| Prestwick-P4 | 1036 | L | 10 | Rimexolone | 1.2% | 3.1% | -4.5% | -13.7% | 49697-38-3 |
| Prestwick-P4 | 1037 | M | 10 | Risperidone | -0.7% | -2.5% | -16.5% | -5.0% | 106266-06-2 |
| Prestwick-P4 | 1038 | N | 10 | Torsemide | 2.2% | 4.8% | -0.7% | -19.9% | 56211-40-6 |
| Prestwick-P4 | 1039 | O | 10 | Levocabastine hydrochloride | -0.3% | -4.2% | 0.5% | -15.6% | 79547-78-7 |
| Prestwick-P4 | 1040 | P | 10 | Pyrvinium pamoate | -28.9% | 68.9% | -82.2% | -65.0% | 3546-41-6 |
| Prestwick-P4 | 1041 | A | 11 | Etomidate | 0.7% | -0.9% | -10.3% | -3.7% | 33125-97-2 |
| Prestwick-P4 | 1042 | B | 11 | Tridihexethyl chloride | 0.5% | 3.9% | -2.3% | -5.1% | 4310-35-4 |
| Prestwick-P4 | 1043 | C | 11 | Moricizine hydrochloride | 1.3% | -6.6% | -5.5% | -1.7% | 31883-05-3 |
| Prestwick-P4 | 1044 | D | 11 | Iopanoic acid | 0.9% | 0.8% | 2.7% | 1.0% | 96-83-3 |
| Prestwick-P4 | 1045 | E | 11 | Phensuximide | 2.0% | 1.7% | -10.3% | -12.1% | 86-34-0 |
| Prestwick-P4 | 1046 | F | 11 | Ioxaglic acid | 1.2% | -4.6% | -3.3% | 8.1% | 59017-64-0 |
| Prestwick-P4 | 1047 | G | 11 | Imidurea | -0.2% | -1.0% | -7.6% | -3.2% | 39236-46-9 |
| Prestwick-P4 | 1048 | H | 11 | Lansoprazole | 0.4% | 2.0% | 7.4% | 1.6% | 103577-45-3 |
| Prestwick-P4 | 1049 | I | 11 | (S)-(-)-Propranolol hydrochloride | 1.4% | -2.3% | -11.1% | 8.8% | 4199-10-4 |
| Prestwick-P4 | 1050 | J | 11 | (-) Eseroline fumarate salt | 4.3% | 6.7% | -51.4% | -5.9% | 104015-29-4 |
| Prestwick-P4 | 1051 | K | 11 | Spaglumic acid | 2.3% | -0.7% | -0.6% | 3.7% | 3106-85-2 |
| Prestwick-P4 | 1052 | L | 11 | Ranolazine | 3.8% | -0.2% | 7.4% | -1.3% | 95635-55-5 |
| Prestwick-P4 | 1053 | M | 11 | Perindopril | 3.7% | -3.3% | -11.0% | -3.6% | 82834-16-0 |
| Prestwick-P4 | 1054 | N | 11 | Fexofenadine hydrochloride | 2.5% | 3.9% | -4.0% | -12.3% | 153439-40-8 |
| Prestwick-P4 | 1055 | O | 11 | Mecamylamine hydrochloride | 2.7% | -2.8% | 0.0% | -1.1% | 826-39-1 |
| Prestwick-P4 | 1056 | P | 11 | Procarbazine hydrochloride | 0.2% | 1.1% | 0.6% | -17.4% | 366-70-1 |
| Prestwick-P4 | 1057 | A | 12 | Penbutolol sulfate | -1.1% | 6.9% | -13.0% | -5.0% | 38363-32-5 |
| Prestwick-P4 | 1058 | B | 12 | Prednicarbate | -0.6% | 6.0% | 4.8% | 0.8% | 73771-04-7 |
| Prestwick-P4 | 1059 | C | 12 | Pivmecillinam hydrochloride | -1.1% | -2.1% | -16.5% | 0.9% | 32887-03-9 |
| Prestwick-P4 | 1060 | D | 12 | Levopropoxyphene napsylate | 0.8% | 0.7% | -0.4% | -1.1% | 5714-90-9 |
| Prestwick-P4 | 1061 | E | 12 | Naftifine hydrochloride | 2.0% | -1.3% | -16.0% | -6.3% | 65473-14-5 |
| Prestwick-P4 | 1062 | F | 12 | Meprylcaine hydrochloride | 1.5% | 4.9% | -7.5% | 3.1% | 956-03-6 |
| Prestwick-P4 | 1063 | G | 12 | Bethanechol chloride | 0.7% | 4.2% | -8.2% | -26.2% | 590-63-6 |
| Prestwick-P4 | 1064 | H | 12 | Cyproterone acetate | -6.1% | 18.3% | -32.2% | -42.7% | 427-51-0 |
| Prestwick-P4 | 1065 | I | 12 | Isosorbide mononitrate | 2.4% | -1.1% | -12.4% | -2.5% | 16051-77-7 |
| Prestwick-P4 | 1066 | J | 12 | Levalbuterol hydrochloride | -0.2% | 0.9% | -0.8% | -1.0% | 50293-90-8 |
| Prestwick-P4 | 1067 | K | 12 | Misoprostol | 1.3% | -1.4% | -3.0% | 1.3% | 59122-46-2 |
| Prestwick-P4 | 1068 | L | 12 | Sulfadoxine | 2.6% | 1.7% | 10.0% | 0.8% | 2447-57-6 |
| Prestwick-P4 | 1069 | M | 12 | 4-aminosalicylic acid | -1.4% | 1.2% | -11.4% | -3.9% | 65-49-6 |
| Prestwick-P4 | 1070 | N | 12 | Clonixin lysinate | 0.9% | 3.3% | 3.4% | 3.5% | 55837-30-4 |
| Prestwick-P4 | 1071 | O | 12 | Viomycin sulfate | 0.5% | -2.6% | -1.9% | -7.7% | 37883-00-4 |
| Prestwick-P4 | 1072 | P | 12 | Saquinavir mesylate | -6.1% | 12.5% | -16.9% | -25.2% | 149845-06-7 |
| Prestwick-P4 | 1073 | A | 13 | Sertaconazole nitrate | -16.8% | 41.9% | -61.1% | -37.9% | 99592-39-9 |
| Prestwick-P4 | 1074 | B | 13 | Repaglinide | -1.6% | 1.9% | -5.1% | -4.2% | 135062-02-1 |
| Prestwick-P4 | 1075 | C | 13 | Piperidolate hydrochloride | -1.4% | 5.9% | -12.8% | -1.7% | 129-77-1 |
| Prestwick-P4 | 1076 | D | 13 | Trifluridine | 2.7% | 3.1% | -5.5% | 0.2% | 70-00-8 |
| Prestwick-P4 | 1077 | E | 13 | Milrinone | 1.8% | -1.0% | -3.6% | -1.4% | 78415-72-2 |
| Prestwick-P4 | 1078 | F | 13 | Methantheline bromide | -1.9% | 5.2% | 15.6% | -2.9% | 53-46-3 |
| Prestwick-P4 | 1079 | G | 13 | (R) Propranolol hydrochloride | -0.7% | 2.9% | -8.6% | -8.0% | 13071-11-9 |
| Prestwick-P4 | 1080 | H | 13 | Ciprofibrate | 0.1% | 1.6% | 2.3% | 5.9% | 52214-84-3 |
| Prestwick-P4 | 1081 | I | 13 | Topiramate | 1.0% | 1.5% | -18.5% | -9.1% | 97240-79-4 |
| Prestwick-P4 | 1082 | J | 13 | D-cycloserine | 0.0% | 0.2% | -6.9% | 3.5% | 68-41-7 |
| Prestwick-P4 | 1083 | K | 13 | Cyclopentolate hydrochloride | 0.4% | 0.3% | 2.7% | 18.7% | 5870-29-1 |
| Prestwick-P4 | 1084 | L | 13 | Estriol | -0.2% | 0.7% | -4.3% | 1.9% | 50-27-1 |
| Prestwick-P4 | 1085 | M | 13 | Verteporfin | -23.0% | -16.9% | -89.2% | 61.9% | 129497-78-5 |
| Prestwick-P4 | 1086 | N | 13 | Meropenem | -1.0% | 5.6% | -2.9% | -3.8% | 96036-03-2 |
| Prestwick-P4 | 1087 | O | 13 | Ronidazole | -1.6% | -2.7% | -3.2% | -5.9% | 7681-76-7 |
| Prestwick-P4 | 1088 | P | 13 | Dorzolamide hydrochloride | -0.9% | 6.2% | 5.5% | -9.4% | 130693-82-2 |
| Prestwick-P4 | 1089 | A | 13 | Piretanide | -1.4% | -0.3% | -18.7% | 6.9% | 55837-27-9 |
| Prestwick-P4 | 1090 | B | 14 | Piperacetazine | -1.3% | 4.8% | -30.1% | 13.7% | 3819-00-9 |

| | | | | | | | | | |
|---|---|---|---|---|---|---|---|---|---|
| Prestwick-P4 | 1091 | C | 14 | Oxprenolol hydrochloride | 0.4% | 1.1% | 5.7% | -5.9% | 6452-73-9 |
| Prestwick-P4 | 1092 | D | 14 | Ondansetron hydrochloride | -0.4% | 4.8% | -5.0% | 8.6% | 103639-04-9 |
| Prestwick-P4 | 1093 | E | 14 | Ticarcillin sodium | 2.2% | -5.8% | -5.7% | 6.1% | 74682-62-5 |
| Prestwick-P4 | 1094 | F | 14 | Thiethylperazine maleate | -6.6% | 36.9% | -77.7% | -19.2% | 1179-69-7 |
| Prestwick-P4 | 1095 | G | 14 | Formestane | 1.3% | 2.6% | 5.1% | 2.4% | 566-48-3 |
| Prestwick-P4 | 1096 | H | 14 | Benzylpenicillin sodium | -0.8% | 4.3% | 1.5% | 5.5% | 69-57-8 |
| Prestwick-P4 | 1097 | I | 14 | Nelarabine | 2.0% | -4.4% | -11.2% | -12.1% | 121032-29-9 |
| Prestwick-P4 | 1098 | J | 14 | Synephrine | 2.3% | 0.3% | -8.5% | -7.8% | 94-07-5 |
| Prestwick-P4 | 1099 | K | 14 | (-) Isoproterenol hydrochloride | 1.7% | -1.3% | -9.3% | -0.2% | 5984-95-2 |
| Prestwick-P4 | 1100 | L | 14 | Sarafloxacin | 3.9% | -1.1% | 9.4% | 4.7% | 98105-99-8 |
| Prestwick-P4 | 1101 | M | 14 | Ramipril | 3.2% | -1.2% | -5.4% | -17.9% | 87333-19-5 |
| Prestwick-P4 | 1102 | N | 14 | Mephenytoin | 3.4% | -1.8% | 5.9% | -5.0% | 50-12-4 |
| Prestwick-P4 | 1103 | O | 14 | Azaperone | -1.7% | -2.5% | -2.1% | -11.7% | 1649-18-9 |
| Prestwick-P4 | 1104 | P | 14 | Cefepime hydrochloride | -0.7% | 4.4% | 8.4% | -7.4% | 123171-59-5 |
| Prestwick-P4 | 1105 | A | 15 | Oxyphenbutazone | -0.4% | -1.4% | 2.1% | -1.4% | 129-20-4 |
| Prestwick-P4 | 1106 | B | 15 | Quinethazone | -0.1% | 1.7% | 3.4% | 15.0% | 73-49-4 |
| Prestwick-P4 | 1107 | C | 15 | Propoxycaine hydrochloride | 2.6% | 0.9% | -0.8% | -0.7% | 550-83-4 |
| Prestwick-P4 | 1108 | D | 15 | Oxaprozin | 0.6% | 5.1% | 7.7% | 5.4% | 21256-18-8 |
| Prestwick-P4 | 1109 | E | 15 | Mesalamine | 0.8% | 3.4% | -1.2% | 2.3% | 89-57-6 |
| Prestwick-P4 | 1110 | F | 15 | Vorinostat | 2.4% | 2.3% | 52.9% | -14.2% | 149647-78-9 |
| Prestwick-P4 | 1111 | G | 15 | Methicillin sodium | 0.3% | 3.0% | -11.5% | -6.5% | 7246-14-2 |
| Prestwick-P4 | 1112 | H | 15 | Methiazole | 3.6% | 16.2% | -18.1% | -32.4% | 108579-67-5 |
| Prestwick-P4 | 1113 | I | 15 | (S,-) Cycloserine | 2.0% | 0.5% | -8.5% | -1.8% | 339-72-0 |
| Prestwick-P4 | 1114 | J | 15 | Homosalate | -27.6% | 73.3% | -81.5% | -63.8% | 118-56-9 |
| Prestwick-P4 | 1115 | K | 15 | Nialamide | 0.8% | 3.9% | 11.4% | 11.5% | 51-12-7 |
| Prestwick-P4 | 1116 | L | 15 | Toltrazuril | -0.7% | 1.2% | -7.7% | 1.7% | 69004-03-1 |
| Prestwick-P4 | 1117 | M | 15 | Rifabutin | 2.1% | 0.7% | -21.7% | -9.9% | 72559-06-9 |
| Prestwick-P4 | 1118 | N | 15 | Parbendazole | -3.7% | 28.3% | -42.2% | -45.7% | 14255-87-9 |
| Prestwick-P4 | 1119 | O | 15 | Clocortolone pivalate | -10.4% | 22.5% | -25.2% | -17.7% | 34097-16-0 |
| Prestwick-P4 | 1120 | P | 15 | Nadifloxacin | -1.9% | 2.3% | -5.2% | -14.2% | 124858-35-1 |
| Prestwick-P4 | 1121 | A | 16 | Buspirone hydrochloride | 0.3% | -3.7% | 1.4% | 7.2% | 33386-08-2 |
| Prestwick-P4 | 1122 | B | 16 | Anastrozole | -0.7% | 1.7% | 2.0% | 5.7% | 120511-73-1 |
| Prestwick-P4 | 1123 | C | 16 | Acetylcysteine | -0.5% | 2.7% | -4.3% | -12.0% | 616-91-1 |
| Prestwick-P4 | 1124 | D | 16 | Melengestrol acetate | -0.6% | 5.9% | -13.1% | 1.2% | 2919-66-6 |
| Prestwick-P4 | 1125 | E | 16 | Famciclovir | 1.6% | -0.3% | 1.7% | 7.3% | 104227-87-4 |
| Prestwick-P4 | 1126 | F | 16 | Dopamine hydrochloride | 0.7% | 1.0% | -6.1% | 6.6% | 62-31-7 |
| Prestwick-P4 | 1127 | G | 16 | 5-fluorouracil | 1.2% | 3.0% | -6.2% | -1.1% | 51-21-8 |
| Prestwick-P4 | 1128 | H | 16 | 2-Mercaptoethanesulfonic acid sodium salt | -0.5% | 6.3% | 2.6% | 3.9% | 19767-45-4 |
| Prestwick-P4 | 1129 | I | 16 | Clofibrate | 3.5% | 1.4% | -20.8% | -10.5% | 637-07-0 |
| Prestwick-P4 | 1130 | J | 16 | Dexrazoxane hydrochloride | 3.1% | 2.4% | -1.1% | -9.5% | 24584-09-6 |
| Prestwick-P4 | 1131 | K | 16 | Topotecan | -4.4% | 20.7% | -27.8% | -46.1% | 123948-87-8 |
| Prestwick-P4 | 1132 | L | 16 | Atorvastatin | -3.4% | 11.4% | -10.0% | 6.8% | 134523-00-5 |
| Prestwick-P4 | 1133 | M | 16 | Gatifloxacin | 0.7% | 2.1% | 2.6% | -9.8% | 112811-59-3 |
| Prestwick-P4 | 1134 | N | 16 | Bosentan | 3.4% | -0.7% | -1.4% | -13.0% | 147536-97-8 |
| Prestwick-P4 | 1135 | O | 16 | Imatinib | -1.0% | 1.8% | -20.7% | -5.4% | 152459-95-5 |
| Prestwick-P4 | 1136 | P | 16 | Moxifloxacin | -2.9% | 1.8% | -1.3% | -4.3% | 151096-09-2 |
| Prestwick-P4 | 1137 | A | 17 | Doxycycline hydrochloride | -1.8% | 2.5% | -16.6% | -9.4% | 10592-13-9 |
| Prestwick-P4 | 1138 | B | 17 | Sulbactam | 0.3% | -1.2% | -6.7% | 10.1% | 68373-14-8 |
| Prestwick-P4 | 1139 | C | 17 | Bromhexine hydrochloride | -20.4% | 49.4% | -68.1% | -55.3% | 611-75-6 |
| Prestwick-P4 | 1140 | D | 17 | Anethole-trithione | 1.3% | 7.0% | 1.3% | -4.9% | 532-11-6 |
| Prestwick-P4 | 1141 | E | 17 | Cefdinir | 2.4% | -1.3% | -11.6% | 9.2% | 91832-40-5 |
| Prestwick-P4 | 1142 | F | 17 | Carprofen | 3.8% | 3.9% | 1.7% | 7.3% | 53716-49-7 |
| Prestwick-P4 | 1143 | G | 17 | Mitotane | -26.0% | 73.8% | -84.0% | -53.6% | 53-19-0 |
| Prestwick-P4 | 1144 | H | 17 | Ambrisentan | 1.6% | -2.1% | -5.4% | -1.0% | 177036-94-1 |
| Prestwick-P4 | 1145 | I | 17 | Aripiprazole | -5.6% | 19.3% | -30.3% | -7.4% | 129722-12-9 |
| Prestwick-P4 | 1146 | J | 17 | Ethinylestradiol | 4.5% | 0.4% | -14.1% | -15.1% | 57-63-6 |
| Prestwick-P4 | 1147 | K | 17 | Azithromycin | 1.8% | 1.3% | -59.7% | 10.3% | 83905-01-5 |
| Prestwick-P4 | 1148 | L | 17 | Ibudilast | 3.9% | -1.9% | 5.2% | -0.6% | 50847-11-5 |
| Prestwick-P4 | 1149 | M | 17 | Gemcitabine | 4.0% | -0.5% | -10.5% | -15.5% | 95058-81-4 |
| Prestwick-P4 | 1150 | N | 17 | Olmesartan | 2.5% | -2.4% | -5.8% | -9.1% | 144689-63-4 |
| Prestwick-P4 | 1151 | O | 17 | Formoterol fumarate | 6.6% | -1.0% | -6.6% | -11.0% | 43229-80-7 |
| Prestwick-P4 | 1152 | P | 17 | Rufloxacin | 0.0% | 2.6% | -8.0% | -14.5% | 101363-10-4 |
| Prestwick-P4 | 1153 | A | 18 | Fleroxacin | -2.2% | 0.5% | -5.4% | 18.6% | 79660-72-3 |
| Prestwick-P4 | 1154 | B | 18 | Clavulanate potassium salt | 0.2% | -1.1% | -0.7% | 8.8% | 58001-44-8 |
| Prestwick-P4 | 1155 | C | 18 | Amcinonide | 1.7% | 5.6% | -17.6% | -5.6% | 51022-69-6 |
| Prestwick-P4 | 1156 | D | 18 | Caffeine | 0.7% | 0.1% | 1.3% | 11.0% | 58-08-2 |
| Prestwick-P4 | 1157 | E | 18 | Celecoxib | -8.6% | 23.6% | -23.1% | -8.2% | 169590-42-5 |
| Prestwick-P4 | 1158 | F | 18 | Candesartan | 1.8% | -1.4% | 4.9% | 0.3% | 139481-59-7 |
| Prestwick-P4 | 1159 | G | 18 | Triclosan | -11.1% | 34.2% | -33.2% | -23.5% | 3380-34-5 |
| Prestwick-P4 | 1160 | H | 18 | Enoxacin | 0.3% | 0.1% | 4.7% | -1.8% | 84294-96-2 |
| Prestwick-P4 | 1161 | I | 18 | Fluocinolone acetonide | 2.0% | 1.7% | -5.4% | -6.4% | 67-73-2 |
| Prestwick-P4 | 1162 | J | 18 | Sparfloxacin | 3.7% | 0.6% | 17.8% | 4.0% | 110871-86-8 |

| | | | | | | | | | |
|---|---|---|---|---|---|---|---|---|---|
| Prestwick-P4 | 1163 | K | 18 | Losartan | 2.3% | 2.4% | -2.0% | 11.9% | 114798-26-4 |
| Prestwick-P4 | 1164 | L | 18 | Benztropine mesylate | 4.6% | 1.3% | 13.3% | 13.2% | 132-17-2 |
| Prestwick-P4 | 1165 | M | 18 | Racepinephrine hydrochloride | 4.5% | 0.7% | -7.3% | 2.2% | 329-63-5 |
| Prestwick-P4 | 1166 | N | 18 | Montelukast | 3.5% | -3.0% | -4.7% | -1.1% | 158966-92-8 |
| Prestwick-P4 | 1167 | O | 18 | Pravastatin | 0.6% | 3.2% | 8.8% | -3.2% | 81093-37-0 |
| Prestwick-P4 | 1168 | P | 18 | Rosiglitazone hydrochloride | -8.2% | 9.4% | -7.9% | 2.5% | 122320-73-4 |
| Prestwick-P4 | 1169 | A | 19 | Valproic acid | -4.3% | 2.8% | -4.6% | 17.3% | 99-66-1 |
| Prestwick-P4 | 1170 | B | 19 | Mepivacaine hydrochloride | -2.2% | 4.1% | -0.3% | 17.2% | 1722-62-9 |
| Prestwick-P4 | 1171 | C | 19 | Carvedilol | 2.5% | 2.1% | -38.2% | -0.2% | 72956-09-3 |
| Prestwick-P4 | 1172 | D | 19 | Methenamine | 2.1% | -0.4% | 4.3% | 8.3% | 100-97-0 |
| Prestwick-P4 | 1173 | E | 19 | Fludarabine | 6.6% | 3.3% | 9.2% | -1.6% | 21679-14-1 |
| Prestwick-P4 | 1174 | F | 19 | Cladribine | 1.2% | 8.4% | 0.4% | -32.2% | 4291-63-8 |
| Prestwick-P4 | 1175 | G | 19 | Olopatadine hydrochloride | 1.8% | 0.8% | -5.2% | 0.2% | 140462-76-6 |
| Prestwick-P4 | 1176 | H | 19 | Granisetron | 0.4% | 3.6% | -4.2% | -4.1% | 109889-09-0 |
| Prestwick-P4 | 1177 | I | 19 | Desloratadine | 1.2% | 1.8% | -18.8% | -3.1% | 100643-71-8 |
| Prestwick-P4 | 1178 | J | 19 | Clarithromycin | 1.1% | 1.2% | -7.2% | 11.3% | 81103-11-9 |
| Prestwick-P4 | 1179 | K | 19 | Vecuronium bromide | 0.6% | 5.9% | -4.6% | 9.8% | 50700-72-6 |
| Prestwick-P4 | 1180 | L | 19 | Telmisartan | 3.0% | 4.6% | 9.7% | 11.8% | 144701-48-4 |
| Prestwick-P4 | 1181 | M | 19 | Docetaxel | 11.5% | 4.8% | -33.5% | -4.1% | 114977-28-5 |
| Prestwick-P4 | 1182 | N | 19 | Cilnidipine | -0.1% | 7.1% | -11.7% | -10.4% | 132203-70-4 |
| Prestwick-P4 | 1183 | O | 19 | Rivastigmine | 1.6% | 0.4% | 9.8% | 5.4% | 123441-03-2 |
| Prestwick-P4 | 1184 | P | 19 | Sildenafil | -0.9% | 6.6% | -2.5% | -8.9% | 139755-83-2 |
| Prestwick-P4 | 1185 | A | 20 | Rifaximin | -3.5% | -4.1% | 4.7% | 28.3% | 80621-81-4 |
| Prestwick-P4 | 1186 | B | 20 | Estradiol Valerate | -20.9% | 63.9% | -57.2% | -43.9% | 979-32-8 |
| Prestwick-P4 | 1187 | C | 20 | Phentermine hydrochloride | 1.1% | -0.1% | 1.4% | 6.2% | 1197-21-3 |
| Prestwick-P4 | 1188 | D | 20 | Diclazuril | -1.5% | 0.9% | -3.5% | -2.1% | 101831-37-2 |
| Prestwick-P4 | 1189 | E | 20 | Vardenafil | 1.4% | 0.3% | -1.0% | -1.6% | 224785-90-4 |
| Prestwick-P4 | 1190 | F | 20 | Fluconazole | -0.5% | 8.2% | -0.7% | -2.2% | 86386-73-4 |
| Prestwick-P4 | 1191 | G | 20 | Anthralin | -3.4% | -0.6% | -39.6% | 4.6% | 1143-38-0 |
| Prestwick-P4 | 1192 | H | 20 | Lamotrigine | 0.1% | 0.2% | 6.1% | 0.3% | 84057-84-1 |
| Prestwick-P4 | 1193 | I | 20 | Tripelennamine hydrochloride | 3.0% | -1.1% | 32.5% | -4.5% | 154-69-8 |
| Prestwick-P4 | 1194 | J | 20 | Tulobuterol hydrochloride | 0.1% | 0.2% | -1.7% | 10.6% | 56776-01-3 |
| Prestwick-P4 | 1195 | K | 20 | Nalmefene hydrochloride | 1.2% | 1.0% | -10.2% | 4.4% | 58895-64-0 |
| Prestwick-P4 | 1196 | L | 20 | Bifonazole | 2.2% | 3.3% | -11.7% | 4.3% | 60628-96-8 |
| Prestwick-P4 | 1197 | M | 20 | Imiquimod | 2.7% | -2.1% | 11.1% | 9.4% | 99011-02-6 |
| Prestwick-P4 | 1198 | N | 20 | Fosinopril | 2.9% | -1.7% | 6.7% | 6.5% | 98048-97-6 |
| Prestwick-P4 | 1199 | O | 20 | Acetylsalicylic acid | 3.2% | -4.1% | 5.1% | -8.8% | 50-78-2 |
| Prestwick-P4 | 1200 | P | 20 | Hexachlorophene | -9.8% | 14.3% | -23.4% | -11.5% | 70-30-4 |
| Prestwick-P4 | 1201 | A | 21 | Nelfinavir mesylate | -6.5% | -1.8% | -24.1% | 18.8% | 159989-65-8 |
| Prestwick-P4 | 1202 | B | 21 | Silodosin | -2.3% | -0.7% | 7.3% | 15.1% | 160970-54-7 |
| Prestwick-P4 | 1203 | C | 21 | Ipriflavone | 1.1% | -0.7% | -17.4% | 0.3% | 35212-22-7 |
| Prestwick-P4 | 1204 | D | 21 | Ezetimibe | -0.5% | 0.6% | -36.4% | 2.4% | 163222-33-1 |
| Prestwick-P4 | 1205 | E | 21 | (R)-Duloxetine hydrochloride | -0.6% | 4.7% | -23.9% | -1.5% | 116539-60-7 |
| Prestwick-P4 | 1206 | F | 21 | Donepezil hydrochloride | -1.3% | 6.9% | -5.0% | -0.2% | 120011-70-3 |
| Prestwick-P4 | 1207 | G | 21 | Aminacrine | -3.5% | 9.5% | -15.3% | -14.1% | 90-45-9 |
| Prestwick-P4 | 1208 | H | 21 | Pidotimod | 0.4% | 1.5% | -0.6% | 1.9% | 121808-62-6 |
| Prestwick-P4 | 1209 | I | 21 | Cefuroxime axetil | 0.6% | 0.7% | -8.4% | 1.7% | 64544-07-6 |
| Prestwick-P4 | 1210 | J | 21 | Anagrelide | 1.2% | 0.8% | 2.2% | 3.9% | 68475-42-3 |
| Prestwick-P4 | 1211 | K | 21 | Irbesartan | 3.1% | 4.1% | 8.6% | 7.1% | 138402-11-6 |
| Prestwick-P4 | 1212 | L | 21 | Indinavir sulfate | 3.2% | -0.6% | -4.2% | 3.4% | 157810-81-6 |
| Prestwick-P4 | 1213 | M | 21 | Epirubicin hydrochloride | -47.6% | -62.1% | -81.3% | -76.3% | 56390-09-1 |
| Prestwick-P4 | 1214 | N | 21 | Loteprednol etabonate | -1.0% | 3.8% | -15.3% | -10.1% | 82034-46-6 |
| Prestwick-P4 | 1215 | O | 21 | Cisatracurium besylate | -0.1% | -3.5% | -4.5% | -4.2% | 64228-79-1 |
| Prestwick-P4 | 1216 | P | 21 | Pemetrexed disodium | -3.8% | 5.1% | 3.4% | -2.8% | 357166-30-4 |
| Prestwick-P4 | co | A | 22 | DMSO | -4.5% | -1.3% | 8.1% | 22.0% | |
| Prestwick-P4 | co | B | 22 | DMSO | -0.4% | -2.3% | -1.5% | 15.2% | |
| Prestwick-P4 | co | C | 22 | Rott2 | -19.9% | 42.4% | -86.9% | -16.0% | |
| Prestwick-P4 | co | D | 22 | Rott2 | -21.3% | 44.8% | -85.5% | -24.0% | |
| Prestwick-P4 | co | E | 22 | Rott0.5 | -10.2% | 3.7% | -55.2% | -5.4% | |
| Prestwick-P4 | co | F | 22 | Rott0.5 | -8.3% | 9.8% | -53.7% | -11.2% | |
| Prestwick-P4 | co | G | 22 | Rott0.13 | -6.7% | 9.7% | -13.0% | -1.2% | |
| Prestwick-P4 | co | H | 22 | Rott0.13 | -4.8% | 3.5% | -15.9% | -3.5% | |
| Prestwick-P4 | co | I | 22 | DMSO | 0.2% | 6.2% | -7.1% | -2.6% | |
| Prestwick-P4 | co | J | 22 | DMSO | 1.9% | -1.5% | -7.6% | 6.4% | |
| Prestwick-P4 | co | K | 22 | DMSO | 1.1% | -0.8% | -6.4% | 3.1% | |
| Prestwick-P4 | co | L | 22 | DMSO | 3.7% | -1.8% | -4.9% | -0.4% | |
| Prestwick-P4 | co | M | 22 | DMSO | 1.6% | 1.8% | -2.3% | -2.6% | |
| Prestwick-P4 | co | N | 22 | DMSO | 1.9% | 0.2% | 26.2% | -12.7% | |
| Prestwick-P4 | co | O | 22 | DMSO | 0.0% | -1.1% | -5.6% | -8.6% | |
| Prestwick-P4 | co | P | 22 | DMSO | -2.1% | 2.5% | 8.3% | -1.4% | |
| Prestwick-P4 | co | A | 23 | DMSO | -2.7% | 2.1% | -5.5% | 25.4% | |
| Prestwick-P4 | co | B | 23 | DMSO | -1.0% | -4.4% | 1.8% | 22.8% | |

| | | | | | | | | | |
|---|---|---|---|---|---|---|---|---|---|
| Prestwick-P4 | co | C | 23 | DMSO | 1.8% | -1.9% | 1.8% | 1.3% | |
| Prestwick-P4 | co | D | 23 | DMSO | -1.1% | 0.9% | 7.1% | 14.9% | |
| Prestwick-P4 | co | E | 23 | DMSO | 2.2% | 6.2% | -1.8% | 6.6% | |
| Prestwick-P4 | co | F | 23 | DMSO | -4.1% | -2.0% | -3.9% | 20.1% | |
| Prestwick-P4 | co | G | 23 | DMSO | -1.9% | 0.1% | -11.2% | 2.5% | |
| Prestwick-P4 | co | H | 23 | DMSO | -1.1% | 3.5% | -8.2% | -11.1% | |
| Prestwick-P4 | co | I | 23 | DMSO | 1.0% | -0.9% | -8.4% | 0.9% | |
| Prestwick-P4 | co | J | 23 | DMSO | 1.0% | -2.7% | -12.9% | -1.0% | |
| Prestwick-P4 | co | K | 23 | DMSO | 0.7% | 0.6% | -3.0% | 6.3% | |
| Prestwick-P4 | co | L | 23 | DMSO | 0.8% | 0.3% | -7.0% | 17.6% | |
| Prestwick-P4 | co | M | 23 | DMSO | 0.8% | 1.8% | -7.1% | -6.5% | |
| Prestwick-P4 | co | N | 23 | DMSO | 0.3% | 1.8% | -5.1% | -2.7% | |
| Prestwick-P4 | co | O | 23 | DMSO | -3.8% | -0.1% | 1.0% | -8.0% | |
| Prestwick-P4 | co | P | 23 | DMSO | -1.2% | 1.5% | 6.4% | 4.6% | |
| Prestwick-P4 | co | A | 24 | noPFF | -2.7% | 4.1% | -100.0% | 9.3% | |
| Prestwick-P4 | co | B | 24 | noPFF | 1.6% | -4.1% | -100.0% | 4.7% | |
| Prestwick-P4 | co | C | 24 | noPFF | 1.1% | -0.5% | -99.9% | 11.8% | |
| Prestwick-P4 | co | D | 24 | Ceph4 | -25.4% | 61.9% | -99.9% | -44.9% | |
| Prestwick-P4 | co | E | 24 | noPFF | 0.4% | -5.8% | -100.0% | 19.9% | |
| Prestwick-P4 | co | F | 24 | noPFF | 0.3% | 1.6% | -99.9% | 16.1% | |
| Prestwick-P4 | co | G | 24 | noPFF | 2.1% | -0.6% | -100.0% | 7.8% | |
| Prestwick-P4 | co | H | 24 | noPFF | 1.1% | 0.8% | -100.0% | 3.6% | |
| Prestwick-P4 | co | I | 24 | noPFF | -1.4% | -0.1% | -100.0% | 9.9% | |
| Prestwick-P4 | co | J | 24 | noPFF | 0.6% | -0.1% | -100.0% | 14.2% | |
| Prestwick-P4 | co | K | 24 | noPFF | -0.1% | -1.6% | -100.0% | 12.6% | |
| Prestwick-P4 | co | L | 24 | noPFF | 2.0% | 0.6% | -99.9% | 6.4% | |
| Prestwick-P4 | co | M | 24 | noPFF | 1.1% | -2.3% | -100.0% | 0.8% | |
| Prestwick-P4 | co | N | 24 | noPFF | -2.0% | -0.4% | -99.9% | -2.5% | |
| Prestwick-P4 | co | O | 24 | noPFF | -1.3% | -0.4% | -99.9% | 10.5% | |
| Prestwick-P4 | co | P | 24 | noPFF | -4.3% | 2.0% | -100.0% | 9.9% | |
| Prestwick-P5 | co | A | 1 | DMSO | 1.7% | -5.2% | 0.6% | -15.3% | |
| Prestwick-P5 | co | B | 1 | DMSO | 1.8% | -1.5% | 9.9% | -7.7% | |
| Prestwick-P5 | co | C | 1 | DMSO | 0.4% | -5.6% | 9.4% | 0.4% | |
| Prestwick-P5 | co | D | 1 | DMSO | 1.3% | -5.1% | 12.9% | -3.1% | |
| Prestwick-P5 | co | E | 1 | DMSO | 0.4% | -0.3% | 8.5% | -7.7% | |
| Prestwick-P5 | co | F | 1 | DMSO | -1.1% | -6.4% | -3.3% | -8.2% | |
| Prestwick-P5 | co | G | 1 | DMSO | -0.6% | -2.2% | -5.8% | -5.0% | |
| Prestwick-P5 | co | H | 1 | DMSO | -0.2% | 2.6% | -9.8% | -5.4% | |
| Prestwick-P5 | co | I | 1 | DMSO | 0.0% | -2.7% | -10.1% | -2.6% | |
| Prestwick-P5 | co | J | 1 | DMSO | 2.7% | 0.0% | -21.0% | -4.0% | |
| Prestwick-P5 | co | K | 1 | DMSO | -0.3% | -3.8% | -23.4% | 6.2% | |
| Prestwick-P5 | co | L | 1 | DMSO | -0.7% | 2.4% | -3.1% | 2.9% | |
| Prestwick-P5 | co | M | 1 | DMSO | -1.4% | -0.7% | -5.5% | 5.4% | |
| Prestwick-P5 | co | N | 1 | DMSO | 3.1% | -1.6% | 2.8% | 0.7% | |
| Prestwick-P5 | co | O | 1 | DMSO | -1.4% | 6.0% | -3.7% | 6.5% | |
| Prestwick-P5 | co | P | 1 | DMSO | -3.1% | 17.9% | -20.8% | 8.5% | |
| Prestwick-P5 | co | A | 2 | DMSO | 0.1% | -6.1% | 18.8% | -1.6% | |
| Prestwick-P5 | co | B | 2 | DMSO | 3.0% | -1.0% | 22.2% | -6.9% | |
| Prestwick-P5 | co | C | 2 | DMSO | 0.1% | -2.4% | 10.3% | -9.8% | |
| Prestwick-P5 | co | D | 2 | DMSO | 1.0% | 3.6% | 10.1% | -9.3% | |
| Prestwick-P5 | co | E | 2 | DMSO | -3.4% | -4.6% | -7.3% | -4.1% | |
| Prestwick-P5 | co | F | 2 | DMSO | -1.3% | 1.3% | 2.8% | 0.0% | |
| Prestwick-P5 | co | G | 2 | DMSO | -0.3% | -2.2% | -11.6% | -5.1% | |
| Prestwick-P5 | co | H | 2 | DMSO | -0.2% | 1.6% | -5.8% | -6.8% | |
| Prestwick-P5 | co | I | 2 | Til2 | -8.0% | 16.5% | -84.1% | -3.8% | |
| Prestwick-P5 | co | J | 2 | Til2 | -5.6% | 19.0% | -81.2% | -4.3% | |
| Prestwick-P5 | co | K | 2 | Til0.5 | -1.1% | 0.4% | -41.8% | 5.4% | |
| Prestwick-P5 | co | L | 2 | Til0.5 | 0.9% | 6.7% | -36.1% | 2.8% | |
| Prestwick-P5 | co | M | 2 | Til0.13 | 1.4% | -2.7% | -18.0% | -2.7% | |
| Prestwick-P5 | co | N | 2 | Til0.13 | 0.2% | 3.5% | -3.8% | 0.6% | |
| Prestwick-P5 | co | O | 2 | DMSO | -4.6% | 3.4% | -4.8% | -0.2% | |
| Prestwick-P5 | co | P | 2 | DMSO | -4.9% | 11.8% | -12.4% | 13.5% | |
| Prestwick-P5 | 1217 | A | 3 | Trimebutine | -1.6% | 9.7% | 0.6% | -13.8% | 39133-31-8 |
| Prestwick-P5 | 1218 | B | 3 | Nevirapine | 1.8% | 2.0% | 7.8% | -11.1% | 129618-40-2 |
| Prestwick-P5 | 1219 | C | 3 | Rizatriptan benzoate | -0.2% | -1.1% | 3.6% | -2.2% | 145202-66-0 |
| Prestwick-P5 | 1220 | D | 3 | Tegaserod maleate | -31.0% | 77.7% | -96.8% | -43.6% | 189188-57-6 |
| Prestwick-P5 | 1221 | E | 3 | 1,8-Dihydroxyanthraquinone | -0.5% | 3.0% | 2.3% | -6.0% | 117-10-2 |
| Prestwick-P5 | 1222 | F | 3 | Nitazoxanide | 0.2% | 7.9% | -6.9% | -3.5% | 55981-09-4 |
| Prestwick-P5 | 1223 | G | 3 | Benidipine hydrochloride | -5.9% | 13.6% | -30.0% | -3.6% | 91599-74-5 |
| Prestwick-P5 | 1224 | H | 3 | Perospirone hydrochloride | 0.7% | 8.8% | -33.1% | -4.5% | 129273-38-7 |
| Prestwick-P5 | 1225 | I | 3 | Clopidogrel | 0.4% | 7.5% | -5.6% | -10.2% | 113665-84-2 |
| Prestwick-P5 | 1226 | J | 3 | Benzoxiquine | -30.6% | 82.2% | -99.4% | -24.5% | 86-75-9 |

| | | | | | | | | | |
|---|---|---|---|---|---|---|---|---|---|
| Prestwick-P5 | 1227 | K | 3 | Terbinafine | -2.2% | 10.1% | -8.3% | -40.9% | 91161-71-6 |
| Prestwick-P5 | 1228 | L | 3 | Histamine dihydrochloride | 4.4% | -2.0% | 5.1% | 2.3% | 56-92-8 |
| Prestwick-P5 | 1229 | M | 3 | Tolterodine tartrate | 9.1% | -4.0% | 35.5% | -0.7% | 209747-05-7 |
| Prestwick-P5 | 1230 | N | 3 | Lomerizine hydrochloride | -2.8% | 12.1% | -46.4% | -17.7% | 101477-54-7 |
| Prestwick-P5 | 1231 | O | 3 | Raltitrexed | 6.3% | 1.4% | 14.3% | -7.3% | 112887-68-0 |
| Prestwick-P5 | 1232 | P | 3 | Ceftibuten | 2.2% | 1.3% | 16.6% | -10.1% | 97519-39-6 |
| Prestwick-P5 | 1233 | A | 4 | Doxapram hydrochloride | 1.9% | -4.3% | 17.9% | -8.5% | 113-07-5 |
| Prestwick-P5 | 1234 | B | 4 | Amlexanox | 1.5% | -3.6% | 6.2% | -5.6% | 68302-57-8 |
| Prestwick-P5 | 1235 | C | 4 | Pantoprazole sodium | 1.6% | 0.1% | 7.9% | -8.3% | 138786-67-1 |
| Prestwick-P5 | 1236 | D | 4 | Tegafur | 0.7% | 1.0% | 7.3% | 2.8% | 17902-23-7 |
| Prestwick-P5 | 1237 | E | 4 | Nateglinide | 0.9% | -4.9% | -5.0% | -0.2% | 105816-04-4 |
| Prestwick-P5 | 1238 | F | 4 | Avobenzone | 1.0% | -5.0% | 10.0% | -3.0% | 70356-09-1 |
| Prestwick-P5 | 1239 | G | 4 | Cefpiramide | 0.2% | -3.2% | -9.4% | -8.3% | 70797-11-4 |
| Prestwick-P5 | 1240 | H | 4 | Fenoldopam | 1.3% | 1.0% | -11.5% | -0.2% | 67227-56-9 |
| Prestwick-P5 | 1241 | I | 4 | Phenothiazine | -2.1% | 4.6% | -27.4% | -6.9% | 92-84-2 |
| Prestwick-P5 | 1242 | J | 4 | Enalaprilat dihydrate | 2.7% | -1.6% | -3.8% | -8.8% | 84680-54-6 |
| Prestwick-P5 | 1243 | K | 4 | Rasagiline | 1.7% | -1.0% | -3.5% | -0.6% | 136236-51-6 |
| Prestwick-P5 | 1244 | L | 4 | Flumethasone pivalate | -1.6% | 5.0% | -2.9% | 3.9% | 2002-29-1 |
| Prestwick-P5 | 1245 | M | 4 | Ampiroxicam | 2.1% | -4.1% | -1.5% | -1.8% | 99464-64-9 |
| Prestwick-P5 | 1246 | N | 4 | Alosetron hydrochloride | 1.0% | 0.4% | 2.0% | 1.8% | 122852-69-1 |
| Prestwick-P5 | 1247 | O | 4 | Valsartan | 2.0% | -3.6% | -2.3% | -7.5% | 137862-53-4 |
| Prestwick-P5 | 1248 | P | 4 | Milnacipran hydrochloride | 0.8% | 0.4% | 1.9% | -4.2% | 101152-94-7 |
| Prestwick-P5 | 1249 | A | 5 | Amorolfine hydrochloride | -28.0% | 69.4% | -84.2% | -63.2% | 78613-38-4 |
| Prestwick-P5 | 1250 | B | 5 | Enrofloxacin | 1.5% | -1.9% | 14.4% | -1.9% | 93106-60-6 |
| Prestwick-P5 | 1251 | C | 5 | Tolcapone | 4.2% | -5.1% | -8.1% | -12.8% | 134308-13-7 |
| Prestwick-P5 | 1252 | D | 5 | Altrenogest | 3.1% | 0.6% | 2.7% | -7.9% | 850-52-2 |
| Prestwick-P5 | 1253 | E | 5 | Algestone acetophenide | -0.5% | 5.6% | -25.3% | -15.9% | 24356-94-3 |
| Prestwick-P5 | 1254 | F | 5 | Actarit | 1.2% | -0.9% | 1.5% | -4.5% | 18699-02-0 |
| Prestwick-P5 | 1255 | G | 5 | Adapalene | -2.6% | 8.8% | -23.0% | -14.3% | 106685-40-9 |
| Prestwick-P5 | 1256 | H | 5 | Diatrizoic acid dihydrate | 1.1% | 5.7% | 12.3% | -0.8% | 50978-11-5 |
| Prestwick-P5 | 1257 | I | 5 | Pregabalin | 0.7% | 0.3% | -6.9% | -3.1% | 148553-50-8 |
| Prestwick-P5 | 1258 | J | 5 | Dolutegravir | 0.8% | -1.9% | 3.7% | 3.8% | 1051375-16-6 |
| Prestwick-P5 | 1259 | K | 5 | Lofepramine | 0.5% | 5.8% | -18.3% | -5.9% | 23047-25-8 |
| Prestwick-P5 | 1260 | L | 5 | Valdecoxib | 4.9% | -3.9% | 0.5% | 3.4% | 181695-72-7 |
| Prestwick-P5 | 1261 | M | 5 | Risedronic acid monohydrate | 2.1% | -3.9% | -3.7% | -2.4% | 105462-24-6 |
| Prestwick-P5 | 1262 | N | 5 | Palonosetron hydrochloride | 6.1% | -0.2% | -7.7% | -4.1% | 135729-62-3 |
| Prestwick-P5 | 1263 | O | 5 | Triclabendazole | -4.6% | 10.1% | -19.6% | -13.7% | 68786-66-3 |
| Prestwick-P5 | 1264 | P | 5 | Brimonidine L-tartrate | -3.7% | 3.6% | -6.1% | 2.8% | 70359-46-5 |
| Prestwick-P5 | 1265 | A | 6 | Ubenimex | -0.7% | 0.1% | -12.6% | -10.7% | 58970-76-6 |
| Prestwick-P5 | 1266 | B | 6 | Troxipide | 2.2% | -0.6% | 11.4% | 4.2% | 99777-81-8 |
| Prestwick-P5 | 1267 | C | 6 | Felbamate | 2.5% | -4.5% | -8.6% | -9.9% | 25451-15-4 |
| Prestwick-P5 | 1268 | D | 6 | Estramustine | 2.0% | 8.5% | -11.5% | -5.8% | 2998-57-4 |
| Prestwick-P5 | 1269 | E | 6 | Ethoxzolamide | 2.7% | -2.4% | -3.5% | -8.5% | 452-35-7 |
| Prestwick-P5 | 1270 | F | 6 | Azatadine maleate | 3.9% | 1.7% | 8.6% | -3.6% | 3978-86-7 |
| Prestwick-P5 | 1271 | G | 6 | Dofetilide | 0.9% | 5.1% | -18.9% | -0.9% | 115256-11-6 |
| Prestwick-P5 | 1272 | H | 6 | Phenprobamate | 1.9% | 2.6% | -12.1% | -12.0% | 673-31-4 |
| Prestwick-P5 | 1273 | I | 6 | Zoledronic acid hydrate | 1.4% | -3.5% | -11.8% | -12.5% | 165800-06-6 |
| Prestwick-P5 | 1274 | J | 6 | Cefpodoxime proxetil | 2.1% | 0.8% | -4.1% | -9.9% | 87239-81-4 |
| Prestwick-P5 | 1275 | K | 6 | Besifloxacin hydrochloride | 3.2% | -0.3% | -10.3% | -14.5% | 141388-76-3 |
| Prestwick-P5 | 1276 | L | 6 | Ritonavir | 2.8% | -3.3% | -7.0% | 0.4% | 155213-67-5 |
| Prestwick-P5 | 1277 | M | 6 | Oxymetholone | -2.1% | 12.5% | -26.7% | -36.0% | 434-07-1 |
| Prestwick-P5 | 1278 | N | 6 | Latanoprost | 2.0% | 7.9% | 20.1% | -13.6% | 130209-82-4 |
| Prestwick-P5 | 1279 | O | 6 | Desonide | 3.7% | -6.3% | -10.4% | -17.8% | 638-94-8 |
| Prestwick-P5 | 1280 | P | 6 | Cefprozil | -2.2% | 1.4% | 1.5% | -3.5% | 121123-17-9 |
| Prestwick-P5 | 1281 | A | 7 | Allopurinol | -0.1% | -0.5% | 2.5% | -11.9% | 315-30-0 |
| Prestwick-P5 | 1282 | B | 7 | Lisuride (S)(-) | 3.4% | -1.9% | -1.9% | -2.6% | 18016-80-3 |
| Prestwick-P5 | 1283 | C | 7 | Carbenicillin disodium salt | 1.5% | -0.1% | -7.2% | -10.8% | 4800-94-6 |
| Prestwick-P5 | 1284 | D | 7 | Ebselen | -21.2% | 58.8% | -99.6% | -21.7% | 60940-34-3 |
| Prestwick-P5 | 1285 | E | 7 | Lapatinib | 2.7% | 3.2% | -16.4% | -18.4% | 231277-92-2 |
| Prestwick-P5 | 1286 | F | 7 | Clodronate disodium | 2.9% | -3.6% | 3.0% | -1.7% | 22560-50-5 |
| Prestwick-P5 | 1287 | G | 7 | Efaroxan hydrochloride | 1.9% | -3.2% | -1.4% | -8.8% | 89197-32-0 |
| Prestwick-P5 | 1288 | H | 7 | Cyromazine | 2.5% | 0.3% | 5.2% | 0.8% | 66215-27-8 |
| Prestwick-P5 | 1289 | I | 7 | Exalamide | 0.5% | -3.0% | -16.7% | -3.3% | 53370-90-4 |
| Prestwick-P5 | 1290 | J | 7 | Anisindione | 4.0% | -6.2% | 1.3% | -8.2% | 117-37-3 |
| Prestwick-P5 | 1291 | K | 7 | Cyclandelate | -4.8% | 23.8% | -29.4% | -14.7% | 456-59-7 |
| Prestwick-P5 | 1292 | L | 7 | Prasterone | 5.8% | -0.5% | -1.0% | 3.3% | 53-43-0 |
| Prestwick-P5 | 1293 | M | 7 | Erdosteine | 5.0% | -1.5% | 3.1% | -1.6% | 84611-23-4 |
| Prestwick-P5 | 1294 | N | 7 | loxoprofen | 5.1% | -4.1% | 2.5% | -2.6% | 68767-14-6 |
| Prestwick-P5 | 1295 | O | 7 | Piperonyl butoxide | -0.4% | 11.1% | -27.8% | -13.0% | 51-03-6 |
| Prestwick-P5 | 1296 | P | 7 | Eprobemide | 0.2% | 4.5% | 1.4% | -5.9% | 87940-60-1 |
| Prestwick-P5 | 1297 | A | 8 | Ampicillin sodium salt | -0.4% | -1.1% | 3.4% | 2.4% | 69-52-3 |
| Prestwick-P5 | 1298 | B | 8 | Flupentixol dihydrochloride cis-(Z) | -5.9% | 22.7% | -47.6% | -16.4% | 2413-38-9 |

| | | | | | | | | | |
|---|---|---|---|---|---|---|---|---|---|
| Prestwick-P5 | 1299 | C | 8 | Cefamandole sodium salt | 1.1% | 1.0% | -16.4% | -13.7% | 30034-03-8 |
| Prestwick-P5 | 1300 | D | 8 | Rolipram | 3.1% | 1.0% | -7.4% | 1.1% | 61413-54-5 |
| Prestwick-P5 | 1301 | E | 8 | Mevastatin | -1.6% | 8.5% | -19.9% | -13.9% | 73573-88-3 |
| Prestwick-P5 | 1302 | F | 8 | Entecavir | 6.1% | 1.2% | 13.6% | -7.1% | 142217-69-4 |
| Prestwick-P5 | 1303 | G | 8 | Oxiglutatione | 3.1% | -3.3% | -5.9% | -5.7% | 27025-41-8 |
| Prestwick-P5 | 1304 | H | 8 | Bucetin | 3.6% | 0.5% | 5.5% | -7.0% | 1083-57-4 |
| Prestwick-P5 | 1305 | I | 8 | Ancitabine hydrochloride | 2.1% | -4.5% | -2.3% | -16.0% | 10212-25-6 |
| Prestwick-P5 | 1306 | J | 8 | Acetaminosalol | 5.5% | -4.8% | -8.3% | -5.4% | 118-57-0 |
| Prestwick-P5 | 1307 | K | 8 | Beclamide | 2.9% | 0.5% | -8.0% | 4.8% | 501-68-8 |
| Prestwick-P5 | 1308 | L | 8 | Deferiprone | 6.3% | -1.2% | 1.0% | -1.6% | 30652-11-0 |
| Prestwick-P5 | 1309 | M | 8 | Acedoben | 4.1% | -2.8% | 1.2% | -2.4% | 556-08-1 |
| Prestwick-P5 | 1310 | N | 8 | Dimetridazole | 2.9% | -5.3% | -6.1% | -0.8% | 551-92-8 |
| Prestwick-P5 | 1311 | O | 8 | Tioguanine | 6.2% | -6.5% | -5.1% | -15.9% | 154-42-7 |
| Prestwick-P5 | 1312 | P | 8 | Trehalose dihydrate | 0.9% | 1.4% | -4.6% | 1.2% | 6138-23-4 |
| Prestwick-P5 | 1313 | A | 9 | Cephalexin monohydrate | 0.1% | -2.6% | 4.5% | 1.4% | 23325-78-2 |
| Prestwick-P5 | 1314 | B | 9 | Cholecalciferol | 1.8% | 3.0% | 6.8% | 6.4% | 67-97-0 |
| Prestwick-P5 | 1315 | C | 9 | Tiletamine hydrochloride | 0.3% | -1.6% | -3.7% | -0.2% | 14176-50-2 |
| Prestwick-P5 | 1316 | D | 9 | Netilmicin sulfate | 3.2% | 3.9% | 3.7% | -6.7% | 56391-57-2 |
| Prestwick-P5 | 1317 | E | 9 | Cyclophosphamide | 2.2% | -1.2% | 3.7% | -2.6% | 50-18-0 |
| Prestwick-P5 | 1318 | F | 9 | Quinine ethyl carbonate | 1.5% | 0.0% | -3.6% | -2.7% | 83-75-0 |
| Prestwick-P5 | 1319 | G | 9 | Oxeladin citrate | -0.5% | 4.7% | -4.8% | -3.4% | 52432-72-1, 468-[illegible] |
| Prestwick-P5 | 1320 | H | 9 | Dichlorisone acetate | -4.7% | 20.3% | -21.1% | -10.7% | 79-61-8 |
| Prestwick-P5 | 1321 | I | 9 | Tolperisone hydrochloride | -0.7% | 0.4% | -10.1% | -0.1% | 70312-00-4 |
| Prestwick-P5 | 1322 | J | 9 | Broxyquinoline | -30.0% | 76.0% | -68.2% | -37.9% | 521-74-4 |
| Prestwick-P5 | 1323 | K | 9 | Orotic acid | 2.9% | -1.3% | 7.7% | 4.5% | 65-86-1 |
| Prestwick-P5 | 1324 | L | 9 | Isaxonine | 3.0% | -4.7% | -6.5% | 6.0% | 4214-72-6 |
| Prestwick-P5 | 1325 | M | 9 | Nikethamide | 2.4% | -1.9% | 29.4% | -1.9% | 59-26-7 |
| Prestwick-P5 | 1326 | N | 9 | Paroxypropione | 2.9% | -0.8% | 4.9% | -8.9% | 70-70-2 |
| Prestwick-P5 | 1327 | O | 9 | Amezinium metilsulfate | 2.3% | 2.3% | 1.9% | -1.8% | 30578-37-1 |
| Prestwick-P5 | 1328 | P | 9 | Allylestrenol | -32.2% | 81.3% | -83.7% | -39.7% | 432-60-0 |
| Prestwick-P5 | 1329 | A | 10 | Methylene blue | -0.3% | -1.4% | -44.3% | 5.7% | 61-73-4 |
| Prestwick-P5 | 1330 | B | 10 | Gramicidin | -39.5% | 100.5% | -95.4% | 2.7% | 1405-97-6 |
| Prestwick-P5 | 1331 | C | 10 | Loracarbef | 0.4% | -2.5% | -0.8% | 6.7% | 76470-66-1 |
| Prestwick-P5 | 1332 | D | 10 | Cyclacillin | 2.1% | 0.7% | -4.3% | 0.0% | 3485-14-1 |
| Prestwick-P5 | 1333 | E | 10 | Edaravone | 2.0% | -1.6% | -3.0% | 2.0% | 89-25-8 |
| Prestwick-P5 | 1334 | F | 10 | Timonacic | 0.2% | 2.7% | 3.2% | -4.8% | 444-27-9 |
| Prestwick-P5 | 1335 | G | 10 | D-phenylalanine | 1.0% | -2.0% | -6.7% | 2.0% | 673-06-3 |
| Prestwick-P5 | 1336 | H | 10 | Flopropione | 2.9% | -1.5% | 1.5% | -6.6% | 2295-58-1 |
| Prestwick-P5 | 1337 | I | 10 | Ethacridine lactate | -5.2% | 2.1% | -61.3% | -12.6% | 1837-57-6; 6402-[illegible] |
| Prestwick-P5 | 1338 | J | 10 | Clofoctol | -35.8% | 90.8% | -96.8% | -59.5% | 37693-01-9 |
| Prestwick-P5 | 1339 | K | 10 | Proflavine hemisulfate | -15.2% | -2.7% | -59.7% | -9.3% | 553-30-0, 1811-2[illegible] |
| Prestwick-P5 | 1340 | L | 10 | Climbazole | 2.3% | 2.1% | -1.7% | -1.4% | 38083-17-9 |
| Prestwick-P5 | 1341 | M | 10 | Nitroxoline | 4.3% | -4.3% | 5.6% | -5.6% | 4008-48-4 |
| Prestwick-P5 | 1342 | N | 10 | Mexenone | -3.6% | 13.5% | -15.4% | -18.3% | 1641-17-4 |
| Prestwick-P5 | 1343 | O | 10 | Carazolol | 2.1% | -3.4% | 17.8% | -6.2% | 57775-29-8 |
| Prestwick-P5 | 1344 | P | 10 | Benactyzine hydrochloride | -6.7% | 7.9% | -26.9% | -24.1% | 57-37-4 |
| Prestwick-P5 | 1345 | A | 11 | Levocarnitine | 2.3% | -4.0% | 2.8% | -5.1% | 541-15-1 |
| Prestwick-P5 | 1346 | B | 11 | DO 897/99 | 1.8% | -0.2% | -17.6% | 7.3% | 314776-92-6 |
| Prestwick-P5 | 1347 | C | 11 | Thimerosal | -16.8% | 49.1% | -99.5% | -55.2% | 54-64-8 |
| Prestwick-P5 | 1348 | D | 11 | Clindamycin Phosphate | 1.1% | 5.9% | -2.1% | -9.4% | 24729-96-2 |
| Prestwick-P5 | 1349 | E | 11 | Tioxolone | 0.8% | -1.6% | -11.7% | -7.7% | 4991-65-5 |
| Prestwick-P5 | 1350 | F | 11 | Metacetamol | 1.9% | 0.9% | 0.1% | -6.1% | 621-42-1 |
| Prestwick-P5 | 1351 | G | 11 | Chromocarb | 1.6% | 1.3% | -1.8% | -11.7% | 4940-39-0 |
| Prestwick-P5 | 1352 | H | 11 | Betamipron | 0.9% | 2.5% | -7.1% | -7.5% | 3440-28-6 |
| Prestwick-P5 | 1353 | I | 11 | Pantethine | 3.4% | -2.4% | -15.9% | -3.7% | 16816-67-4 |
| Prestwick-P5 | 1354 | J | 11 | Cepharanthine | 1.7% | 4.8% | -91.9% | -17.1% | 481-49-2 |
| Prestwick-P5 | 1355 | K | 11 | Moguisteine | 2.2% | 0.2% | 2.6% | 4.0% | 119637-67-1 |
| Prestwick-P5 | 1356 | L | 11 | Nefiracetam | 1.8% | 1.9% | 1.2% | 6.5% | 77191-36-7 |
| Prestwick-P5 | 1357 | M | 11 | Sulfacarbamide | 3.6% | -3.6% | -5.6% | -0.9% | 547-44-4 |
| Prestwick-P5 | 1358 | N | 11 | Ethenzamide | 3.5% | 2.4% | 7.7% | -4.3% | 938-73-8 |
| Prestwick-P5 | 1359 | O | 11 | Brivudine | 3.9% | -2.6% | 0.7% | -10.3% | 69304-47-8 |
| Prestwick-P5 | 1360 | P | 11 | Sulfisomidine | -0.9% | 6.0% | 3.8% | -10.9% | 515-64-0 |
| Prestwick-P5 | 1361 | A | 12 | Epalrestat | -1.0% | 0.2% | -7.1% | 7.4% | 82159-09-9 |
| Prestwick-P5 | 1362 | B | 12 | Cefcapene pivoxil hydrochloride | -0.6% | 4.3% | -6.7% | -8.0% | 147816-23-7 |
| Prestwick-P5 | 1363 | C | 12 | Lobenzarit disodium | 1.3% | -3.0% | -3.1% | -6.8% | 64808-48-6 |
| Prestwick-P5 | 1364 | D | 12 | Dalfampridine | 1.4% | 3.2% | 7.0% | -6.9% | 504-24-5 |
| Prestwick-P5 | 1365 | E | 12 | Bendazac | 2.7% | 0.3% | -11.5% | -7.5% | 20187-55-7 |
| Prestwick-P5 | 1366 | F | 12 | Bithionol | -3.9% | 9.9% | -23.4% | -24.4% | 97-18-7 |
| Prestwick-P5 | 1367 | G | 12 | Ecabet sodium | -0.3% | -0.6% | 1.5% | -10.8% | 219773-47-4 |
| Prestwick-P5 | 1368 | H | 12 | Prifinium Bromide | 0.6% | 7.0% | -14.0% | -20.1% | 4630-95-9 |
| Prestwick-P5 | 1369 | I | 12 | Dichlorophen | -5.8% | 13.0% | -41.3% | -1.0% | 97-23-4 |
| Prestwick-P5 | 1370 | J | 12 | Glutathione | 0.7% | -0.2% | -5.0% | -2.5% | 70-18-8 |

| | | | | | | | | | |
|---|---|---|---|---|---|---|---|---|---|
| Prestwick-P5 | 1371 | K | 12 | Abemaciclib | 1.5% | 16.6% | -98.8% | -13.8% | 1231929-97-7 |
| Prestwick-P5 | 1372 | L | 12 | Clopidol | 1.8% | -1.5% | -4.3% | -0.8% | 2971-90-6 |
| Prestwick-P5 | 1373 | M | 12 | Mizoribine | -1.4% | -0.3% | -5.4% | 0.6% | 50924-49-7 |
| Prestwick-P5 | 1374 | N | 12 | Imidapril hydrochloride | 2.6% | 1.9% | 2.8% | -0.7% | 89396-94-1 |
| Prestwick-P5 | 1375 | O | 12 | Fipronil | -1.1% | -1.0% | -14.1% | -4.8% | 120068-37-3 |
| Prestwick-P5 | 1376 | P | 12 | Cefpirome | -2.0% | 2.1% | -4.9% | -11.4% | 84957-29-9 |
| Prestwick-P5 | 1377 | A | 13 | Ceftezole | 2.2% | -0.9% | 5.4% | 1.8% | 26973-24-0 |
| Prestwick-P5 | 1378 | B | 13 | Carbazochrome | 0.6% | -1.4% | 7.7% | 2.8% | 69-81-8 |
| Prestwick-P5 | 1379 | C | 13 | L-tryptophan | 2.2% | 2.9% | -14.7% | -11.4% | 73-22-3 |
| Prestwick-P5 | 1380 | D | 13 | Fenclozic acid | 1.5% | 0.8% | -4.6% | -6.8% | 17969-20-9 |
| Prestwick-P5 | 1381 | E | 13 | Methandrostenolone | 2.3% | -1.7% | -14.2% | -3.4% | 72-63-9 |
| Prestwick-P5 | 1382 | F | 13 | (R)-Tamsulosin hydrochloride | 1.1% | 5.8% | 19.7% | 0.1% | 106463-17-6 |
| Prestwick-P5 | 1383 | G | 13 | Trandolapril | 0.5% | 2.3% | 1.7% | -8.7% | 87679-37-6 |
| Prestwick-P5 | 1384 | H | 13 | Benziodarone | -0.4% | 3.4% | -18.2% | -7.5% | 68-90-6 |
| Prestwick-P5 | 1385 | I | 13 | Valethamate bromide | 2.1% | 1.4% | -18.9% | -1.2% | 90-22-2 |
| Prestwick-P5 | 1386 | J | 13 | Ebastine | -28.3% | 73.4% | -96.4% | -63.5% | 90729-43-4 |
| Prestwick-P5 | 1387 | K | 13 | Phthalylsulfacetamide | 2.1% | -1.4% | -3.5% | 3.9% | 131-69-1 |
| Prestwick-P5 | 1388 | L | 13 | Cresopirine | -0.5% | 1.0% | 4.7% | 9.6% | 4386-39-4 |
| Prestwick-P5 | 1389 | M | 13 | Cefsulodin Sodium Salt Hydrate | 1.5% | 0.6% | -10.4% | -12.1% | 1426397-23-0 |
| Prestwick-P5 | 1390 | N | 13 | Malotilate | -3.8% | 11.1% | -22.9% | -13.8% | 59937-28-9 |
| Prestwick-P5 | 1391 | O | 13 | �fenamate | -4.4% | 13.0% | -7.0% | -33.4% | 67330-25-0 |
| Prestwick-P5 | 1392 | P | 13 | Benzarone | -4.9% | 13.6% | -26.7% | -14.7% | 1477-19-6 |
| Prestwick-P5 | 1393 | A | 13 | Ivabradine hydrochloride | -0.1% | 1.3% | -10.3% | -6.6% | 148849-67-6 |
| Prestwick-P5 | 1394 | B | 14 | Oxiracetam | -0.3% | -2.0% | 2.4% | -3.1% | 62613-82-5 |
| Prestwick-P5 | 1395 | C | 14 | Exifone | -0.5% | -0.1% | 12.2% | -14.1% | 52479-85-3 |
| Prestwick-P5 | 1396 | D | 14 | Phenolphthalein | 0.6% | 1.0% | 0.6% | -7.0% | 77-09-8 |
| Prestwick-P5 | 1397 | E | 14 | Fasudil Hydrochloride | 1.7% | 2.1% | -13.2% | -2.0% | 103745-39-7 |
| Prestwick-P5 | 1398 | F | 14 | Acetyl-L-glutamine | 1.1% | 1.5% | -7.3% | 3.0% | 2490-97-3 |
| Prestwick-P5 | 1399 | G | 14 | Azaribine | 3.0% | -1.7% | 11.0% | -8.0% | 2169-64-4 |
| Prestwick-P5 | 1400 | H | 14 | Mebanazine | 3.3% | 3.7% | 8.7% | -5.4% | 65-64-5 |
| Prestwick-P5 | 1401 | I | 14 | Pirfenidone | 0.2% | -3.9% | -17.2% | -9.6% | 53179-13-8 |
| Prestwick-P5 | 1402 | J | 14 | Chlorquinaldol | -4.7% | 27.6% | -28.5% | -28.6% | 72-80-0 |
| Prestwick-P5 | 1403 | K | 14 | Rufinamide | 2.3% | -1.7% | -8.2% | 6.2% | 106308-44-5 |
| Prestwick-P5 | 1404 | L | 14 | Tinoridine hydrochloride | 0.8% | 2.6% | -10.0% | -11.5% | 25913-34-2 |
| Prestwick-P5 | 1405 | M | 14 | Artesunate | 2.2% | 0.0% | -5.4% | 2.8% | 88495-63-0 |
| Prestwick-P5 | 1406 | N | 14 | Gestodene | 1.5% | -1.2% | -10.6% | -4.0% | 60282-87-3 |
| Prestwick-P5 | 1407 | O | 14 | Bergenin | 0.9% | -0.7% | -5.7% | -11.6% | 477-90-7 |
| Prestwick-P5 | 1408 | P | 14 | Nepafenac | -1.7% | 1.7% | -7.8% | -4.0% | 78281-72-8 |
| Prestwick-P5 | 1409 | A | 15 | Zaltoprofen | 2.7% | -2.9% | 6.1% | 0.8% | 74711-43-6 |
| Prestwick-P5 | 1410 | B | 15 | Cianidanol | 2.5% | -4.5% | -0.3% | 0.2% | 154-23-4 |
| Prestwick-P5 | 1411 | C | 15 | Quatacaine | 3.0% | -0.1% | 3.6% | -2.1% | 17692-45-4 |
| Prestwick-P5 | 1412 | D | 15 | Eperisone HCl | -0.9% | 8.0% | -11.4% | -2.5% | 56839-43-1 |
| Prestwick-P5 | 1413 | E | 15 | Tulobuterol Hydrochloride | 1.1% | -1.6% | -6.2% | 4.4% | 56776-01-3 |
| Prestwick-P5 | 1414 | F | 15 | Delapril Hydrochloride | 0.1% | 1.7% | -2.2% | -4.6% | 83435-67-0 |
| Prestwick-P5 | 1415 | G | 15 | Acetanilide | 0.9% | 2.2% | -8.1% | 10.6% | 103-84-4 |
| Prestwick-P5 | 1416 | H | 15 | Fudosteine | -1.0% | 1.3% | -4.0% | 5.2% | 13189-98-5 |
| Prestwick-P5 | 1417 | I | 15 | Cypermethrin | -0.3% | 3.4% | -17.8% | -8.9% | 52315-07-8 |
| Prestwick-P5 | 1418 | J | 15 | Phenothrin | 0.8% | 8.3% | 1.3% | -11.9% | 26002-80-2 |
| Prestwick-P5 | 1419 | K | 15 | Chlormidazole | -1.9% | 7.6% | -6.6% | 2.9% | 3689-76-7 |
| Prestwick-P5 | 1420 | L | 15 | Indometacin farnesil | 2.7% | 6.2% | -19.1% | 2.1% | 85801-02-1 |
| Prestwick-P5 | 1421 | M | 15 | Stiripentol | 2.3% | 3.3% | -4.0% | -9.9% | 49763-96-4 |
| Prestwick-P5 | 1422 | N | 15 | Safinamide | 1.0% | 5.7% | -6.8% | -1.9% | 133865-89-1 |
| Prestwick-P5 | 1423 | O | 15 | Voglibose | 1.7% | -1.4% | 2.1% | -14.7% | 83480-29-9 |
| Prestwick-P5 | 1424 | P | 15 | Arotinolol hydrochloride | -1.5% | 5.0% | -6.6% | -5.5% | 68377-91-3 |
| Prestwick-P5 | 1425 | A | 16 | Sulbenicillin sodium salt | -1.2% | -0.6% | 0.7% | -0.3% | 28002-18-8 |
| Prestwick-P5 | 1426 | B | 16 | Carbazochrome sodium sulfonate | 0.0% | -1.9% | 2.1% | 13.0% | 51460-26-5 |
| Prestwick-P5 | 1427 | C | 16 | Prosultiamine | 1.2% | 3.6% | -13.8% | -5.1% | 59-58-5 |
| Prestwick-P5 | 1428 | D | 16 | Tiaramide Hydrochloride | 1.9% | 0.2% | -0.8% | -8.6% | 35941-71-0 |
| Prestwick-P5 | 1429 | E | 16 | Suplatast Tosylate | -1.0% | 3.9% | -7.1% | 7.1% | 94055-76-2 |
| Prestwick-P5 | 1430 | F | 16 | Manidipine Dihydrochloride | -4.9% | 15.8% | -75.6% | -12.1% | 89226-75-5 |
| Prestwick-P5 | 1431 | G | 16 | Mosapride | 5.1% | 0.4% | -22.5% | -13.5% | 112885-41-3 |
| Prestwick-P5 | 1432 | H | 16 | Inosine | 1.9% | 0.8% | 5.1% | -4.3% | 58-63-9 |
| Prestwick-P5 | 1433 | I | 16 | Chloramine-T hydrate | 0.5% | -0.4% | -8.3% | -5.1% | 127-65-1 |
| Prestwick-P5 | 1434 | J | 16 | Tolonium chloride | -7.0% | 16.4% | -94.9% | -25.1% | 92-31-9 |
| Prestwick-P5 | 1435 | K | 16 | Bosutinib | -0.3% | 9.5% | -95.3% | -7.7% | 380843-75-4 |
| Prestwick-P5 | 1436 | L | 16 | Tiagabine hydrochloride | 0.8% | 0.8% | 4.4% | 5.1% | 115103-54-3 |
| Prestwick-P5 | 1437 | M | 16 | Cephradine monohydrate | 1.7% | 3.0% | -3.5% | 1.4% | 75975-70-1 |
| Prestwick-P5 | 1438 | N | 16 | Doripenem monohydrate | 2.1% | -1.7% | 4.3% | -8.4% | 364622-82-2 |
| Prestwick-P5 | 1439 | O | 16 | Clevidipine | -1.7% | 12.7% | -16.1% | -0.5% | 167221-71-8 |
| Prestwick-P5 | 1440 | P | 16 | Artenimol | -0.8% | 0.6% | -14.3% | -9.9% | 81496-81-3 |
| Prestwick-P5 | 1441 | A | 17 | Befunolol | -1.0% | 0.9% | -10.3% | 3.5% | 39552-01-7 |
| Prestwick-P5 | 1442 | B | 17 | Eletriptan | 4.7% | -4.7% | 12.9% | 4.3% | 143322-58-1 |

| | | | | | | | | | |
|---|---|---|---|---|---|---|---|---|---|
| Prestwick-P5 | 1443 | C | 17 | Budralazine | 2.0% | 0.9% | -12.6% | -13.1% | 36798-79-5 |
| Prestwick-P5 | 1444 | D | 17 | Bunazosin hydrochloride | 0.8% | 2.8% | -8.5% | 1.3% | 52712-76-2 |
| Prestwick-P5 | 1445 | E | 17 | Kanamycin B | 0.3% | 3.2% | -7.6% | 9.1% | 4696-76-8 |
| Prestwick-P5 | 1446 | F | 17 | Mitomycin C | -5.8% | 20.4% | -41.9% | -37.6% | 50-07-7 |
| Prestwick-P5 | 1447 | G | 17 | Prasugrel | 1.3% | 2.4% | 4.3% | -2.5% | 150322-43-3 |
| Prestwick-P5 | 1448 | H | 17 | Ramosetron hydrochloride | 2.7% | -1.8% | -9.7% | -0.1% | 132907-72-3 |
| Prestwick-P5 | 1449 | I | 17 | Temocapril | 1.7% | -0.1% | -13.9% | -3.3% | 111902-57-9 |
| Prestwick-P5 | 1450 | J | 17 | Flosequinan | 2.6% | -1.1% | 2.3% | -2.9% | 76568-02-0 |
| Prestwick-P5 | 1451 | K | 17 | Ebrotidine | 3.0% | 5.4% | -18.8% | 3.8% | 100981-43-9 |
| Prestwick-P5 | 1452 | L | 17 | Nedocromil sodium | 6.7% | -0.8% | -5.6% | 1.7% | 69049-73-6 |
| Prestwick-P5 | 1453 | M | 17 | Cinacalcet hydrochloride | -23.8% | 67.7% | -81.1% | -23.1% | 226256-56-0 |
| Prestwick-P5 | 1454 | N | 17 | Ibufenac | 5.4% | -0.2% | 6.3% | -7.5% | 1553-60-2 |
| Prestwick-P5 | 1455 | O | 17 | Vandetanib | 4.8% | 0.4% | -82.8% | -26.7% | 443913-73-3 |
| Prestwick-P5 | 1456 | P | 17 | Bromfenac | -1.6% | 3.7% | -11.7% | -11.2% | 91714-94-2 |
| Prestwick-P5 | 1457 | A | 18 | Pirprofen | -0.6% | 0.0% | 6.2% | 0.0% | 31793-07-4 |
| Prestwick-P5 | 1458 | B | 18 | Tripamide | 1.3% | 3.3% | 15.5% | 3.0% | 73803-48-2 |
| Prestwick-P5 | 1459 | C | 18 | Chlorpromazine Sulfoxide | 1.9% | -2.9% | 9.2% | -2.9% | 969-99-3 |
| Prestwick-P5 | 1460 | D | 18 | Cycotiamine | 1.0% | -0.1% | 1.7% | -5.3% | 6092-18-8 |
| Prestwick-P5 | 1461 | E | 18 | Nafamostat Mesylate | 1.9% | 2.7% | -7.0% | -4.6% | 82956-11-4 |
| Prestwick-P5 | 1462 | F | 18 | Bisbentiamine | 2.4% | 4.9% | -14.3% | -2.1% | 2667-89-2 |
| Prestwick-P5 | 1463 | G | 18 | Seratrodast | 2.6% | 5.1% | 5.5% | 0.2% | 112665-43-7 |
| Prestwick-P5 | 1464 | H | 18 | Sofalcone | -3.5% | 11.6% | 1.1% | -14.7% | 64506-49-6 |
| Prestwick-P5 | 1465 | I | 18 | Etravirine | -8.3% | 25.7% | -38.6% | -12.1% | 269055-15-4 |
| Prestwick-P5 | 1466 | J | 18 | Empagliflozin | 2.9% | 0.2% | 6.3% | -2.5% | 864070-44-0 |
| Prestwick-P5 | 1467 | K | 18 | Frovatriptan succinate | 2.7% | -1.0% | 1.4% | -3.0% | 158930-17-7 |
| Prestwick-P5 | 1468 | L | 18 | Vonoprazan | 2.5% | 1.4% | -7.7% | -6.0% | 881681-00-1 |
| Prestwick-P5 | 1469 | M | 18 | Mofezolac | -0.9% | 0.7% | -15.0% | 3.2% | 78967-07-4 |
| Prestwick-P5 | 1470 | N | 18 | Hopantenate calcium | 3.9% | -5.7% | 0.8% | -8.2% | 17097-76-6 |
| Prestwick-P5 | 1471 | O | 18 | Acetyl spiramycin | -0.2% | 8.0% | -77.7% | 2.0% | 24916-51-6 |
| Prestwick-P5 | 1472 | P | 18 | Talampicillin hydrochloride | -1.5% | 6.0% | -16.7% | -7.5% | 39878-70-1 |
| Prestwick-P5 | 1473 | A | 19 | Zanamivir | -0.4% | 0.3% | 12.5% | 17.1% | 139110-80-8 |
| Prestwick-P5 | 1474 | B | 19 | Alclofenac | 0.8% | 2.3% | 0.4% | 7.8% | 22131-79-9 |
| Prestwick-P5 | 1475 | C | 19 | Fenclofenac | 2.0% | -2.1% | 0.9% | 2.0% | 34645-84-6 |
| Prestwick-P5 | 1476 | D | 19 | Flurbiprofen axetil | 0.4% | 2.0% | -1.5% | 0.3% | 91503-79-6 |
| Prestwick-P5 | 1477 | E | 19 | Camostat mesylate | -0.8% | 0.5% | 3.9% | 2.7% | 59721-29-8 |
| Prestwick-P5 | 1478 | F | 19 | Ticagrelor | -6.7% | 25.6% | -13.0% | -34.2% | 274693-27-5 |
| Prestwick-P5 | 1479 | G | 19 | Sarpogrelate hydrochloride | 1.9% | 4.7% | -4.8% | 3.5% | 135159-51-2 |
| Prestwick-P5 | 1480 | H | 19 | Sapropterin dihydrochloride | 1.3% | -0.3% | 1.1% | -5.9% | 69056-38-8 |
| Prestwick-P5 | 1481 | I | 19 | Crisaborole | 1.3% | 2.0% | 0.2% | -3.4% | 906673-24-3 |
| Prestwick-P5 | 1482 | J | 19 | Avanafil | 0.7% | -1.5% | -12.7% | 4.8% | 330784-47-9 |
| Prestwick-P5 | 1483 | K | 19 | Azasetron hydrochloride | 2.0% | 1.4% | -8.2% | 7.9% | 123040-16-4 |
| Prestwick-P5 | 1484 | L | 19 | Buformin | 2.3% | -1.6% | 12.7% | 5.0% | 692-13-7 |
| Prestwick-P5 | 1485 | M | 19 | Pyricarbate | 12.2% | 0.6% | -16.3% | -11.3% | 1882-26-4 |
| Prestwick-P5 | 1486 | N | 19 | Phenoxypropazine dihydrochloride | 3.4% | -3.8% | 4.8% | -1.6% | 3818-37-9 |
| Prestwick-P5 | 1487 | O | 19 | Cinepazide maleate | 1.8% | -2.3% | 9.6% | -7.2% | 26328-04-1 |
| Prestwick-P5 | 1488 | P | 19 | Procaterol hydrochloride | -1.4% | 3.3% | -0.7% | -13.8% | 59828-07-8 |
| Prestwick-P5 | 1489 | A | 20 | Angiotensin II (Human) Trifluoroacetic acid salt | -0.8% | -1.5% | 11.5% | 19.1% | 4474-91-3 |
| Prestwick-P5 | 1490 | B | 20 | Anisotropine Methylbromide | -2.4% | 4.3% | -2.7% | 11.7% | 80-50-2 |
| Prestwick-P5 | 1491 | C | 20 | Ibuprofen Piconol | -11.7% | 40.5% | -38.9% | -30.1% | 64622-45-3 |
| Prestwick-P5 | 1492 | D | 20 | Pilsicainide Hydrochloride | -1.6% | -0.1% | -10.6% | 2.3% | 88069-49-2 |
| Prestwick-P5 | 1493 | E | 20 | Feprazone | -0.6% | 5.9% | -4.2% | 2.0% | 30748-29-9 |
| Prestwick-P5 | 1494 | F | 20 | Lanoconazole | -1.8% | 9.1% | -15.9% | 2.6% | 101530-10-3 |
| Prestwick-P5 | 1495 | G | 20 | Mibefradil | -22.0% | 49.6% | -99.0% | -50.2% | 116644-53-2 |
| Prestwick-P5 | 1496 | H | 20 | Midecamycin | -1.7% | -0.3% | -11.7% | 9.0% | 35457-80-8 |
| Prestwick-P5 | 1497 | I | 20 | Salinomycin sodium | -21.4% | 40.5% | -98.8% | -49.0% | 55721-31-8 |
| Prestwick-P5 | 1498 | J | 20 | Enocitabine | 7.1% | 0.2% | -13.4% | -3.9% | 55726-47-1 |
| Prestwick-P5 | 1499 | K | 20 | Propanidid | -0.2% | 2.3% | -6.4% | 10.3% | 1421-14-3 |
| Prestwick-P5 | 1500 | L | 20 | Lomitapide | -23.5% | 62.0% | -98.5% | -67.5% | 182431-12-5 |
| Prestwick-P5 | 1501 | M | 20 | Valganciclovir hydrochloride | 0.3% | -2.3% | -1.3% | 8.8% | 175865-59-5 |
| Prestwick-P5 | 1502 | N | 20 | Sobuzoxane | -0.2% | 0.7% | -2.7% | -0.1% | 98631-95-9 |
| Prestwick-P5 | 1503 | O | 20 | Chicago sky blue 6B | -0.8% | -6.8% | -10.3% | 0.7% | 2610-05-1 |
| Prestwick-P5 | 1504 | P | 20 | Etiroxate hydrochloride | -4.0% | 7.8% | -7.9% | -18.2% | 55327-22-5 |
| Prestwick-P5 | 1505 | A | 21 | Barnidipine Hydrochloride | -3.9% | 4.6% | -11.2% | 6.5% | 104757-53-1 |
| Prestwick-P5 | 1506 | B | 21 | Benzylhydrochlorothiazide | -1.1% | -0.3% | 12.6% | 10.1% | 1824-50-6 |
| Prestwick-P5 | 1507 | C | 21 | Tacrolimus Monohydrate | 0.9% | 1.8% | -16.0% | 1.0% | 109581-93-3 |
| Prestwick-P5 | 1508 | D | 21 | Tandospirone | 0.4% | 0.2% | 2.3% | -5.0% | 87760-53-0 |
| Prestwick-P5 | 1509 | E | 21 | Lumiracoxib | 1.4% | -1.5% | -0.4% | -5.3% | 220991-20-8 |
| Prestwick-P5 | 1510 | F | 21 | Oxyphenisatin | -1.0% | 3.8% | -8.3% | 0.5% | 125-13-3 |
| Prestwick-P5 | 1511 | G | 21 | Thyrotropin-Releasing Hormone | -0.7% | 7.7% | -35.0% | 0.4% | 24305-27-9 |
| Prestwick-P5 | 1512 | H | 21 | Cyclofenil | 1.7% | 3.6% | -7.8% | -12.5% | 2624-43-3 |
| Prestwick-P5 | 1513 | I | 21 | Dilevalol | 0.7% | 1.5% | -15.8% | -9.2% | 75659-07-3 |
| Prestwick-P5 | 1514 | J | 21 | Repirinast | 1.5% | 1.1% | -4.7% | 2.3% | 73080-51-0 |

| | | | | | | | | | |
|---|---|---|---|---|---|---|---|---|---|
| Prestwick-P5 | 1515 | K | 21 | Rimonabant | 0.1% | 7.3% | -12.4% | 8.8% | 168273-06-1 |
| Prestwick-P5 | 1516 | L | 21 | Flupirtine | 0.9% | -0.5% | -4.6% | 9.3% | 56995-20-1 |
| Prestwick-P5 | 1517 | M | 21 | Cefozopran hydrochloride | -15.5% | 18.0% | -41.7% | -47.9% | 113981-44-5 |
| Prestwick-P5 | 1518 | N | 21 | Oseltamivir phosphate | 2.0% | -2.8% | -4.6% | 3.2% | 204255-11-8 |
| Prestwick-P5 | 1519 | O | 21 | Hippuric acid | 1.2% | -2.7% | -0.4% | 1.6% | 495-69-2 |
| Prestwick-P5 | 1520 | P | 21 | Riboflavine | 0.1% | -0.2% | 5.6% | -13.6% | 83-88-5 |
| Prestwick-P5 | co | A | 22 | DMSO | -4.0% | -1.1% | 5.0% | 20.5% | |
| Prestwick-P5 | co | B | 22 | DMSO | 1.3% | -3.0% | 14.6% | 7.5% | |
| Prestwick-P5 | co | C | 22 | Rott2 | -20.1% | 39.7% | -86.8% | -15.3% | |
| Prestwick-P5 | co | D | 22 | Rott2 | -22.0% | 46.1% | -87.6% | -24.1% | |
| Prestwick-P5 | co | E | 22 | Rott0.5 | -10.0% | 8.3% | -51.5% | -11.1% | |
| Prestwick-P5 | co | F | 22 | Rott0.5 | -10.9% | 10.9% | -52.0% | -13.2% | |
| Prestwick-P5 | co | G | 22 | Rott0.13 | -3.6% | 8.4% | -12.5% | -11.1% | |
| Prestwick-P5 | co | H | 22 | Rott0.13 | -4.0% | 4.2% | -10.8% | -3.0% | |
| Prestwick-P5 | co | I | 22 | DMSO | -0.5% | -0.8% | -5.2% | 11.1% | |
| Prestwick-P5 | co | J | 22 | DMSO | 2.1% | -1.4% | -0.2% | -0.4% | |
| Prestwick-P5 | co | K | 22 | DMSO | 3.2% | -1.9% | 4.3% | 10.4% | |
| Prestwick-P5 | co | L | 22 | DMSO | 2.1% | -1.1% | -7.6% | 8.5% | |
| Prestwick-P5 | co | M | 22 | DMSO | -0.2% | -2.4% | -3.5% | 3.4% | |
| Prestwick-P5 | co | N | 22 | DMSO | 0.4% | 1.4% | 8.8% | 1.2% | |
| Prestwick-P5 | co | O | 22 | DMSO | 2.5% | -1.0% | 1.6% | -3.8% | |
| Prestwick-P5 | co | P | 22 | DMSO | -2.5% | 0.4% | 11.4% | -3.1% | |
| Prestwick-P5 | co | A | 23 | DMSO | -1.4% | -3.2% | 5.3% | 15.3% | |
| Prestwick-P5 | co | B | 23 | DMSO | -1.7% | -1.2% | 2.9% | 7.8% | |
| Prestwick-P5 | co | C | 23 | DMSO | -0.3% | 0.0% | 0.8% | 0.1% | |
| Prestwick-P5 | co | D | 23 | DMSO | 0.4% | 1.5% | 6.4% | 3.9% | |
| Prestwick-P5 | co | E | 23 | DMSO | 1.1% | 5.6% | 2.1% | -2.3% | |
| Prestwick-P5 | co | F | 23 | DMSO | 0.3% | 5.0% | -0.1% | 7.6% | |
| Prestwick-P5 | co | G | 23 | DMSO | 0.1% | -2.5% | -5.0% | -8.9% | |
| Prestwick-P5 | co | H | 23 | DMSO | -2.4% | 6.8% | -16.8% | 3.5% | |
| Prestwick-P5 | co | I | 23 | DMSO | 1.0% | 2.7% | -0.1% | -1.4% | |
| Prestwick-P5 | co | J | 23 | DMSO | 2.1% | -1.2% | -2.5% | 10.0% | |
| Prestwick-P5 | co | K | 23 | DMSO | 1.7% | -2.8% | -5.0% | -1.1% | |
| Prestwick-P5 | co | L | 23 | DMSO | 2.2% | -0.2% | 7.5% | 4.6% | |
| Prestwick-P5 | co | M | 23 | DMSO | 1.9% | -2.4% | -4.8% | -1.8% | |
| Prestwick-P5 | co | N | 23 | DMSO | -0.4% | 2.2% | 20.5% | -3.8% | |
| Prestwick-P5 | co | O | 23 | DMSO | 0.4% | 1.0% | 4.3% | -23.0% | |
| Prestwick-P5 | co | P | 23 | DMSO | -1.5% | 0.5% | -4.7% | -6.8% | |
| Prestwick-P5 | co | A | 24 | noPFF | -3.9% | 1.7% | -99.8% | 12.3% | |
| Prestwick-P5 | co | B | 24 | noPFF | 0.5% | -3.1% | -99.7% | 3.8% | |
| Prestwick-P5 | co | C | 24 | noPFF | 1.9% | -1.2% | -100.0% | 3.8% | |
| Prestwick-P5 | co | D | 24 | noPFF | 1.8% | -0.7% | -100.0% | 5.9% | |
| Prestwick-P5 | co | E | 24 | Ceph4 | -28.6% | 45.2% | -100.0% | -57.1% | |
| Prestwick-P5 | co | F | 24 | noPFF | -1.8% | -4.7% | -100.0% | 10.6% | |
| Prestwick-P5 | co | G | 24 | noPFF | 0.9% | -2.6% | -100.0% | 4.1% | |
| Prestwick-P5 | co | H | 24 | noPFF | -2.5% | -0.3% | -100.0% | 5.9% | |
| Prestwick-P5 | co | I | 24 | noPFF | 1.7% | -2.3% | -100.0% | 8.9% | |
| Prestwick-P5 | co | J | 24 | noPFF | 2.8% | -3.4% | -100.0% | -0.5% | |
| Prestwick-P5 | co | K | 24 | noPFF | 3.6% | -4.0% | -99.8% | 12.5% | |
| Prestwick-P5 | co | L | 24 | noPFF | 2.2% | -3.2% | -100.0% | 8.6% | |
| Prestwick-P5 | co | M | 24 | noPFF | 1.7% | 1.8% | -99.7% | 3.7% | |
| Prestwick-P5 | co | N | 24 | noPFF | -0.1% | -2.9% | -100.0% | 1.1% | |
| Prestwick-P5 | co | O | 24 | noPFF | 0.3% | -3.3% | -100.0% | 4.9% | |
| Prestwick-P5 | co | P | 24 | noPFF | -1.4% | -2.8% | -100.0% | -6.9% | |

61-1

23-9

8-5
